# Extensive 5’-Surveillance Guards Against Non-Canonical NAD-Caps of Nuclear mRNAs in Yeast

**DOI:** 10.1101/2020.04.28.065920

**Authors:** Yaqing Zhang, David Kuster, Tobias Schmidt, Daniel Kirrmaier, Gabriele Nübel, David Ibberson, Vladimir Benes, Hans Hombauer, Michael Knop, Andres Jäschke

**Author notes:** Correspondence: Andres Jäschke.

## Abstract

The ubiquitous redox coenzyme nicotinamide adenine dinucleotide (NAD) acts as a non-canonical cap structure on prokaryotic and eukaryotic ribonucleic acids. Here we find that in budding yeast, NAD-RNAs are abundant (>1400 species), short (<170 nt), and mostly correspond to mRNA 5’-ends. The modification percentage is low (<5%). NAD is incorporated during the initiation step by RNA polymerase II, which uses distinct promoters with a YAAG core motif for this purpose. Most NAD-RNAs are 3’-truncated. At least three decapping enzymes, Rai1, Dxo1, and Npy1, guard against NAD-RNA at different cellular locations, targeting overlapping transcript populations. NAD-mRNAs do not support translation *in vitro*. Our work indicates that in budding yeast, most of the NAD incorporation into RNA seems to be accidental and undesirable to the cell, which has evolved a diverse surveillance machinery to prematurely terminate, decap and reject NAD-RNAs.

**In Brief:** In budding yeast, most of the NAD incorporation into RNA seems to be accidental and undesirable to the cell, which has evolved a diverse surveillance machinery to prematurely terminate, decap and reject NAD-RNAs.

**Graphical Abstract:** 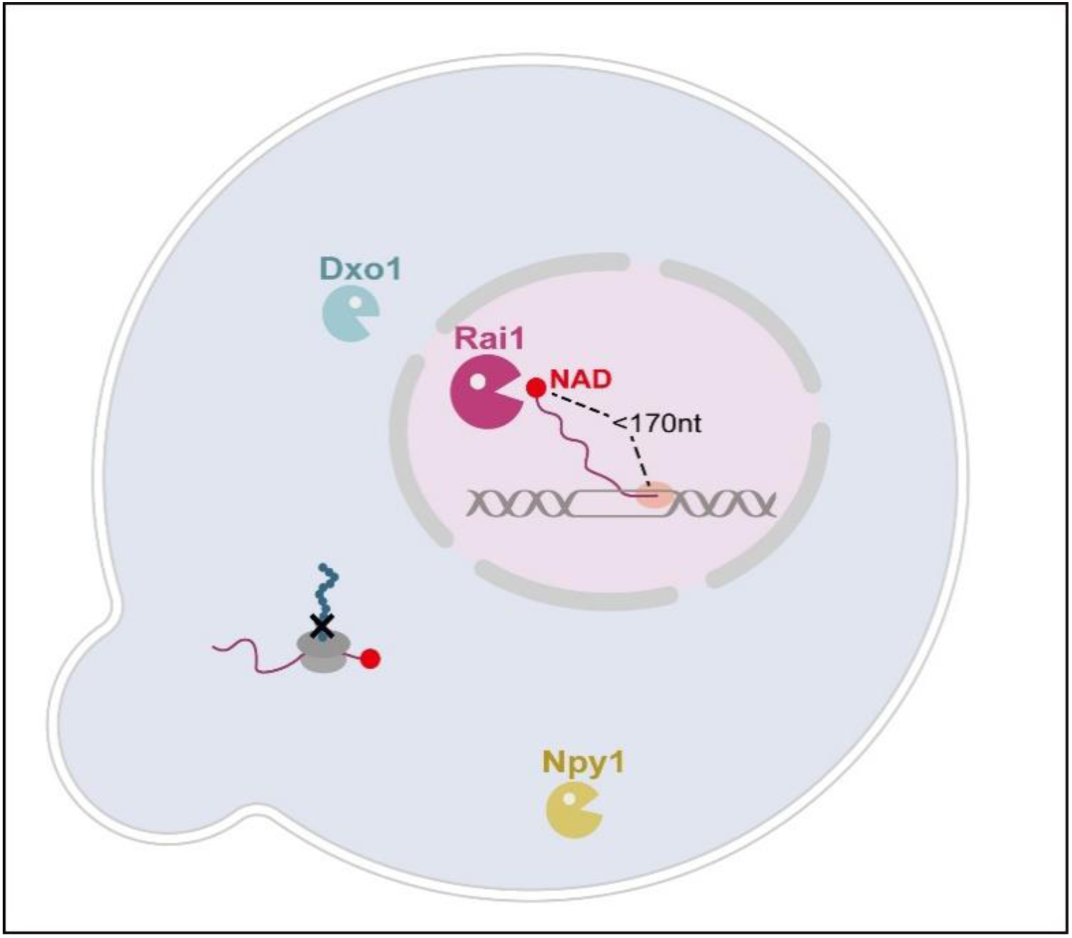

**Highlights:**

- Yeast cells have thousands of short NAD-RNAs related to the 5’-ends of mRNAs
- RNA polymerase II prefers a Y<u>A</u>AG promoter motif for NAD incorporation into RNA
- NAD-RNA is strongly guarded against by Rai1, Dxo1, and Npy1 decapping enzymes at different subcellular sites
- *In vitro*, NAD-mRNAs are rejected from translation

## Introduction

In eukaryotes, the 5’-terminus of messenger RNAs is protected by a m^7^-guanosine (m^7^G) cap (Shatkin, 1976), which modulates pre-mRNA splicing, polyadenylation, nuclear exit, and translation initiation (Topisirovic et al., 2011). The cap is hydrolyzed by various decapping enzymes (Belasco, 2010), thereby triggering RNA degradation. Recently, another type of 5’-cap structure was discovered, both in prokaryotes (Cahova et al., 2015; Frindert et al., 2018; Morales-Filloy et al., 2020) and eukaryotes (Jiao et al., 2017; Walters et al., 2017; Wang et al., 2019; Zhang et al., 2019), that is derived from the ubiquitous redox coenzyme NAD. While in human and plant cells a diverse landscape of NAD-RNAs was found, only 37 RNA species and low abundance were reported in budding yeast (Walters et al., 2017), questioning the biological significance of NAD capping in this organism. As the protocol used in this work excluded the small-RNA fraction, which had been particularly rich in NAD-RNAs in prokaryotes (Cahova et al., 2015), we address here the whole landscape of NAD transcripts in yeast using the original NAD captureSeq protocol (Winz et al., 2017). We find that NAD-RNAs are ubiquitous (1400 in wild-type, several thousands in mutants), most of them being short species (<170 nt). Only very few RNAs are detected with lengths over 250 nt. *Ab initio* incorporation by RNA polymerase (RNAP) II is found to be the predominant mechanism for NAD incorporation, and for about half of the transcripts, RNAP II uses transcription start sites different from those utilized for m^7^G-capped RNAs of the same gene. A YAAG core promoter motif is found to correlate with efficient transcriptional NAD incorporation. On average, NAD-RNAs are shorter than non-NAD-RNAs. By deleting the (putative) NAD-RNA decapping enzymes Npy1, Dxo1, and Rai1 we identified overlapping populations of RNAs decapped by these enzymes. Sequence analysis of total RNA from each mutant strain supports a hierarchical order of NAD-RNA processing, in agreement with the subcellular locations of the enzymes. NAD-mRNAs that escape decapping do not support translation by cytosolic ribosomes *in vitro*. We propose that (at least for the nuclear transcripts studied here) NAD incorporation into yeast RNA is largely accidental, due to competition of NAD and ATP in transcription initiation. We speculate that the NAD modification is in most cases undesirable to the cell, which first disfavors the synthesis of full-length NAD-RNAs, then decaps them rapidly using a multi-tiered machinery localized in different compartments, and – even if they reach full length and escape decapping – ultimately rejects them from ribosomes.

## RESULTS

### Short NAD-RNAs are abundant in yeast

To comprehensively address NAD-RNAs in budding yeast, we isolated total RNA from yeast strain BY4742 and applied the original NAD captureSeq protocol (Cahova et al., 2015), in which the enzyme ADP ribosyl cyclase (ADPRC) tags NAD-RNAs at the NAD moiety, followed by “click”-chemistry biotinylation and selective isolation by streptavidin binding. Enrichment was determined by quantitative comparison with a minus ADPRC negative control. After PCR amplification, amplicons were size-selected, thus this library represented mostly RNA species with sizes between 20 and 170 nt present in the original sample. In this unfragmented library, 1460 RNAs were found to be enriched, with changes reaching up to 1200-fold (**Figure 1A**). 69% of the genome-mapped reads corresponded to mRNA 5’-ends (**Figure 1B**), while only little enrichment was observed for mRNA fragments starting further downstream (for details and validation, see Supplementary Methods and **Figures S1A-D**). Small nucleolar RNAs (snoRNAs) and ribosomal RNA (rRNA) fragments comprised 9.6 and 7.1% of the reads, but represented only 2 and 1 different RNA species, respectively. 13% corresponded to RNA fragments too small for unique genome mapping (12 – 17 nt, **Figure 1B**), many of which showed homology to enriched members of the mRNA 5’-end group (**Figure S1E**). To probe the existence of full-length NAD-capped mRNAs, two additional datasets were generated from total RNA that was random-sheared prior to NAD captureSeq using different size selection windows (“small fragmented”: 20 – 170 nt; “large fragmented”: 170 – 350 nt). Consistently, these fragmented libraries revealed much lower numbers of enriched species and enrichment values (small fragmented: 145 RNAs, maximum fold change <7, **Figure 1C**, large fragmented: 200 RNAs, maximum fold change <9, **Figure S1F**), mainly due to increased background in the minus ADPRC controls. Extensive overlap was detected between the two fragmented libraries and the unfragmented one (83.8% and 76.0%, respectively, **Figure S1G**). Five out of the 12 genes explicitly reported in the previous study ((Walters et al., 2017), **Figure S1H**) overlap with the enriched species in our unfragmented library (COX2; LSM6; ERG2; UBC7; YJR112W-A) and two with the fragmented library (LSM6; UBC7, black dots in **Figures 1A and 1C**).

**Figure 1.**
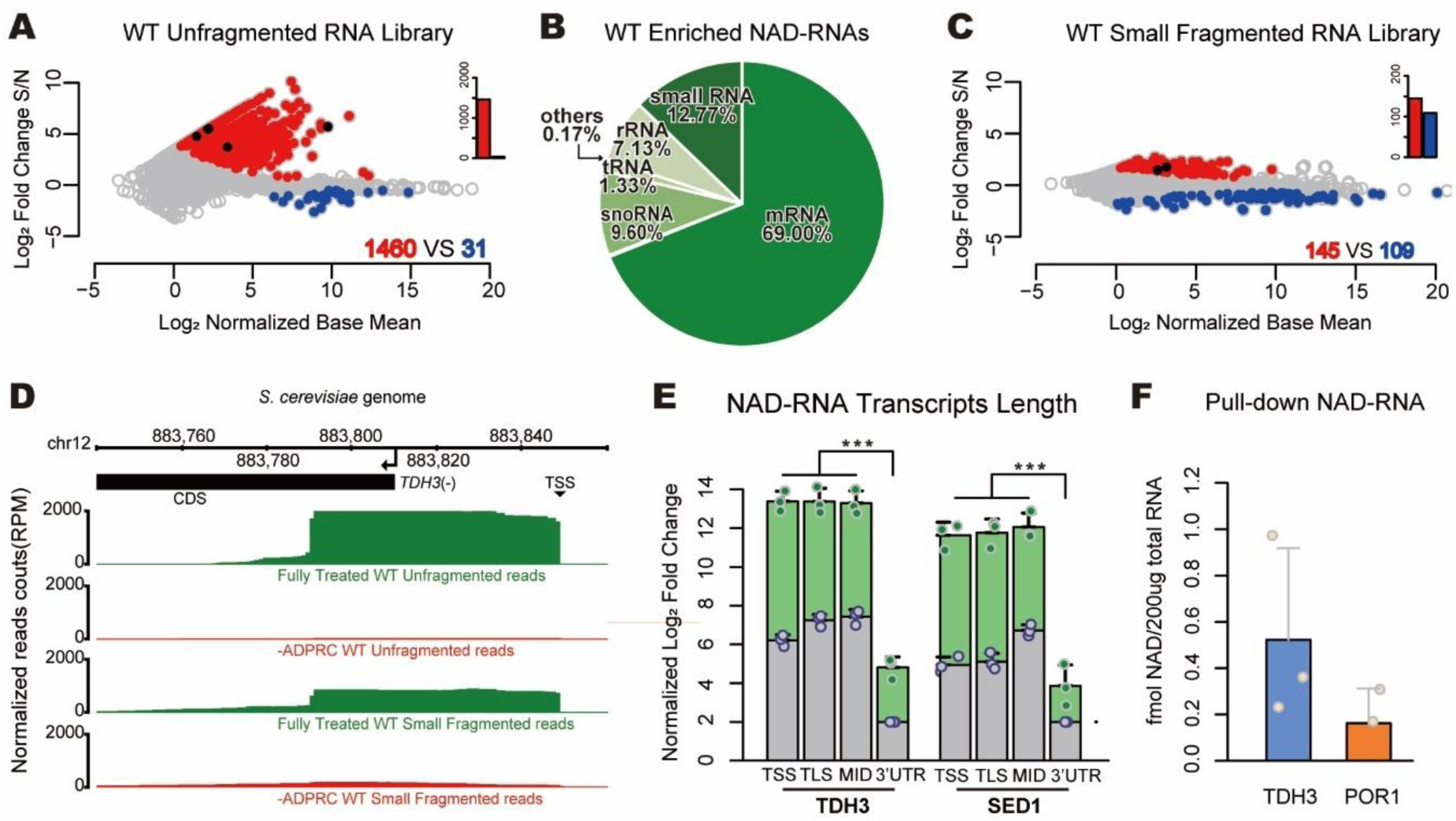
NAD-RNAs are abundant in yeast *S. cerevisiae.* (A) Enriched NAD-RNAs from unfragmented NAD captureSeq on the wild type (WT) strain. The log_2_ fold change between fully treated sample (S) and minus ADPRC negative control (N) is plotted versus the log_2_-normalized base mean. 7620 different transcripts are analyzed and represented as a dot. Red dots represent enriched NAD-transcripts (fold change (FC) >1.414, normalized base mean (NBM) >1, p <0.05), blue dots are negatively enriched transcripts (FC <0.707, NBM >1, p <0.05), black dots are transcripts previously reported as NAD-capped (Walters et al. 2017). Biological triplicates. (B) Reads of enriched NAD-RNA from WT unfragmented NAD captureSeq library, mapped to genome annotated RNAs. Chart contains reads classified as genome annotated RNAs (reads ≥18nt, same statistics parameters as in panel A), plus the self-categorized small RNAs (12nt ≤ reads <18nt, fold change S/N >1.414). Other categories included pseudogene (0.093%), ncRNA (0.057%), and long-term repeat (0.017%). (C) Enriched NAD-RNAs from small fragmented NAD captureSeq on WT strain. Random-sheared total RNA as input and sized selected cDNA (insert <172bp) was utilized for library building. Statistics parameters are as in panel A. Biological triplicates. (D) Aligned sequencing reads of *TDH3* gene were visualized in the integrated genome browser (IGB). Reads count was normalized as per million genome mapped reads (RPM). The arrow indicates the transcription direction, and the coding sequence is displayed as black bar. (E) qRT-PCR of NAD-RNAs targeting different regions of the gene: the 5’-end including the transcription start site (TSS), the region around the translation start site (TLS), a middle (MID) region around the first in-frame ATG after the TLS, and the 3’ UTR. Green bar heights represent relative transcript numbers of S while grey bar heights represent the same for N. Log_2_ fold change of TLS, TSS and MID was normalized to the 3’ UTR of N (set as 2). Dots represent the measured values. Error bars represent mean + standard deviation (SD), n=3. p values are denoted by asterisks: (*) p <0.05; (**) p <0.01; (***) p <0.001(the minimum value among TSS, TLS, MID versus the value from the 3’ UTR, Student’s t test). (F) Pulled-down NAD-RNA quantified by LC-MS. NAD-RNAs were quantified by the NAD content which was determined via the signal intensity of nicotinamide riboside (NR) (fmol) from NudC treated pull-downed RNA minus the RNA without NudC treatment. Statistics parameters are as in Figure 1E.

To test whether highly expressed transcripts are generally more likely to be enriched in NAD captureSeq, we compared the enrichment levels observed in the unfragmented NAD captureSeq library with the transcript abundance determined by transcriptome sequencing. This analysis revealed no correlation (**Figure S1I**).

As NAD captureSeq enriches the 5’-ends of NAD-RNAs and sequences in 5’ to 3’-direction, the overall lengths of transcripts larger than the Illumina read length cannot be reliably inferred from the sequencing reads (**Figure 1D**). We therefore carried out the first steps of the NAD captureSeq protocol (until the enriched RNAs were bound to streptavidin) and carried out RT-qPCR on two RNA species that were enriched in the fragmented and unfragmented libraries (TDH3 and SED1) using four different primer pairs each. These data revealed roughly equal abundance from the 5’-end through ∼300 nt, while their 3’ UTRs were reduced in abundance by several orders of magnitude (**Figure 1E**, green bars).

For two transcripts (TDH3 and POR1 mRNA), the 5’-NAD modification was directly identified and quantified by mass spectrometry after pull-down (**Figure 1F**), confirming the chemical identity of the NAD modification. Thus, NAD-RNAs are abundant, short, and mostly correspond to mRNA 5’-ends in budding yeast.

### Nudix pyrophosphohydrolase Npy1 processes NAD-RNA *in vitro* and *in vivo*

In *E. coli*, the Nudix hydrolase NudC acts as an efficient decapping enzyme for NAD-RNA (Cahova et al., 2015; Höfer et al., 2016; Zhang et al., 2016). The yeast homolog Npy1 is known to hydrolyze the pyrophosphate bond in NAD to yield nicotinamide mononucleotide (NMN) and adenosine monophosphate (AMP) (Xu et al., 2000) and was recently suggested as an NAD-RNA decapping enzyme (Zhang et al., 2016). The only support for this claim was, however, its *in vitro* processing of a synthetic NAD-RNA 12mer into a product that migrated on HPLC like a 12mer-5’-monophosphate RNA (p-RNA), and the inactivity of an active-site mutant to produce this product (Zhang et al., 2016). To characterize the *in vitro* activity of Npy1, we purified the protein from *E. coli* and analyzed its reaction kinetics with an *in vitro* transcribed NAD-RNA (a 98 nt 5’-fragment of TDH3 RNA) on acryloylaminophenyl boronic acid (APB) gels which separate NAD-RNA from p-RNA (Nübel et al., 2017). Purified Npy1 decapped NAD-RNA without inducing nucleolytic degradation and had no effect on m^7^G-RNA *in vitro* (**Figure 2A**). Furthermore, efficient decapping of NAD-RNA required Mn^2+^ ions (**Figures S2A-C**). A Npy1 mutant in which a catalytic glutamate was replaced (E276Q) showed no decapping activity (**Figure S2D**). In addition to NAD-RNA, Npy1 also hydrolyzed NAD into NMN and AMP in a Mn^2+^-dependent manner (**Figure S2E**), while the E267Q mutant was inactive (**Figure S2F**). Thus, recombinant Npy1 decaps NAD-RNA *in vitro*.

**Figure 2.**
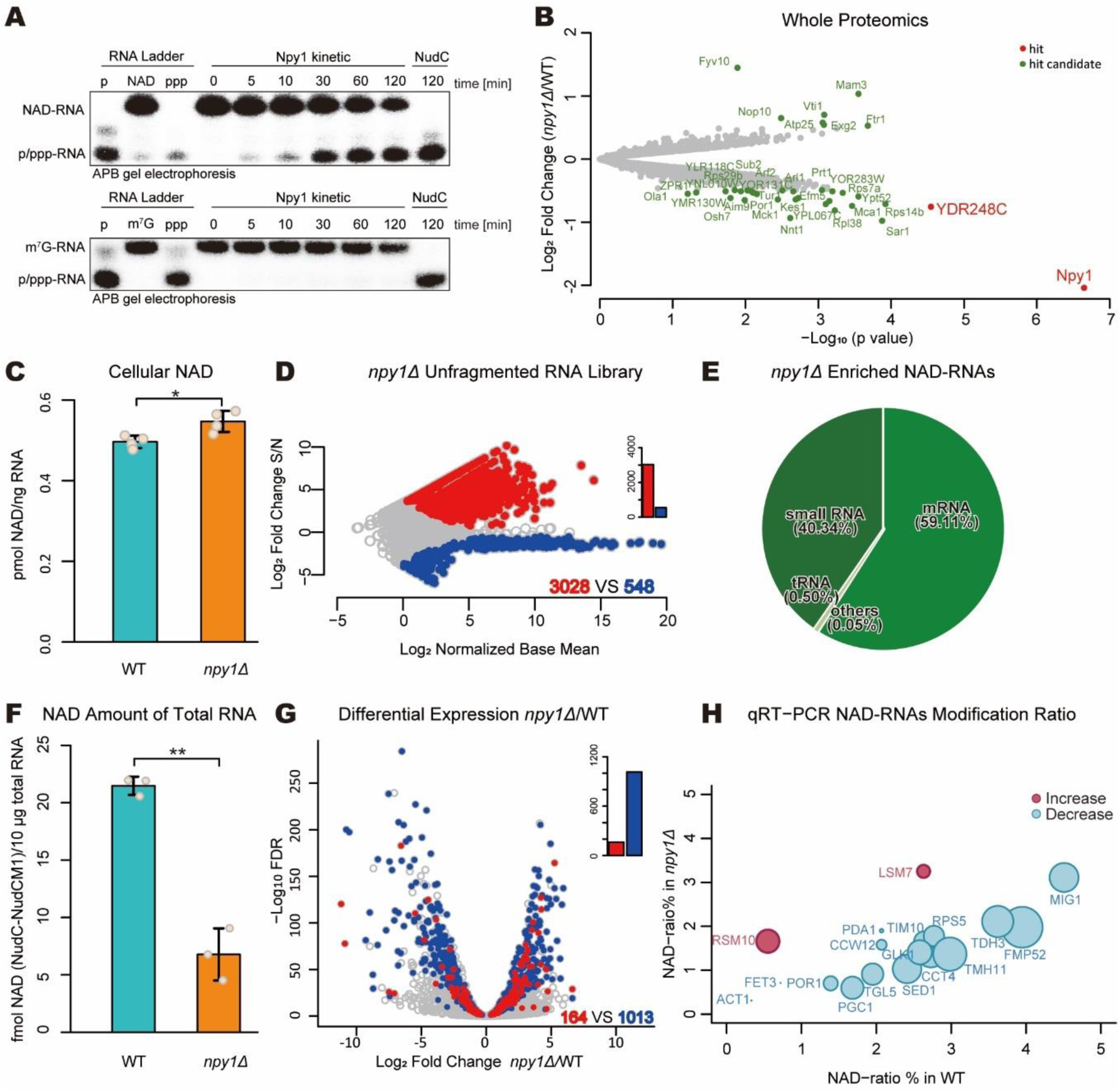
Npy1 affects NAD-RNA *in vitro* and *in vivo*. (A) Npy1 WT kinetics of decapping NAD- and m^7^G-capped RNA with Mn^2+^*in vitro*. A α-^32^P body-labeled 5’-fragment of TDH3 RNA (98nt) and a corresponding ppp-RNA control were treated with Npy1 in the presence of 2 mM Mg^2+^ and 1 mM Mn^2+^ and separated by APB gel electrophoresis. (B) Whole proteome analysis of cell lysates (WT strain *vs. npy1Δ*). Red dots indicate enriched hits (fold change >1.5 and FDR <0.05) while green dots are hit candidates (fold change >1.4, 0.05 ≤ FDR < 0.2) and grey dots are proteins without significant change. (C) Quantification of cellular NAD in WT and *npy1Δ* strain by enzyme cycling assay. The amount of NAD was normalized to the amount of total RNA determined in cell lysate from each sample. Error bars represent mean ± sd, n=4. p values are denoted by asterisks: (*) p <0.05 (Student’s t test). (D) Enriched NAD-RNAs from unfragmented NAD captureSeq of the Npy1 depletion (*npy1Δ*) strain. Biological triplicates. (E) Reads of enriched NAD-RNA from *npy1Δ* unfragmented NAD captureSeq Library, mapped to genomically annotated RNAs. All parameters as in Figure 1B. (F) Quantification of NAD-RNA from total RNA by LC-MS. Integrated NAD content which was determined via the NR signal intensity (fmol) from NudC-treated washed total RNA (10 µg) minus the RNA treated with a NudC mutant. Dots represent individual biological triplicate measurements. Error bars represent mean ± sd, n=3. p values are denoted by asterisks: (**) p <0.01 (Student’s t test). (G) Volcano plot of transcripts comparing WT with *npy1Δ* samples by RNA expression level and NAD modification ratio. The log_2_ fold change of transcript abundance from transcriptome sequencing (*npy1Δ*/WT) is plotted versus the log_10_ false discovery rate (FDR). 7620 different transcripts are analyzed and represented as dots. Red dots represent RNAs for which the NAD-ratio increased upon *NPY1* gene deletion according to NAD captureSeq (NAD-ratio: *npy1Δ* > WT > 0, transcriptome normalized base mean >100, p< 0.05, FDR <0.1), and blue dots represent RNAs for which the NAD-ratio decreased (NAD-ratio: WT > *npy1Δ*> 0, transcriptome normalized base mean >100, p <0.05, FDR <0.1). (H) Bubble plot of the relative NAD modification of 18 RNA species by qRT-PCR. Blue bubbles represent RNAs for which the NAD modification ratio decreased upon *NPY1* gene deletion, while red bubbles show those with increased NAD modification. The bubble size indicates the extent of the relative change.

To address whether Npy1 also functions on NAD-RNA *in vivo*, we investigated a yeast strain lacking Npy1. In agreement with the yeast SGA database (Baryshnikova et al., 2010), the absence of Npy1 caused no severe phenotypical changes under a variety of growth conditions (**Figure S2G**). While gene expression analysis by transcriptome sequencing indicated changes in abundance for almost 50% of all detected transcripts (**Figure S2H**), mass-spectrometric whole proteome analysis detected only very few proteins with significant (more than two-fold) changes in expression, in comparison to the WT strain (**Figure 2B**). Deletion of *npy1* slightly increased the total cellular concentration of NAD (by ∼10%, **Figure 2C**). When we applied NAD captureSeq to RNA purified from the *npy1Δ* strain, twice as many uniquely mapped RNAs (3028, unfragmented library) were NAD-capped (relative to WT), which were almost half of all detected RNA species (**Figure 2D**). Consistent with the WT, NAD-RNAs from the *npy1Δ* strain were mostly short transcripts, as only 242 and 220 NAD-RNA species were enriched in the small and large fragmented RNA libraries, respectively (**Figures S2I and S2J**). Compared to the WT, a similar proportion of the reads allocated to mRNA 5’-ends in the unfragmented library (59.1%) but three times more on very small RNAs (40.3%), suggesting that Npy1 is involved in the decapping of small NAD-RNAs. rRNAs and snoRNAs disappeared almost completely (**Figure 2E**).

Contrary to our expectations (but as also observed in *B. subtilis* (Frindert et al., 2018)), removal of Npy1 reduced the total amount of NAD attached to RNA by ∼60% (**Figure 2F**). We assessed the change in the apparent modification ratio (percentage of an RNA species that carries NAD) transcriptome-wide by integrating transcriptome and NAD captureSeq data, using the enrichment values in NAD captureSeq as proxy (**Figure 2G**, for details and method validation, see Supporting Information). This analysis indicated that upon *NPY1* gene deletion, the NAD modification ratio was reduced for 1013 species, while it increased for 164. There was no correlation between expression level (change) and modification ratio (change). Plotting the modification ratio of *npy1Δ* mutant vs. WT confirmed that the global reduction of NAD modification is not caused by few strongly reduced species that override the effects of many weakly increased ones. The slope < 1 (0.61) of this plot confirms that, on average, the modification in the *npy1Δ* mutant is lower than in the WT (**Figure S2K**). To independently support the decreased modification ratios in the npy1Δ mutant derived from NAD captureSeq, we quantified 18 different RNAs by qRT-PCR in the unfragmented cDNA libraries of sample (S) and negative control (N) for WT and mutant strain. After normalization of the cp values to the same amount of input RNA in WT and npy1Δ and background subtraction, those 16 genes with decreased NAD modification showed indeed reduced PCR amplification, while those two with increased modification PCR-amplified stronger (**Figure 2H**). Collectively, these data indicate that Npy1 decaps NAD-RNAs *in vitro* and *in vivo*.

### Deletion of Npy1, Rai1, and Dxo1 influences the NAD-RNA landscape

The non-Nudix enzymes Rai1 and Dxo1 were previously reported to decap NAD-RNA *in vitro* and *in vivo* by a mechanism different from Npy1, namely by removal of the entire NAD moiety *en bloc* (Grudzien-Nogalska et al., 2018; Jiao et al., 2017). To compare the influence of all three enzymes on the global NAD-modification landscape of RNAs *in vivo*, we created all possible combinations of *rai1Δ, dxo1Δ*, and *npy1Δ* deletion mutants. Phenotypically, the removal of Rai1 (from the WT and from mutant strains) had the strongest negative effect on growth in normal medium and in the presence of increasing concentrations of ethanol (**Figures 3A and S3A**). On the transcriptome level, we detected in all mutant strains ∼1000 upregulated and ∼1000 downregulated RNA species (at least four-fold, relative to WT), together corresponding to ∼30% of all mRNAs (**Figure S3B**). There was over 60% overlap in regulated genes between the three different single-knockout mutants, while ∼600 genes were selectively regulated by only one decapping enzyme (**Figure S3C**). A systematic analysis of the effect of the deletion of one particular enzyme in WT and mutant strains revealed high agreement within one group (e.g., all strains carrying a deletion of the *NPY1* gene), and the strongest global effect on RNA expression was noticed for removal of *RAI1* (**Figure 3B**). Analysis of total RNA isolated from the knockout mutants by NAD captureSeq revealed enrichment of more than half of all detected RNA species (3765 in *dxo1Δ*; 3810 in *rai1Δ*), indicative of their modification with NAD. No significant further increase was observed in the double- and triple-deletion strains (**Figure 3C**). In the triple knockout *dxo1Δ rai1Δ npy1Δ*, only mRNA fragments (63%) and small RNAs (35%) were detected by NAD captureSeq (**Figure 3D**). Unlike the WT, the top 250 enriched NAD-RNA species of all mutants functionally clustered (by GO terms) as rRNA metabolic process and translation (**Figure S3E**). Thus, all three enzymes act on NAD-RNA *in vivo*.

**Figure 3.**
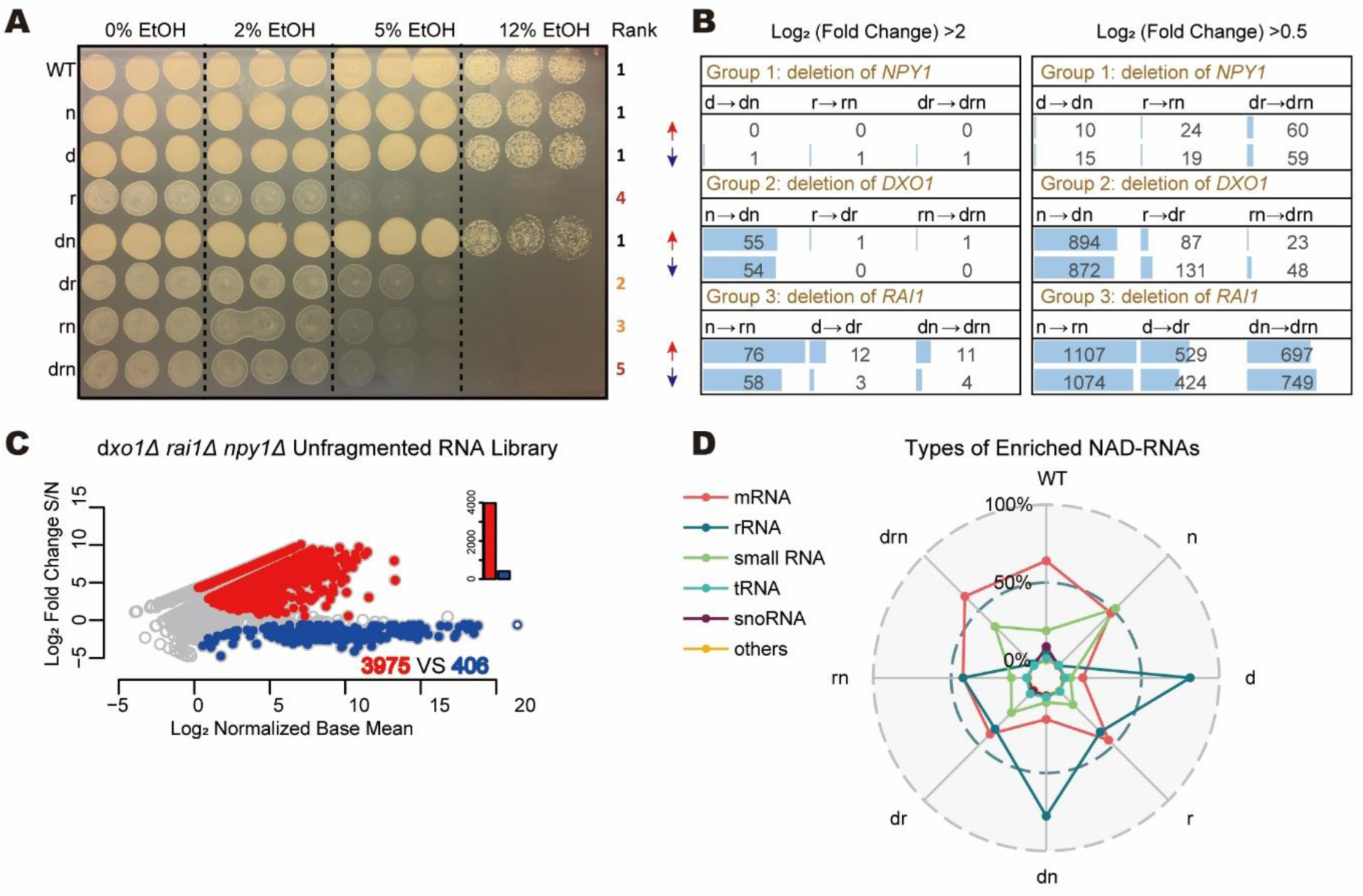
Deletion of decapping enzymes influences the NAD-RNA landscape. (A) Growth phenotype of the WT and mutant strains In the presence of different ethanol concentrations. The rank of approximate growth density is given on the right side. Each condition was tested in triplicates. Strain abbreviations (left) are *npy1Δ*(n), *dxo1Δ*(d), *rai1Δ*(r), *dxo1Δ npy1Δ* (dn), *dxo1Δ rai1Δ* (dr), *rai1Δ npy1Δ (rn), dxo1Δ rai1Δ npy1Δ* (drn). (B) Effect of the deletion of one particular enzyme in different strains on the up-regulation (red arrows) or down-regulation (blue arrows) of RNA species, assessed by transcriptome sequencing. Strain abbreviations as in Figure 3A. Group 1 summarizes the deletion of *NPY1* gene from the single mutants (d, r) and from the double mutant (dr). Groups 2 and 3 show deletion of *DXO1* gene and of *RAI1* gene, respectively. The number of RNA species with log_2_FC >2 is on the left panel while with log_2_FC >0.5 is on the right panel. (C) Enriched NAD-RNAs from unfragmented NAD captureSeq on the *dxo1Δ rai1Δ npy1Δ* strain. All parameters are as in Figure 1A. Biological triplicates. (D) Radar plot of the distribution of different classes of enriched NAD-RNAs (in %) in different deletion strains. The colors indicate the type of RNAs. Strain abbreviations as in the Figure 3A.

### Transcription start sites for NAD-RNAs differ from those for m^7^G-RNAs

The above analysis suggested that the landscape of NAD-RNA transcripts is shaped by (at least) four enzymes: RNAP II, Rai1, Dxo1, and Npy1. Using the deletion mutants, we first analyzed transcriptional preferences. While the sequencing read profiles of some RNAs revealed homogenous 5’-ends (indicative of a defined transcription start site, TSS), others showed irregular patterns suggesting pervasive transcription or multiple TSSs (See **Figures 1D and S4A** for examples). From the NAD captureSeq data we selected all significantly enriched RNAs starting with an A which had homogenous 5’-ends (‘sharpA’ selection). When we compared our experimentally determined 5’-ends of these NAD-RNAs with published next generation sequencing (NGS)-derived and 5’-rapid amplification of cDNA ends (RACE)-validated TSSs for canonical (i.e., non-NAD-) RNAs (Nagalakshmi et al., 2008), for nearly half of all species the 5’-ends differed (**Figures 4A and 4B**). For the WT strain, 98 RNAs were observed in which the 5’-transcript leader (TL) sequences were longer than in the database, while 63 species got shorter, in some cases by more than 100 nt (**Figure 4C**). This TL length change was not only observed in the WT strain (in both unfragmented and fragmented libraries, **Figures 4C and S4B-C**), but also in all mutants, including the dxo1Δrai1Δnpy1Δ triple mutant (**Figures 4B and S4D**), suggesting that RNAP II might select a different TSS for initiating transcription with NAD instead of ATP, compared to the canonical TSS (Gilbert et al., 2007; Rojas-Duran and Gilbert, 2012). The changed TL length upon NAD incorporation could be corroborated by qRT-PCR with primers targeting either our NAD captureSeq-observed TSS or the canonical ones from the database, comparing the RNAs enriched in NAD captureSeq with a non-enriched total RNA preparation (**Figure 4D**). Thus, NAD-RNAs tend to have longer TL sequences than non-NAD-RNAs, indicative of their synthesis starting at a more distal TSS.

**Figure 4.**
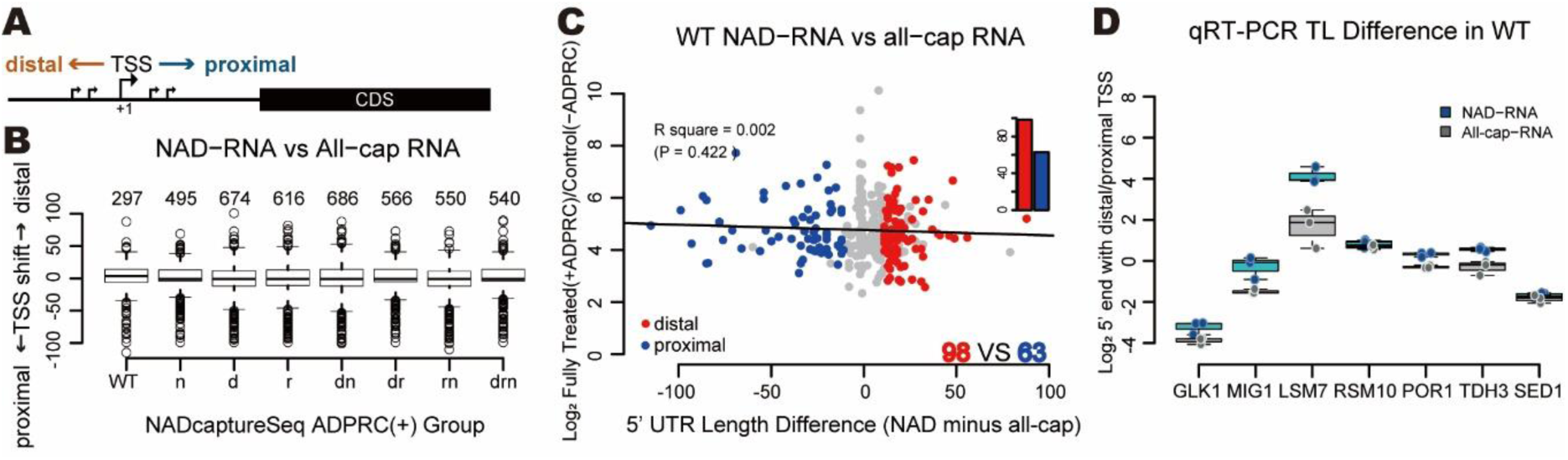
NAD-RNAs tend to have longer 5’ UTRs than non-NAD-RNAs. (A) Scheme of TSS shifting in proximal or distal direction. (B) Global TSS shifting between NAD-RNA (according to NAD captureSeq, unfragmented libraries) in all strains and canonical RNA (according to Nagalakshmi dataset (Nagalakshmi et al., 2008)). The numbers above each box group indicates the number of RNA species analyzed in this strain. Strain abbreviations are as in Figure 3A. (C) Detailed TSS shifting between NAD-RNA and All-cap RNA in WT strain, 5’ UTR length difference is NAD-RNA minus canonical (all-cap) RNA. Red dots represent NAD-RNA species with distal TSS (>10 nt difference) and blue dots represent with proximal TSS (>10 nt difference). FDR_wilcox_ <0.1. (D) qRT-PCR with different primer pairs confirming different transcript leader (TL) lengths between NAD-capped RNAs (enriched fraction from NAD captureSeq, blue boxes) and all-capped RNAs (from non-enriched input RNA, grey boxes). The box plot displays the log_2_ of the ratio of the number of transcripts derived from the distal TSS to that from the proximal TSS. Dots represent individual biological triplicate measurements, and error bars represent corresponding standard deviation.

### Efficient *in vivo* NAD incorporation by RNAP II is supported by a YAAG core promoter motif

We supposed that analysis of the *dxo1Δ rai1Δ npy1Δ* triple knockout mutant strain would reveal the least biased information about the factors that govern transcriptional NAD incorporation by RNAP II. We mapped nucleotides -10 to +10, relative to the RNA 5’-end inferred from the NAD captureSeq reads, for the 25 most enriched ‘sharpA’ NAD-RNAs (log_2_FC >8, FDR <0.00002), and for appropriate control groups (i.e., RNA species not enriched in NAD captureSeq). In the enriched fraction we observed a highly conserved motif Y<u>A</u>AG (with the first A being the 5’-terminal nucleotide of the transcript, i.e., the site where the NAD is incorporated), followed by an A-rich stretch of lower significance, while in the non-enriched fraction no preferences were found (**Figure 5A**). The motif was not observed when for the same top 25 candidate RNAs the published canonical TSSs (Nagalakshmi et al., 2008) were mapped (**Figure 5B**). When for those 25 genes all TSSs listed in the yeast TSS database (McMillan et al., 2019) (top 5 abundant TSSs per gene) were analyzed, only a Y<u>A</u> motif (Zhang and Dietrich, 2005) was identified (**Figure S4A**). However, when only the TSS (from this database) closest to our observed one was utilized, the Y<u>A</u>AG motif appeared prominently (**Figure S4B**). This analysis implies that our identification procedure revealed real TSSs. The Y<u>A</u>AG motif was also observed (although less prominently) in the top 100 and 200 enriched RNAs, and its prominence decreased with decreasing NAD captureSeq enrichment values (**Figures S5C and S5D**). It could also be detected in WT and all mutant strains, whereby generally the significance decreased with increasing number of decapping enzymes present (**Figures S5E-K**). The motif was not observed when the NAD captureSeq-enriched snoRNAs or transfer RNAs (tRNAs) were mapped (**Figures S5L and S5M**), suggesting that these candidates may have a different biogenesis. To exclude the possibility that the motif reflects a bias introduced by the enzymes applied in NAD captureSeq (ADPRC, reverse transcriptase, terminal deoxynucleotidyl transferase, two ligases), we mapped the top 25 enriched sequences from our previously published *E. coli, B. subtilis* and *S. aureus* NAD captureSeq datasets by the same procedure, finding neither Y<u>A</u>AG nor an A-rich tail (**Figures S5N-P**). Further analysis revealed that this motif constitutes a fraction of known ‘good’ RNAP II core promoter sequences, having all conserved features (Hahn et al., 1985; Lubliner et al., 2013; Maicas and Friesen, 1990), namely: 1. being A/T rich between positions -30 and +10, 2. a switch from T-rich to A-rich in the coding strand around position -8, 3. a pyrimidine at position -1, and 4. an A at position +1. In addition, two specific features distinguish good NAD-incorporating promoters, namely a slightly increased probability for an A at position +2, and a strongly conserved G at +3.

**Figure 5.**
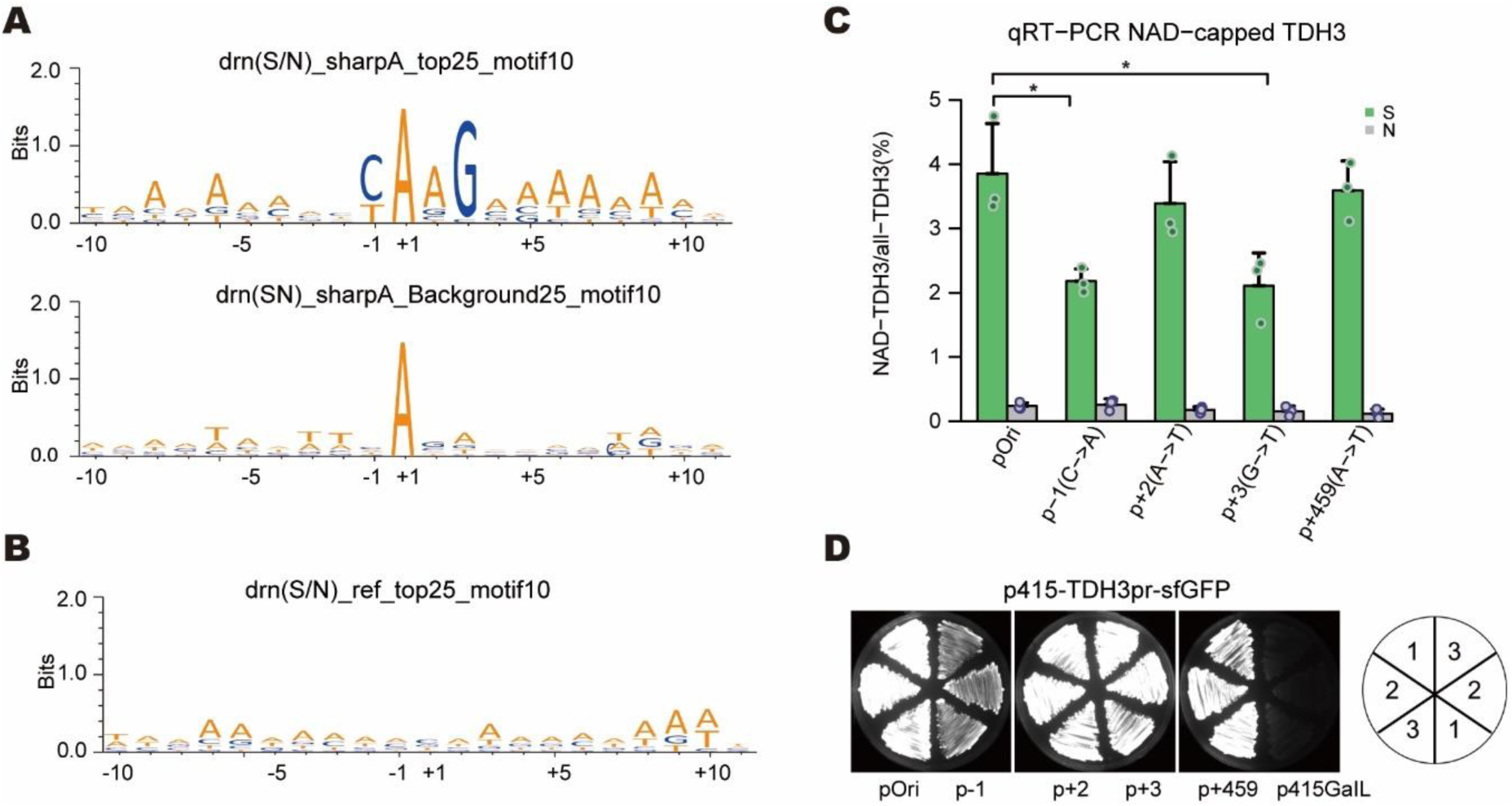
NAD is incorporated by RNAP II in a promoter-dependent manner. (A) Motif analysis of the -10 to +10 region around the TSS based on NAD captureSeq data. Top25 and background25 represent the 25 NAD-mRNA species with the highest enrichment values and the 25 most abundant, but not significantly enriched mRNA species (0.707⩽S/N⩽1.414) with a “sharpA” feature at position +1(TSS) in the *dxo1Δ rai1Δ npy1Δ* (drn) strain. S/N is the enrichment of NAD-RNA in the sample group (+ADPRC) compared to the negative control group (-ADPRC). “SharpA” means an ‘A’ at position +1 with a more than three-fold higher signal than that of the nucleotide in the -1 position. (B) Motif analysis of the same Top25 enriched RNA species, using the canonical TSS from Nagalakshmi dataset. (C) Quantification of the NAD-modification ratio using the *TDH3* gene promoter and relevant mutants *in vivo*. The height of the green bar indicates the TDH3 RNA NAD-ratio in the sample group (+ADPRC) while the grey bar indicates that in N group (-ADPRC). The pOri represents the original TDH3 promoter, and promoter mutants are indicated below the bar chart. Dots are biological triplicates and error bars represent standard deviations. p values are denoted by asterisks: (*) p <0.05 (Student’s t test). (D) Expression of sfGFP under a *TDH3* gene promoter or relevant promoter mutants *in vivo*. pOri, p-1, p+2, p+3, and p+459 are same in Figure 5C. p415GalL is negative control strain for background. All strains were cultured as biological triplicates.

To test whether this motif actually modulates NAD incorporation by RNAP II *in vivo*, we deleted gene *tdh3*, a highly enriched NAD-RNA observed in every strain, and added a low-copy plasmid in which we inserted a DNA fragment containing the 600 bp upstream of the *TDH3* gene, containing the entire promoter region, plus the first 54 nt after the experimentally observed TSS of the TDH3 RNA (39 nt 5’-UTR, 15 nt coding sequence), followed by the ORF of *superfold-GFP* to monitor gene expression (**Figure S5Q**). Mutants were prepared in which the Y at position -1, the A at position +2 and the G at position +3 were individually varied. An additional mutant was generated in which all A’s in the tail region (+4, +5 and +9) were replaced. Cells were transfected with these plasmids, and harvested around OD_600_ = 0.8. Total RNA was isolated, treated with ADPRC, followed by click biotinylation, streptavidin purification, and reverse transcription. qPCR with gene-specific primers was used to assess the percentage of NAD-modified TDH3 RNA in each strain, using pure synthetic spike-in NAD-RNA and ppp-RNA to ensure equal reactivity of each sample. This analysis revealed indeed strong (∼2-fold) reduction of relative NAD incorporation upon mutating positions -1 and +3, while for positions +2 and the A-rich tail the observed effects were not statistically significant (**Figures 5C and S5R**). Quantification of GFP expression levels revealed that mutating position -1 significantly decreases both NAD-RNA and non-NAD-RNA, while mutating position +3 modulates exclusively NAD-RNA (**Figures 5D and S5S**). Thus, a specific promoter sequence and particularly a G at position +3 are responsible for efficient NAD incorporation *in vivo*.

### Most NAD-RNAs are 3’-truncated

The observation that most yeast RNAs enriched in NAD captureSeq are much shorter than full-length mRNAs and their preferential mapping to mRNA 5’-ends lead us to ask whether there is an influence of NAD incorporation on the transcript length. Globally, we determined the percentage of mRNA-mapped full-length reads for WT and all mutants and compared this value for sample (+ADPRC, S) and negative control (-ADPRC, N). After normalization to non-enriched species, in all eight libraries the sample group contained less full-length reads than the negative control and more abortive fragments (**Figure S6A**). At the individual transcript level, we determined for the highly expressed (and enriched) TDH3 RNA the transcript start and end nucleotide analyzing each read individually (Tome et al., 2018). According to this analysis, both S and N groups feature the same dominating TSS (**Figures 6A and 6B**, histogram on top), while a dramatically different abundance and size distribution of truncated 3’-ends was observed between S group and N group. The proportion of full-length Illumina reads with identical TSS differed by a factor of 2.7 (33.4% in S and 88.8% in NC, **Figure 6A**, histograms to the right of the 2D plot). While the reasons for this increased proportion of 3’-truncated NAD-RNAs remain unclear, these findings may suggest that unidentified quality control mechanisms detect NAD incorporation into RNA as an error quite early and interfere with efficient transcript elongation.

**Figure 6.**
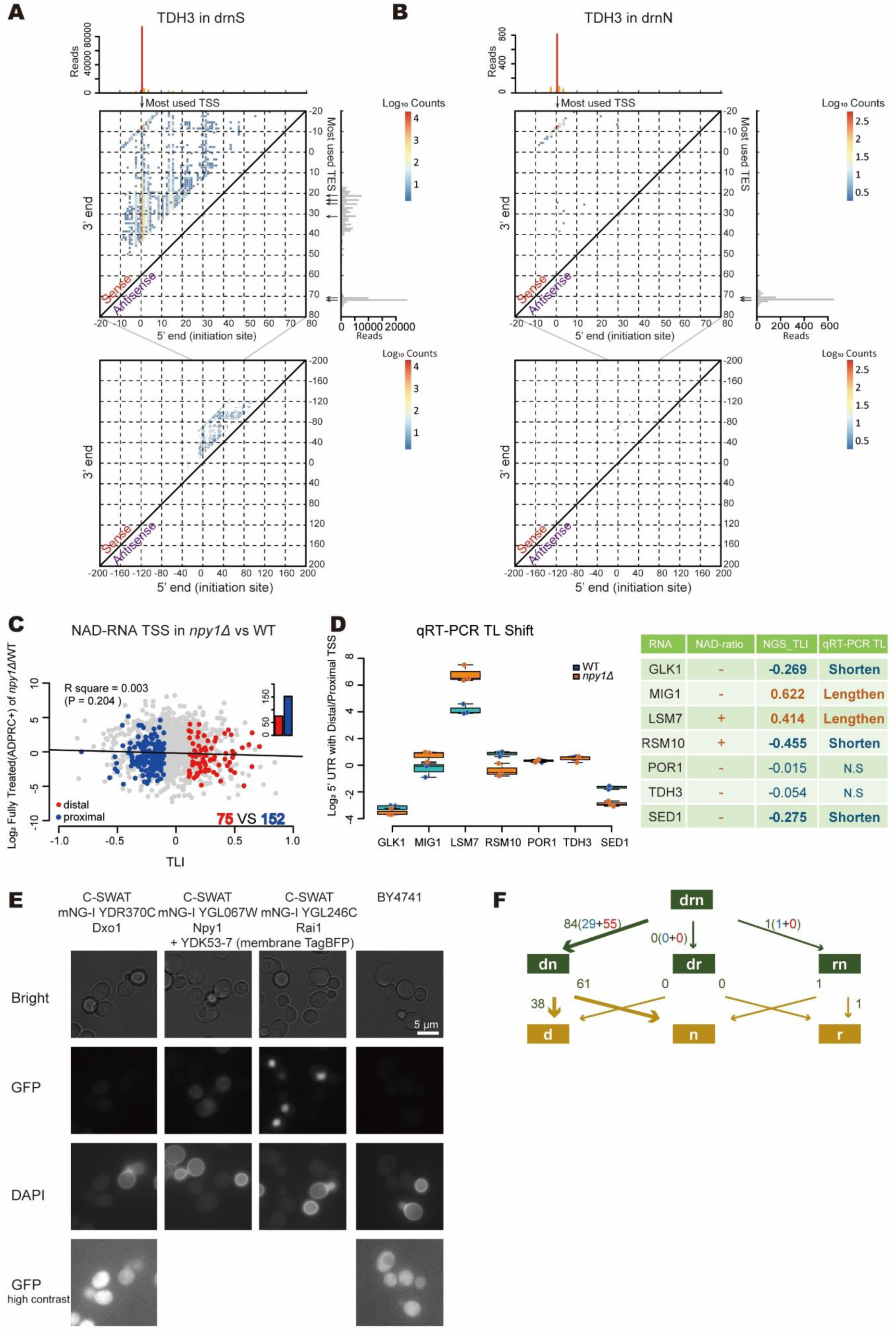
5’- and 3’-end heterogeneity of NAD-RNA and their modulation by decapping enzymes. (A, B) 2D NAD capping single transcript plot for TDH3 RNA in the *dxo1Δ rai1Δ npy1Δ* strain for sample group (+ADPRC, drnS, (A)) and N group (-ADPRC, drnN, (B)). 0 is the RefSeq TSS (-39 nt of the TLS site). Each bin represents a unique 5’ (initiation site, TSS, x axis) and 3’ (transcript end site, TES, y axis) pairing colored by the number of reads mapped to that bin. Expanded view below. For the 5-end initiation site histogram (above the 2D plot), the red bar indicates the ‘A’ position with Y<u>A</u>AG feature, while orange bars indicate ‘A’s without this feature, and grey bars indicate U/C/G. For the 3’-end histogram, no color differentiation was performed. Arrows represent preferred TSSs in the 5’-dimension and TESs in the 3’-dimension. (C) Global effect of the deletion of *NPY1 gene* on the transcript leader length by comparing NAD captureSeq read starts in the *npy1Δ* mutant with the WT (unfragmented libraries). A Transcript Leader length Index (TLI) >0 indicates that upon *npy1Δ* mutation the TSS towards more distal positions, while a TLI <0 indicates a proximal shift; RNAs with significantly (FDR <0.05) shifted TSS are shown as colored dots. Red: distal shift (TLI >0.1) blue: proximal shift (TLI <-0.1). The y axis indicates the relative NAD-RNA enrichment difference (ratio) from NAD captureSeq between the two strains. The bar charts represent the total number of RNA species with distal (red) or proximal (blue) TSS shift. (D) qRT-PCR with different primer pairs confirming different NAD-RNA transcript leader (TL) lengths in the enriched fractions from NAD captureSeq between WT (blue boxes) and *npy1Δ* strain (orange boxes). All parameters are as in Figure 4B. In the table, symbol ‘-’ and ‘+’ indicate a decrease or increase of the NAD modification ratio, respectively. ‘N.S indicates no significant difference. (E) Subcellular localization of Npy1, Dxo1, and Rai1. The target protein was labeled as C-SWAT Neonl fusion. Membrane TagBFP is the background control and was visualized in the DAPI channel. Due to the low intensity of the GFP signal for Dxo1, this image is shown again in high contrast. (F) Analysis of how many RNA species show a TL shift upon introduction of the first decapping enzyme (upper part, numbers next to the green errors), and how many of those disappear entirely or show reduced enrichment upon introduction of the second enzyme (lower part, numbers next to yellow arrows. TL shifting is defined by TLI >0.1 or <-0.1 and FDR <0.1. Blue and red numbers represent TL shifting to proximal and distal side, respectively. For example, the route from the drn triple knockout to the dn double knockout (i.e., introduction of Rai1) identifies 84 RNA species with shifted TL’s, 38 of which disappear or show reduced enrichment upon additional introduction of npy1 (going from the dn to the d mutant). Strain abbreviations are as in Figure 3A.

### Npy1, Dxo1, and Rai1 target different NAD-RNA populations and appear to act in a hierarchical order

The observation that the promoter motif got increasingly ‘blurry’ with increasing number of decapping enzymes present (**Figures 5A and S5E-K**) supported our assumption that the NAD captureSeq data actually reflect a superposition of RNAP II and decapping enzyme preferences. The comparison of the datasets of the three single mutants revealed extensive overlap, and 1544 species (>60%) were enriched in all three mutants, compared to the WT (**Figure S6B**). Similar findings were observed comparing the three double mutants. Computational sequence and secondary structure analysis of RNAs of uniquely or commonly enriched RNAs did not reveal specific features indicative of substrate preferences of these enzymes. However, for Rai1 we observed a slightly decreased minimum free energy of folding (Meijer et al., 2013) for preferred RNA substrates, compared to poor ones (**Figure S6C**). This finding may suggest that Rai1 tends to have a preference for less structured 5’-ends.

We noticed that the removal of decapping enzymes not only influenced the number of RNA species enriched in NAD captureSeq and their enrichment values, but also the (apparent) length of their 5’-ends (TL). This phenomenon was observed for ∼20% of all RNA species, and occurred in both directions, namely (apparent) TL lengthening and shortening upon knockout. For example, among the 1100 enriched sequences in common between the WT and *npy1Δ* strain, 152 apparently got shorter and 75 got longer TLs (**Figure 6C**). For all other mutants, similar observations were made. For several candidate RNAs, these length differences could be confirmed by qRT-PCR with the cDNA from the NAD captureSeq samples (**Figure 6D**). This phenomenon was observed almost exclusively for RNAs with read patterns indicative of pervasive transcription or multiple TSSs, and not for those with homogenous TLs. We assumed that the most likely explanation for these results may be that the decapping enzyme, when presented with a transcript mixture with different TLs, decaps some more rapidly than the others, due to sequence or structural preferences, thereby causing changes in the NAD captureSeq read profiles that look like shifted TSSs. A direct modulation of transcription (e.g., as transcription factors) is difficult to reconcile with the currently assumed roles and locations of these proteins, at least for Dxo1 and Npy1.

Rai1 has been reported as a nuclear protein and was detected as a component of the RNAP II elongation complex (Harlen and Churchman, 2017), while for Dxo1 both nuclear and cytosolic locations were claimed (Chang et al., 2012). Npy1 was described as a peroxysomal protein (AbdelRaheim et al., 2001). Localization microscopy using three different C-SWAT fluorescent protein fusions (Meurer et al., 2018) for each candidate gene revealed strong localized fluorescence in the nucleus for Rai1, while Dxo1 showed only a very weak and ubiquitous fluorescence (**Figure 6E**), consistent with the reported localizations of these enzymes. For Npy1, however, a rather homogenous cellular distribution without enrichment at specific sites was observed, consistent with cytosolic localization (**Figure 6E**). This localization may imply a temporal order, in which Rai1 processes its NAD-RNA substrates during or shortly after transcription, while Npy1 can only act once the transcripts (or their primary degradation products) arrive in the cytosol. For Dxo1, both options are conceivable. Therefore, we tried to find evidence in our NAD captureSeq data for a temporal order of processing by these enzymes. In particular, we searched for examples where – when starting with the triple knockout and then “adding” one by one the decapping enzymes (i.e., comparing the triple knockout with the appropriate double and single knockouts) – a significant TL length change is observed upon “addition” of the first decapping enzyme (suggesting that this enzyme decaps a fraction but not all TL variants) and upon “addition” of the second one the transcript disappears entirely (or is significantly reduced in enrichment) from the enriched fraction (suggesting that the second enzyme decaps the remaining TLs). Indeed, from the 84 species with TL length changes between triple knockout and *dxo1Δ npy1Δ* double knockout, 61 disappeared in the *npy1Δ* single knockout and 38 in the *dxo1Δ* single knockout. Importantly, hardly any examples were found for the pathways via the other double mutants (1 example for *npy1Δ rai1Δ* and 0 for *dxo1Δ rai1Δ*) (**Figure 6F**). These findings are consistent with our assumption that Rai1 is the first player in NAD-RNA decapping.

### NAD-RNAs do not support translation in budding yeast

Finally, we tested whether the NAD cap in combination with different TL lengths and sequences may modulate translation. Reports on NAD-RNA translatability are conflicting: Jiao et al. had reported that NAD-RNA is not translated in human (HEK293T) cell extracts, based on a single mRNA luciferase construct with a single fixed TL sequence (Jiao et al., 2017) while a recent study in the model plant *Arabidopsis thaliana* demonstrated that NAD-capped mRNAs are enriched in the polysomal fraction, associate with translating ribosomes, and can probably be translated (Wang et al., 2019). No data for yeast have been reported yet. For seven different mRNAs, we prepared luciferase fusion constructs with long and short TLs by *in vitro* transcription, followed by removal of the accompanying ppp-RNA by treatment with polyphosphatase and exonuclease Xrn-1. While the control constructs harboring an m^7^G-capped 5’-end were efficiently translated in a yeast *in vitro* extract and showed significant differences in luminescence depending on the TL length (Gilbert et al., 2007; Rojas-Duran and Gilbert, 2012), NAD-capped RNA was not translated to any significant extent, even less than ppp-RNA and p-RNA of the same sequence (**Figure 7A**). These results suggest that NAD-capped RNAs (at least the nuclear transcripts investigated here) are not translated in budding yeast.

**Figure 7.**
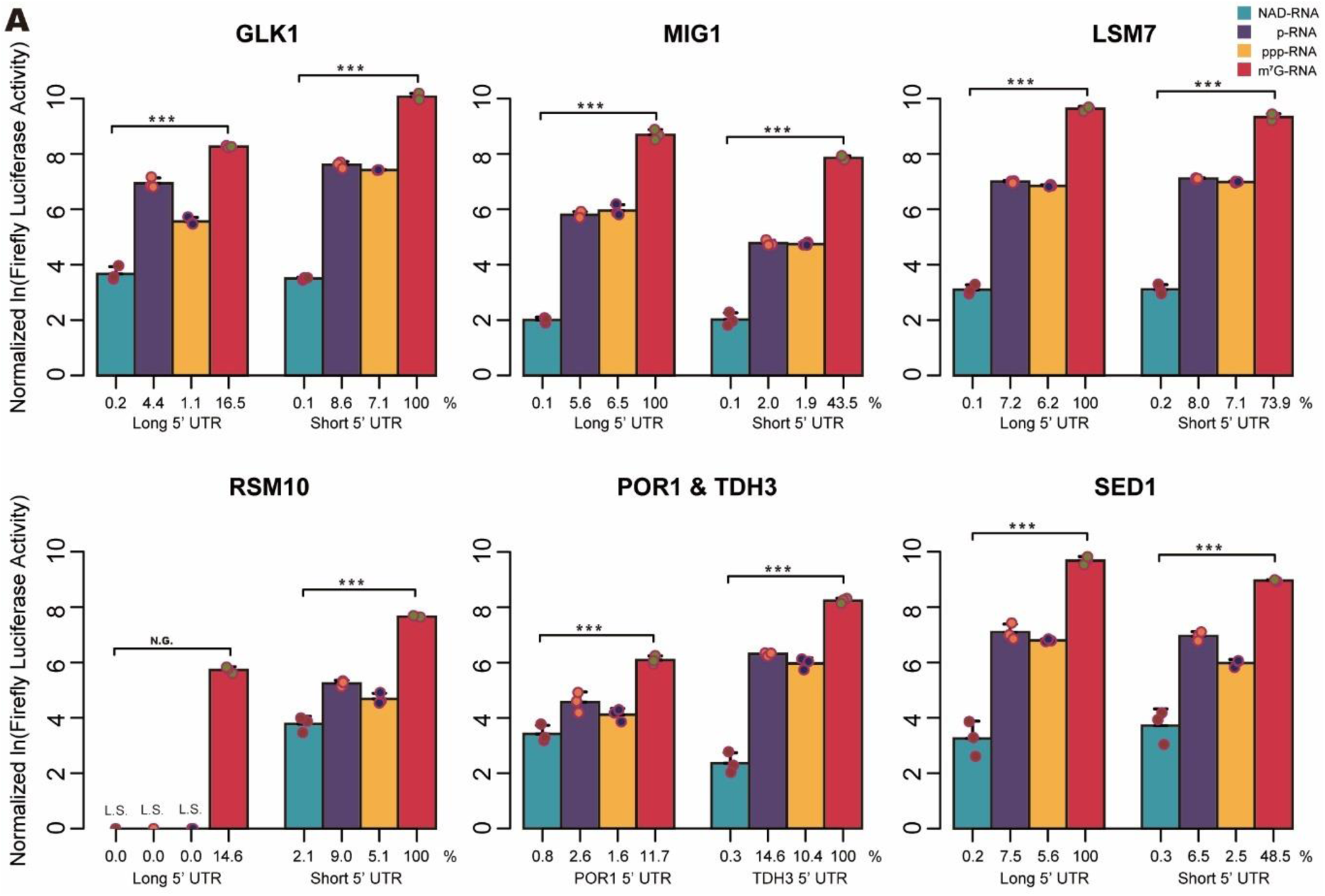
*In vitro* translation of NAD-RNA with shifted TL. (A) *In vitro* translation of NAD-capped RNA, p-RNA, ppp-RNA, and m^7^G-RNA with short or long TL sequences. mRNAs contain the alternative TL sequences identified in NAD captureSeq, followed by 22nt of the CDS of the corresponding gene, followed by firefly luciferase (1653nt) and poly(A)_30_. And reference m7G-capped mRNA contains renilla luciferase (936nt) and poly(A)_30_. Firefly luciferase activity was normalized to renilla luciferase, then normalized to the C_t_ value of the full-length mRNA determined by qRT-PCR. NAD-capped mRNAs, p-mRNAs, and ppp-mRNAs contain the same sequence as the corresponding m^7^G-capped RNAs. Logarithmic representation of normalized luciferase activity. Blue - NAD-RNA; purple – p-RNA; orange – ppp-RNA; red – m^7^G-RNA. The percentage values below the bars indicate relative luciferase activity normalized to the m^7^G-capped mRNA of that species with the higher luciferase expression. L.S. represents luciferase signal was not significantly above the background. Error bars represent mean + sd, n=3. p values are denoted by asterisks: (*) p <0.05; (**) p <0.01; (***) p <0.005 (ANOVA); (N.S.) not significant. Arrows indicated TSS shifting direction after removal of Npy1.

## Discussion

Taken together, our results indicate that in budding yeast, NADylation of RNAs is a very common phenomenon. A previous study reported only 37 species enriched in NAD captureSeq in budding yeast grown in the same medium (Walters et al., 2017). This study, however, focused on full-length mRNAs, and used a library preparation protocol that discarded the small-RNA fraction (⪅200 nt). Our work confirms that there are hardly any full-length NAD-mRNAs, but additionally reveals a rich landscape of thousands of short NAD-mRNA fragments whose purpose is apparently not to encode for proteins.

This conclusion may not apply in yeast mitochondria, however, where several lines of evidence suggest that transcriptional incorporation of the coenzyme is, in fact, an evolved feature. First, the mitochondrial transcription machinery exhibits NAD-mediated RNA initiation efficiencies that are at least 10-fold higher, compared to the nuclear RNA polymerase II (Bird et al., 2018). Second, individual mRNA species in this organelle were found to be highly 5’-NAD modified, comprising up to 60% of the respective transcript pools. And third, the redox state of mitochondrial NAD caps was observed to vary, depending on the metabolic growth conditions of the cell (Bird et al., 2018). These findings may indeed indicate a regulatory role of the NAD cap in this organelle, that harbors redox-intensive energy conversion pathways highly dependent upon the coenzyme, and are contrasting sharply with our discovery of a tightly policed landscape of nuclear-derived NAD-mRNA fragments, subject to perpetual decay.

NAD is initially incorporated into several thousands of transcripts by transcription initiation by RNAP II in a largely statistical manner reflecting the competition of NAD with the canonical initiator ATP. As in prokaryotes (Bird et al., 2016; Frindert et al., 2018; Vvedenskaya et al., 2018), the promoter sequence determines the efficiency of NAD incorporation, which is for most yeast nuclear transcripts between 1 and 5%. We observe that a Y<u>A</u>AG motif supports efficient NAD incorporation by RNAPII *in vivo*, with the G at position 3 being particularly important. In contrast, a preference for HRR<u>A</u>SWW was reported for *E. coli* RNAP (Vvedenskaya et al., 2018), W<u>A</u>RR for *B. subtilis* RNAP (Frindert et al., 2018), and R<u>A</u> for yeast mtRNAP (Bird et al., 2018), with the underlined <u>A</u> always indicating the TSS. We find that, as a consequence of this promoter dependence, for many RNAs the NADylated species originate from different TSSs and have therefore different (shorter or longer) 5’-UTRs than the canonical ones. This phenomenon may modulate the secondary structure of these RNAs and hence their stability, molecular interactions, and biological fate. Of note, the discovery of alternative TSS selection and the YAAG core promoter motif have been made possible by the combination of 5’-end selection by ADPRC treatment and ligation-based attachment of the 5’-adapter, which allowed the determination of NAD-RNA 5’-ends with single-nucleotide precision, in contrast to random-primed library preparation methods that create heterogeneous ends.

NAD-RNAs are – on average – shorter than non-NAD-RNAs and only rarely reach the size of a typical primary mRNA transcript. The most likely explanation is that some unidentified quality control mechanism detects 5’-NADylation of RNA as an error early during transcription and prevents efficient elongation, as it does with uncapped or incompletely capped transcripts (Bresson and Tollervey, 2018). Alternatively, NAD-RNAs might be subject to accelerated degradation after transcription is complete, but it is unclear how 5’-NAD can accelerate degradation at the 3’-end.

The discovery that budding yeast maintains at least three different, partly redundant, pathways for NAD cap removal, using enzymes with different chemistry and cellular localization, implies that decapping unwanted NAD-RNAs is important for the cell. Our data are in agreement with the hypothesis that Rai1 acts earlier than the other two enzymes. As the nuclear protein Rai1 is known to associate with RNAPII during elongation (Harlen and Churchman, 2017) and to act in RNA surveillance by assisting the 5’- to 3’-exonuclease Rat1 in the co-transcriptional degradation of uncapped transcripts (Jiao et al., 2010; Kim et al., 2004), such an order appears plausible.

The observed combination of the low efficiency of RNAPII transcription initiation by NAD, the reduced length of NAD-RNAs, and the abundance of NAD-RNA decapping enzymes warrants that hardly any NAD-RNAs occur in the cell that could give rise to translation into proteins. Our data indicate, however, that yeast ribosomes, like mammalian ones (Jiao et al., 2017), hardly translate synthetic NAD-mRNAs, suggesting that the ribosomal machinery contains additional safeguards against NAD-mRNAs. Thus, budding yeast protects itself at different stages of gene expression against NAD-RNA.

## STAR METHODS

Detailed methods are provided in the online version of this paper and include the following:

- KEY RESOURCES TABLE
- LEAD CONTACT AND MATERIALS AVAILABILITY
- EXPERIMENTAL MODEL AND SUBJECT DETAILS
- METHOD DETAILS
  ∘ Fluorescence Microscopy and Colony Fluorescence Imaging
  ∘ Total RNA Isolation and Purification
  ∘ NAD captureSeq and Transcriptome Libraries Preparation
  ∘ NGS Analysis
  ∘ Proteomics Sample Preparation and TMT Labeling
  ∘ Proteomics Mass Spectrometry Data Acquisition and Analysis
  ∘ RNA Pull-down and UPLC-MS Analysis
  ∘ Flow Cytometry Data Acquisition and Analysis
  ∘ Gel Electrophoresis
  ∘ *In vitro* Transcription and NAD/ppp/p/m^7^G-capped RNA Preparation
  ∘ Plasmid Construction
  ∘ gDNA Extraction
  ∘ Cell-free Extraction and *in vitro* Translation
  ∘ Cellular NAD quantification
  ∘ Protein Expression and Purification
  ∘ RNA in vitro Decapping and NAD Hydrolysis Kinetic Assays
  ∘ Determination of RNA NAD-Capping Ratios in Total RNA and *in vitro* transcribed NAD-/ppp-mRNA Mixtures
  ∘ Quantitative Reverse Transcription PCR (RT-qPCR) and Standard PCR Procedures
- QUANTIFICATION AND STATISTICAL ANALYSIS
- DATA AND SOFTWARE AVAILABILITY

## SUPPLEMENTAL INFORMATION

Supplemental Information can be found online at:

## ACKNOWLEDGMENTS

The authors would like to thank Jäschke Lab members, V. Winkler (Heidelberg University), and A. Hotz-Wagenblatt (DKFZ, Heidelberg) for discussions, H.C. Lee and D. Grimm for strains and plasmids, M. Brunner for access to LightCycler, ZMBH Flow Cytometry & FACS Core Facility for the FCM measurement, M. Rettel and F. Stein (EMBL Proteomics Core Facility, Heidelberg) for proteomics analysis, and bwHPC (BwForCluster MLS&WISO) for cluster computation resources. This work was supported by the German Research Foundation (DFG, grant # Ja794/10, SPP 1784, to A.J.).

## AUTHOR CONTRIBUTIONS

Conceptualization: Y.Z. and A.J., Methodology: Y.Z., D.Ku., T.S., D.Ki., G.N., D.I., and V.B., Investigation: Y.Z., D.Ku., T.S., D.Ki., G.N., D.I., H.H., Formal Analysis: Y.Z., M.K., H.H., and A.J., Supervision: H.H., M.K., and A.J., Administration: A.J., Writing – Original Draft: Y.Z. and A.J., Writing – Review and Editing: all authors.

---

**Figure S1.**
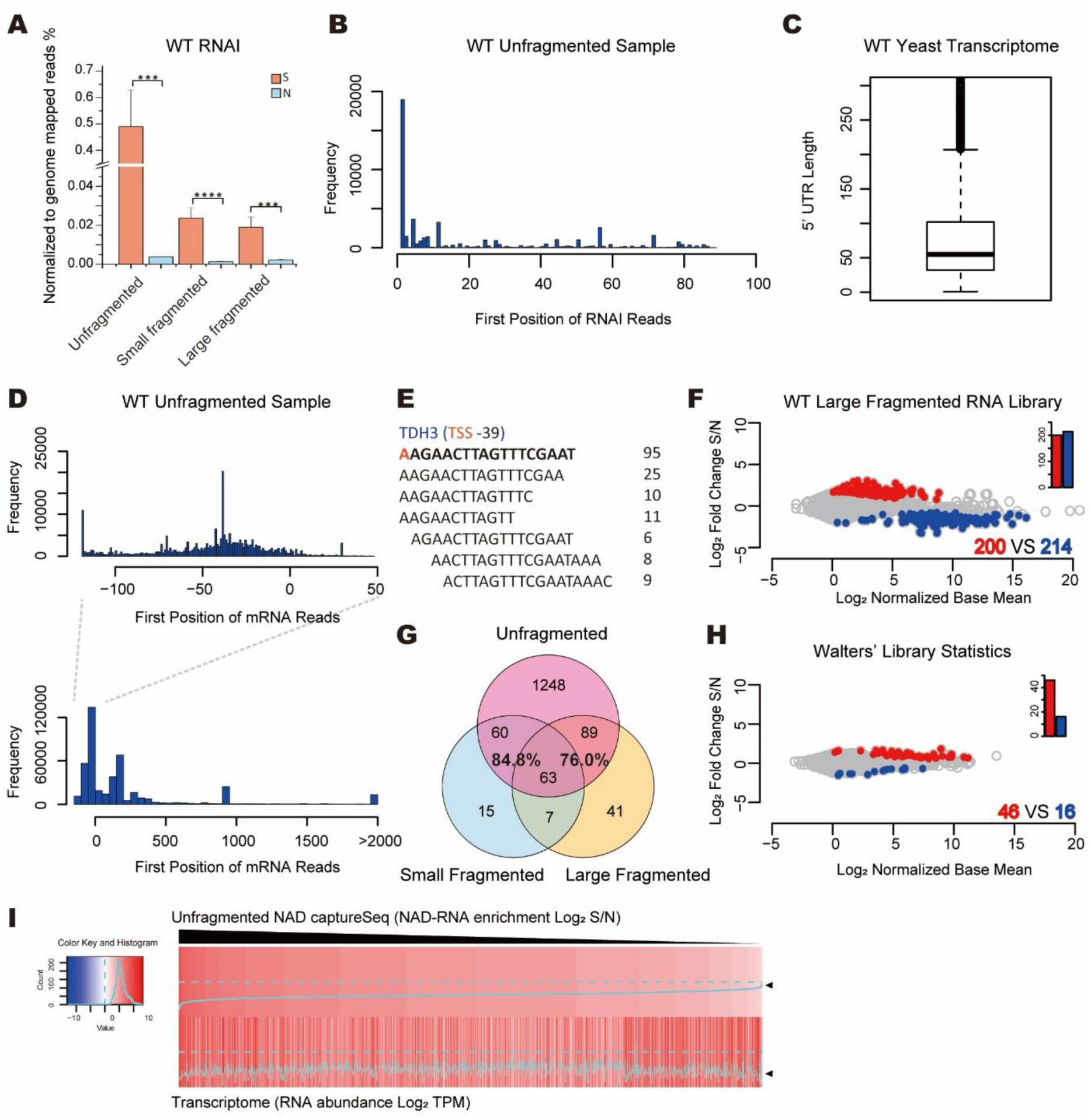
NAD captureSeq library reference features and further analysis of the WT strain. (A) Method validation: Enrichment of spike-in NAD-RNAI (a regulatory RNA from *E. coli*) reads in WT NAD captureSeq. The height of the orange bars indicates the percentage of NAD-RNAI reads among total genome-mapped reads in the sample group (S, ADPRC fully-treated), and blue bar height indicates the same in the negative control group (N, minus ADPRC). Error bars represent mean + standard deviation (SD), n=3. p values are denoted by asterisks: (*) p <0.05; (**) p <0.01; (***) p <0.001; (****) p <0.0001 (Student’s t test). This analysis revealed efficient enrichment of the synthetic pure NAD-RNA in all samples. Particularly high enrichment was observed in the unfragmented libraries. (B) Positions of the first nucleotide of spike-in RNAI-mapped reads in the WT sample, confirming the reliable identification of NAD-RNA 5’-ends from the NAD captureSeq data. (C) Distribution of 5’ UTR lengths in *S. cerevisiae* according to published data (Nagalakshmi et al., 2008). The boxplot shows from bottom to top “minimum” (Q1-1.5 interquartile range (IQR, 25% to 75%)), first quartile(Q1, 25%), median (solid line, 50%), third quartile (Q3,75%), “maximum” (Q3+1.5IQR), and outliers (black dots). (D) Positions of the first nucleotide of genome-mapped reads in the WT unfragmented NAD captureSeq sample group. The 5’ UTR region (-120 to +50, relative to the translation start site (TLS) as ‘0’) is zoomed in in the upper panel, confirming the 5’ UTR length distribution expected from the literature data visualized in Figure S1C. (E) Alignment of small RNAs (12-17 nt) with homology to TDH3 RNA observed in the WT unfragmented NAD captureSeq library. The red ‘A’ is the +1 nucleotide of the assumed transcription start site. The number on the right side of the sequence is the number of reads. (F) Enriched NAD-RNAs from large fragmented NAD captureSeq on the WT strain. All statistics parameters are as in Figure 1A. Biological triplicates. (G) Intersection of enriched NAD-RNA species among the unfragmented, small fragmented, and large fragmented WT NAD captureSeq libraries. The percentage values are defined as number of shared species, divided by total number of species in fragmented library. (H) Enriched NAD-RNA from Walters’ published WT BY4742 yeast library (Walters et al., 2017). All statistics parameters are as in Figure 1A. (I) Heatmap correlation between NAD-RNA enrichment (NAD captureSeq) and transcript abundance (transcriptome sequencing).

**Figure S2.**
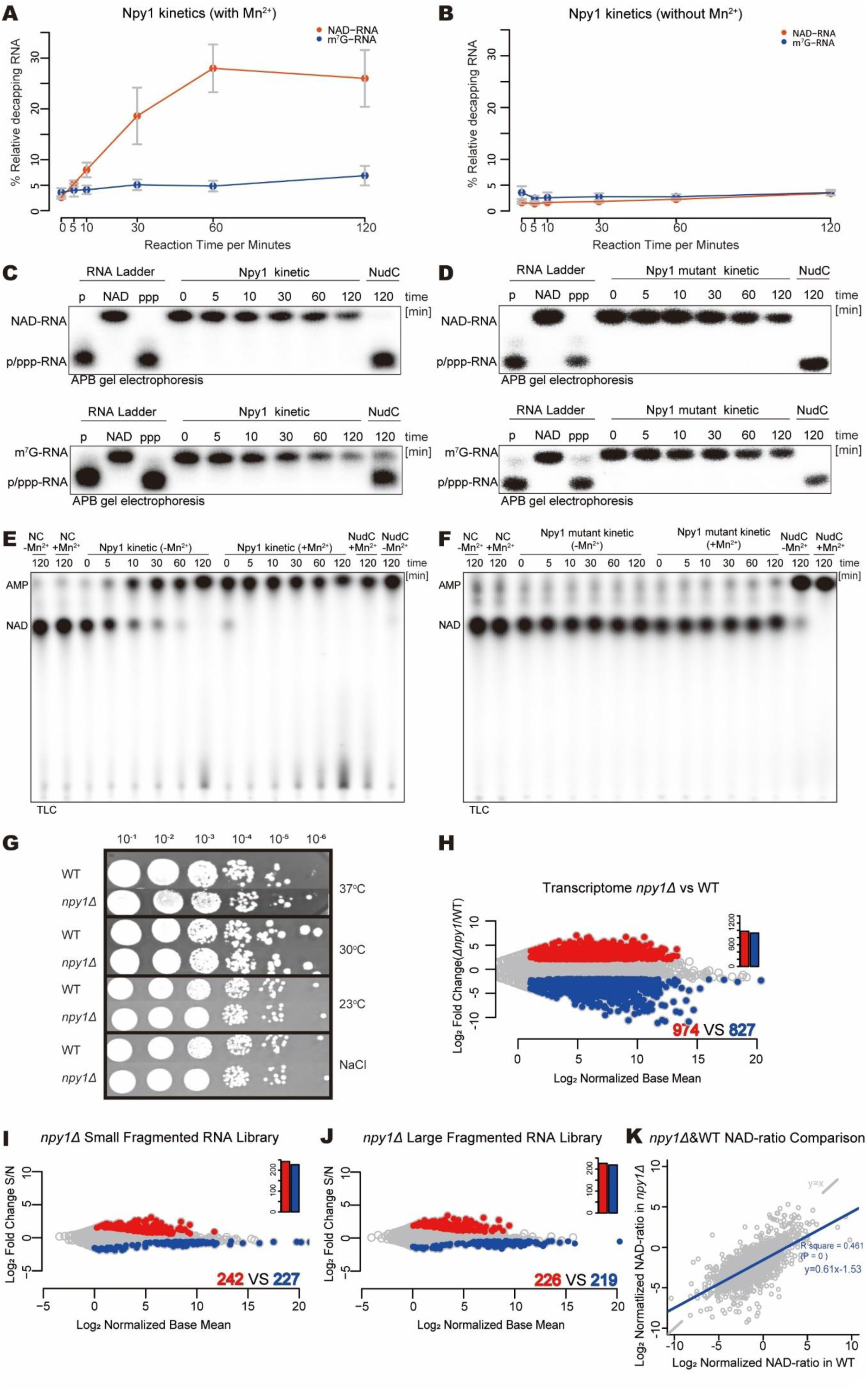
Npy1 kinetics and comparison with WT NAD captureSeq. (A) & (B) Quantification curve of Npy1 kinetics with NAD- and m^7^G-capped TDH3 RNA in the presence and absence of Mn^2+^ as shown in Figure 2A and S2C. Error bars represent mean ± standard deviation (SD), n=3. (C) Npy1 WT kinetics of decapping NAD- and m^7^G-capped RNA without Mn^2+^ *in vitro*. All conditions as in Figure 2A. (D) Npy1 mutant (E276Q) kinetics of decapping NAD- and m^7^G-capped RNA with Mn^2+^ *in vitro*. (E) & (F) Hydrolysis of NAD (into NMN and ATP) by Npy1 WT and mutant (E276Q) Npy1 *in vitro*. ^32^P-NAD was treated with the respective enzyme in the presence of 2 mM Mg^2+^ and 1 mM Mn^2+^ and reaction mixtures separated by thin layer chromatography (TLC, NH_4_OAc/EtOH 4:6). (G) Growth phenotype comparison between WT and *npy1Δ* strain under different conditions. Cells were spotted in 10-fold serial dilution starting from OD_600_ = 1.. The cells were cultured in normal YPD medium at 30 °C, while the NaCl set was additionally supplemented with 0.5 M NaCl. (H) Analysis of expression level changes of transcripts upon removal of Npy1 by transcriptome sequencing. 7620 different transcripts are analyzed and represented as a dot on the plot. Red dots: up-regulated transcripts (fold change >1.414, normalized base mean >1, p <0.05), blue dots: down-regulated transcripts (fold change <0.707, normalized base mean >1, p <0.05). Biological triplicates. (I) Enriched NAD-RNAs from small fragmented NAD captureSeq on the *npy1Δ* strain. Other parameters are as in Figure 1A. n=3. (J) Enriched NAD-RNAs from large fragmented NAD captureSeq on the *npy1Δ* strain. Other parameters are as in Figure 1A. n=3. (K) Relative RNA NAD-modification ratio trend from integration of NADcaptureSeq, transcriptome, and LC-MS data. Grey dashed line is y=x as reference, while the blue solid line represents the linear regression of the experimental data: NAD-ratio_ΔNpy1_=k* NAD-ratio_WT_+c (k is slope and c is intercept).

**Figure S3.**
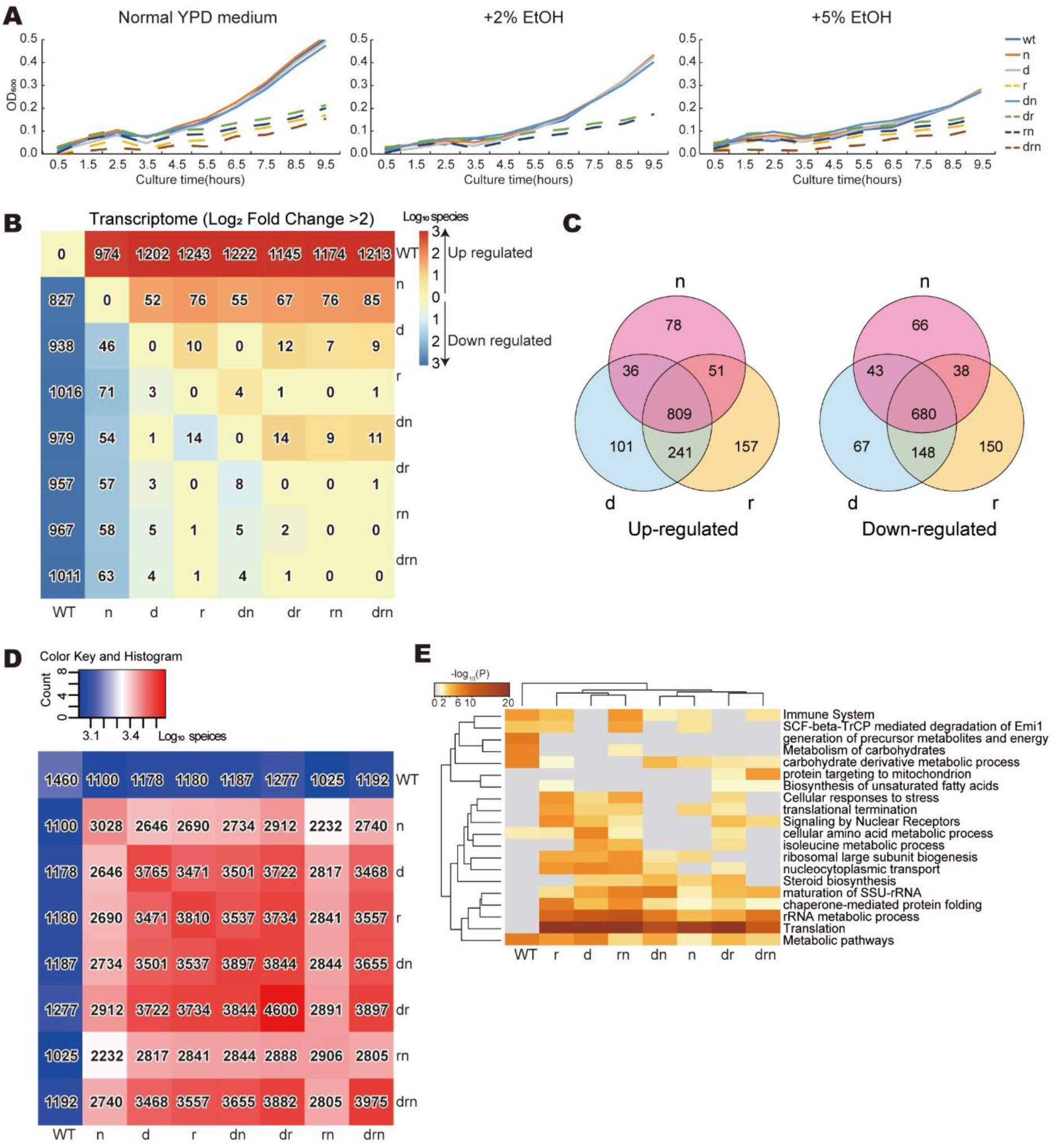
Further analysis of deletion mutants. (A) Time course of the cell density (OD_600_) for WT and all deletion mutants. The mutant strains having a *RAI1* gene deletion are indicated by dashed lines while all other strains are plotted with normal lines. The cells were incubated in YPD medium on 96 well plates at 30 °C with shaking. The cell density was measured by TECAN at certain time points. Strain abbreviations are as in Figure 3A. (B) Heatmap of the differential expression assessed by transcriptome sequencing, comparing the number of up-regulated (red, above the “0 0 0” diagonal) and down-regulated (blue, below the “0 0 0” diagonal) transcripts between two strains. The number of RNA species was log_10_-transformed and scaled by color intensity. Example: Comparison of *npy1Δ* and WT strain yields 974 species with up-regulation and 827 species with down-regulation. (C) Intersection of up-/down-regulated RNA species between the three single-deletion strains by transcriptome sequencing. (D) Heatmap of the intersection of enriched NAD-RNAs assessed by NAD captureSeq. The number of RNA species that overlap between two strains was log_10_ transformed and scaled by color intensity. (E) Heatmap of functional clustering of the top 250 enriched NAD-RNA species for WT and all 7 deletion strains. The color intensity represents the log_10_(p value).

**Figure S4.**
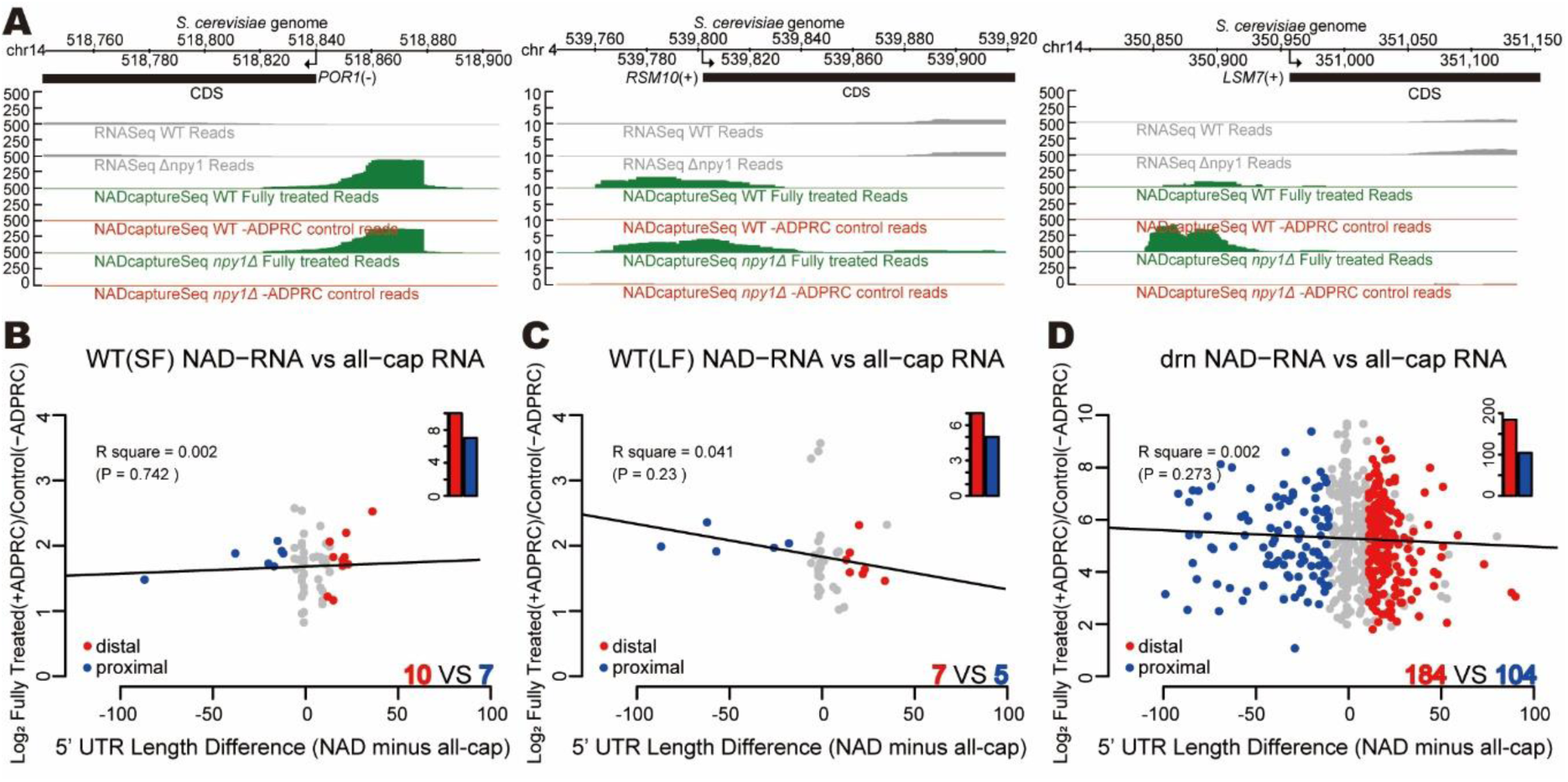
Genome-wide TL features of NAD-RNAs. (A) Comparison of the read profiles of WT and *npy1Δ* samples. Aligned reads of NAD captureSeq and transcriptome sequencing were normalized as RPM and visualized in the IGB. Green patterns represent accumulated reads in the fully treated sample group (+ADPRC), while the red traces are derived from the – ADPRC negative control in NAD captureSeq (unfragmented libraries). The grey patterns represent the read distribution of transcripts from transcriptome sequencing. (B) Same experiment as in Figure 4C, but using the small fragmented WT NAD captureSeq libraries. (C) Same experiment as in Figure 4C, but using the large fragmented WT NAD captureSeq libraries. (D) Same experiment as in Figure 4C, but using the *dxo1Δ rai1Δ npy1Δ* triple knockout NAD captureSeq libraries.

**Figure S5.**
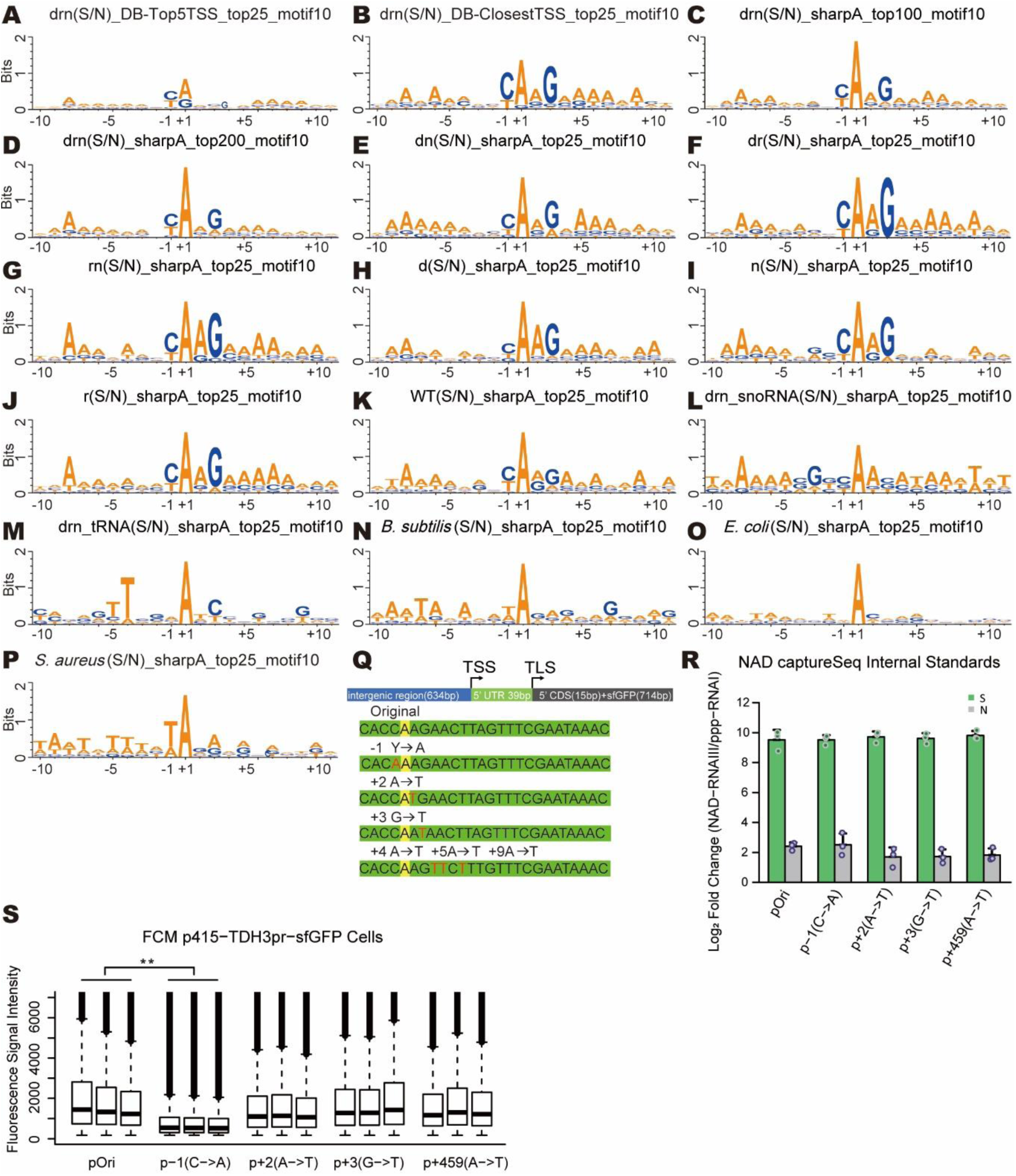
Key promoter YAAG motif in other mutants and organisms. (A-M) Motif analysis of the -10 to +10 region around the TSS based on NAD captureSeq data as in Figure 5A for different selected populations and mutant strains. Top 100 and top 200 represent 100 and 200 most enriched NAD-mRNA species, respectively. Enriched NAD-capped snoRNA and tRNA are represent snoRNA(S/N) and tRNA(S/N), respectively. Strain abbreviations are as in Figure 3E. (N-P) Motif analysis of the top 25 enriched NAD-mRNA species in *B. subtilis* (Frindert et al., 2018*), E. coli* (Cahova et al., 2015), and *S. aureus* (Morales-Filloy et al., 2020*), respectively. All parameters are as in Figure 4E.* (Q) *Scheme illustrating the TDH3* gene promoter and relevant mutations. The yellow ‘A’ is the TSS and referenced as +1. The red letters highlight mutations. (R) Spike-in standards NAD-RNAIII and ppp-RNAI for the standards of TDH3 RNA NAD-ratio quantification. Bar heights, color, dots and error bars are as in Figure 5C. (S) Flow cytometry analysis and analysis of GFP expression levels of the yeast strains shown in Figure 5D, using three independent clones. The solid lines in the boxplot represent the median fluorescence signal intensity for each group (>75000 events in pOri, p-1, p+2, p+3, and p+459). The median values of each group were compared, p value is denoted by asterisks: (**) p <0.01 (Student’s t test). Outliers above 7000 are omitted.

**Figure S6.**
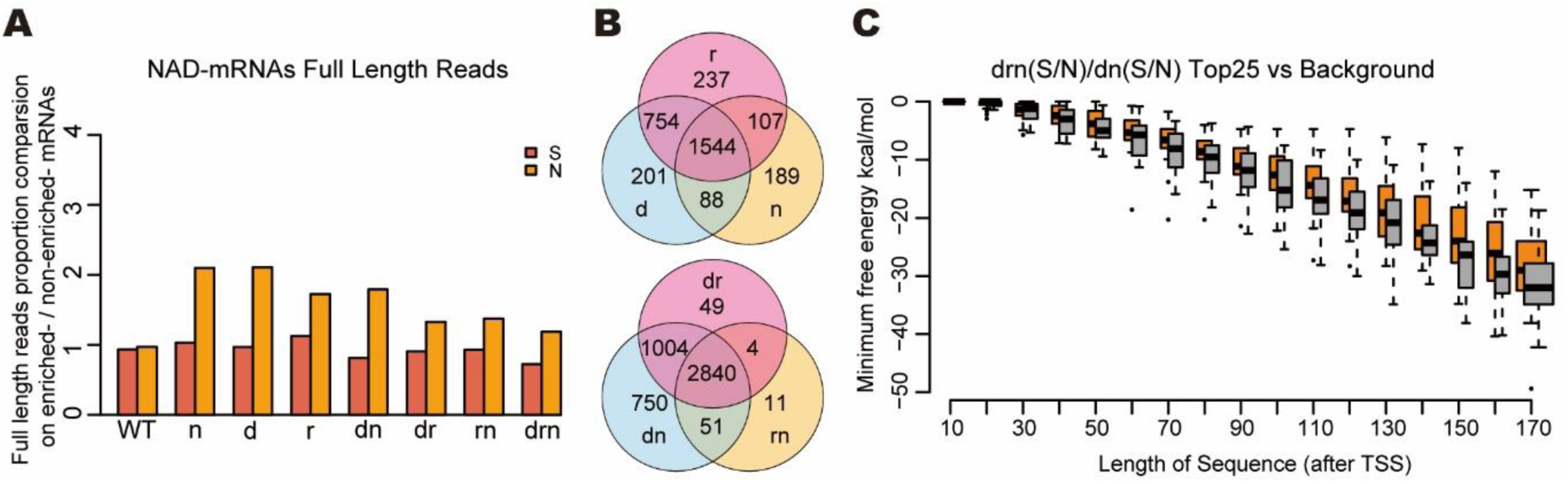
Decapping enzymes target different NAD-RNAs. (A) Ratio of full length reads of mRNAs in NAD captureSeq libraries. The bar heights indicate the proportion of full length reads from enriched NAD-RNA species divided by the proportion from non-enriched RNA species. The red bars represent the percentage ratio in NAD captureSeq sample group (+ADPRC, S), and orange bars represent in NAD captureSeq negative control group (-ADPRC, N). Strain abbreviations are as in Figure 3A. (B) Intersection of the enriched NAD-RNA species identified in the three single mutant (top) and double mutant strains (bottom, unfragmented libraries). (C) Prediction of TL folding energy using RNAfold (Meijer et al., 2013) as a function of the assumed transcript length. The orange bars represent the predicted minimum free energy of the top 25 highest-enriched NAD-RNAs when comparing the drn triple knockout (S/N) with the dn double knockout (S/N), while the grey bars refer to background RNAs (no significant NAD enrichment change (0.707 < drn(S/N)/dn(S/N) < 1.414) between the two strains).

## STAR METHODS

### KEY RESOURCES TABLE

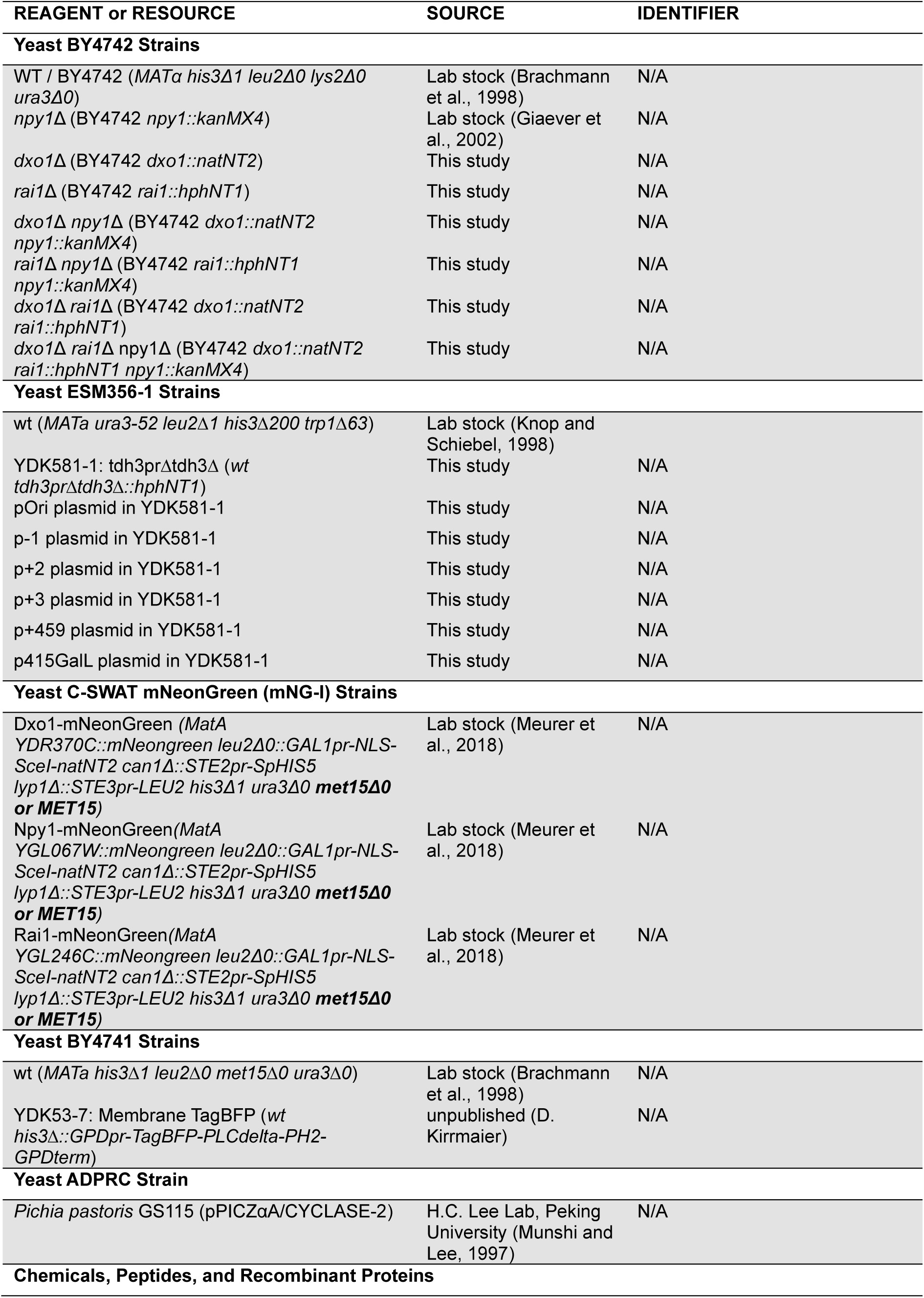
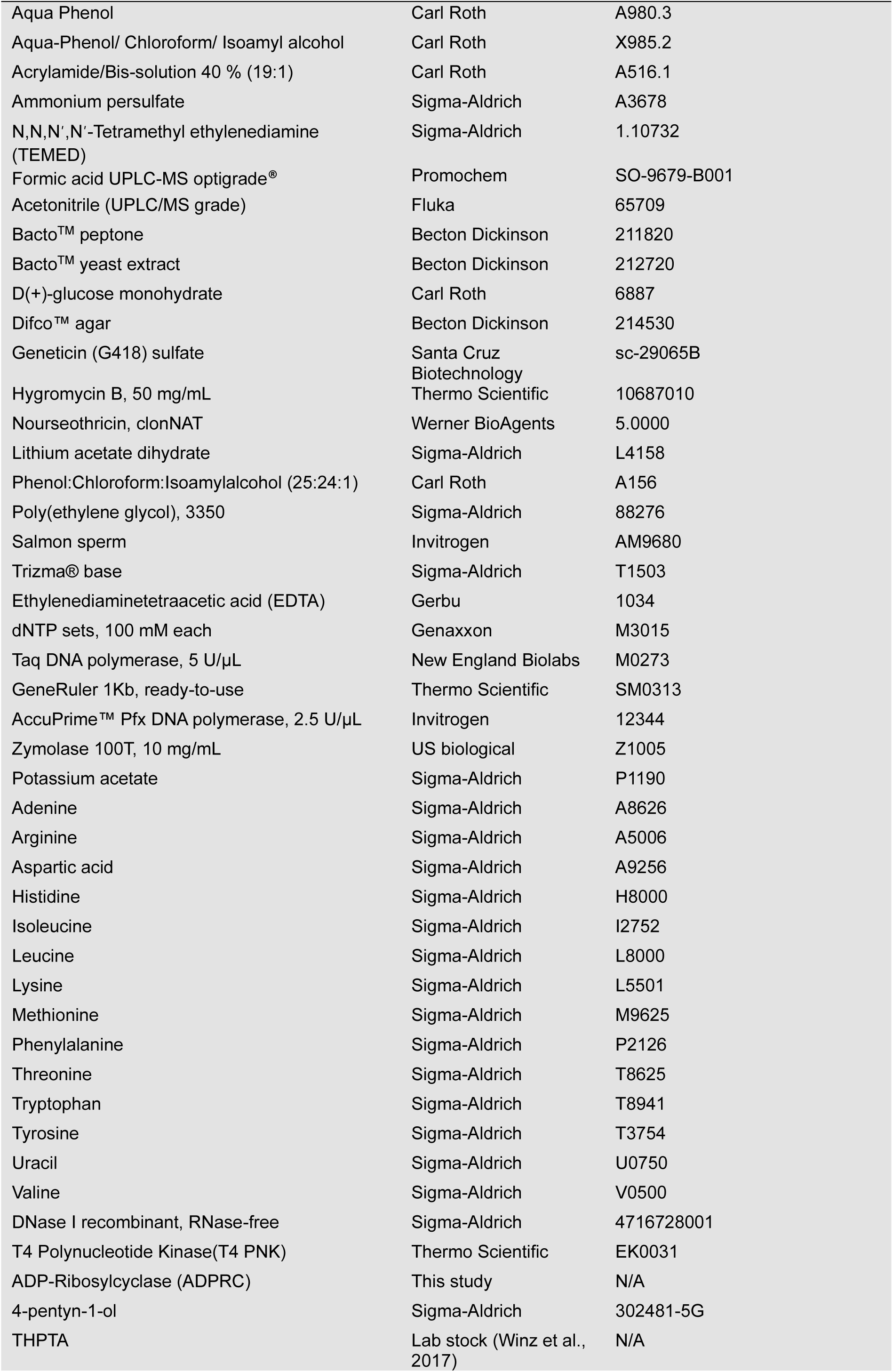
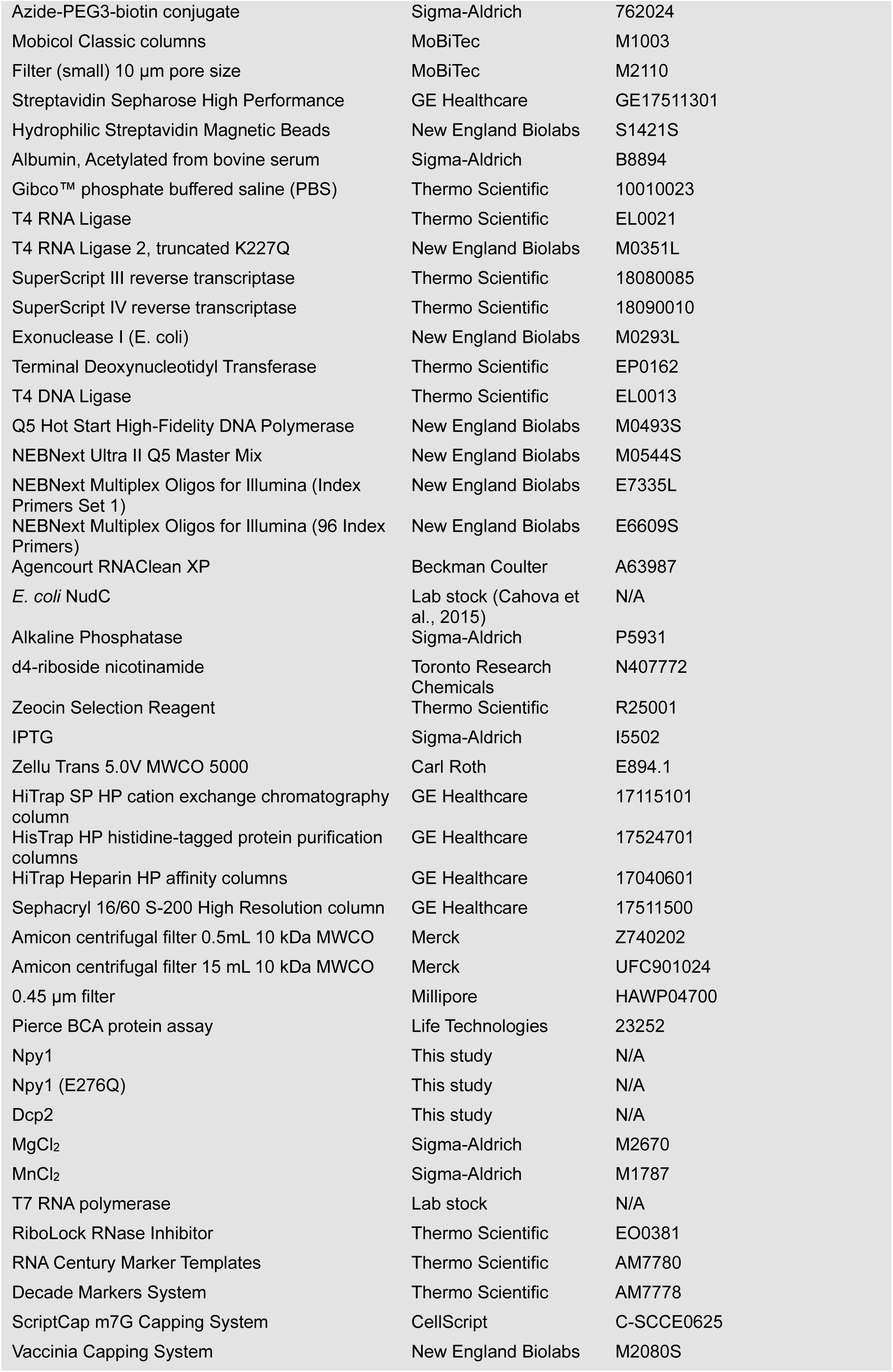
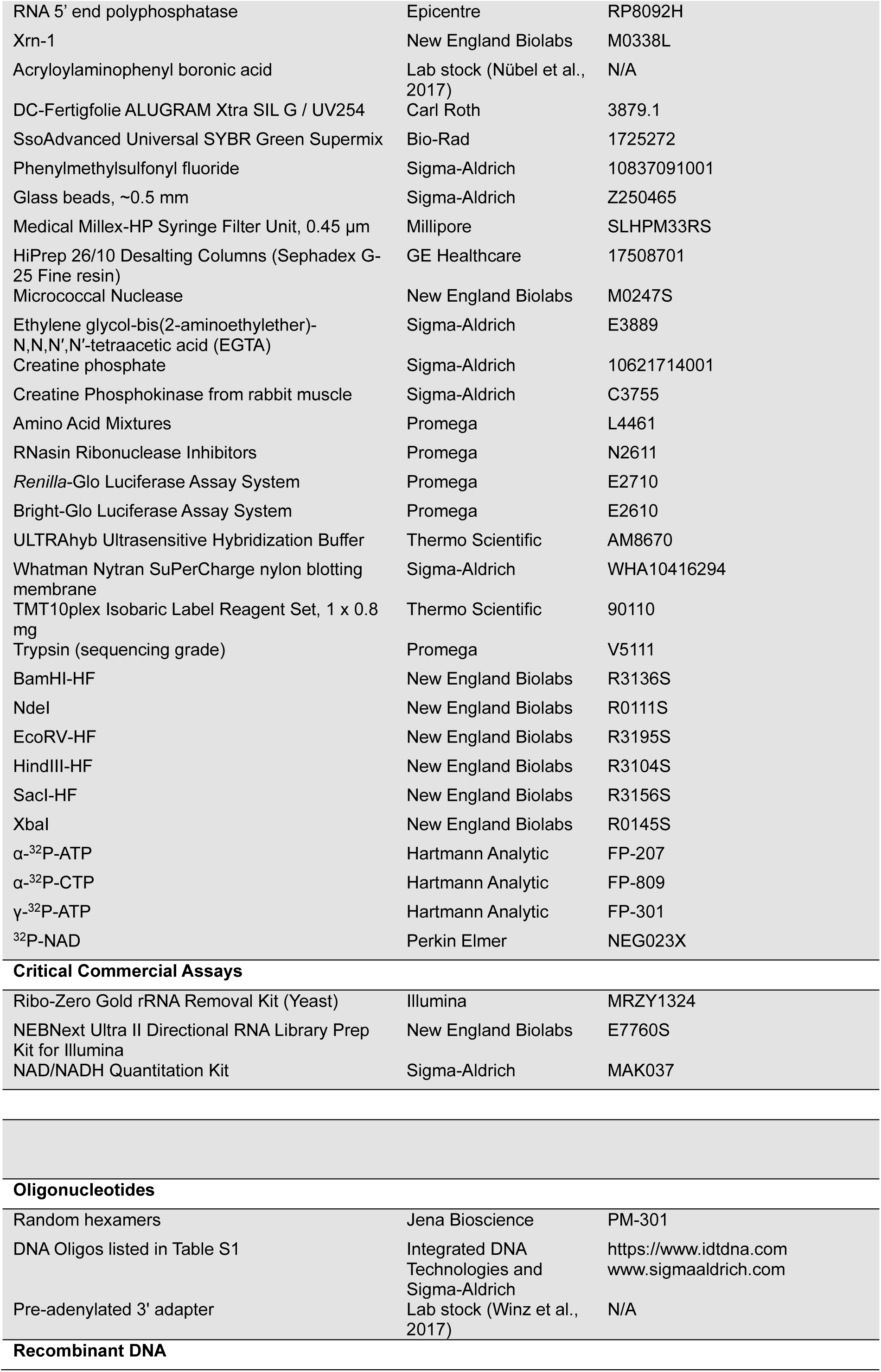
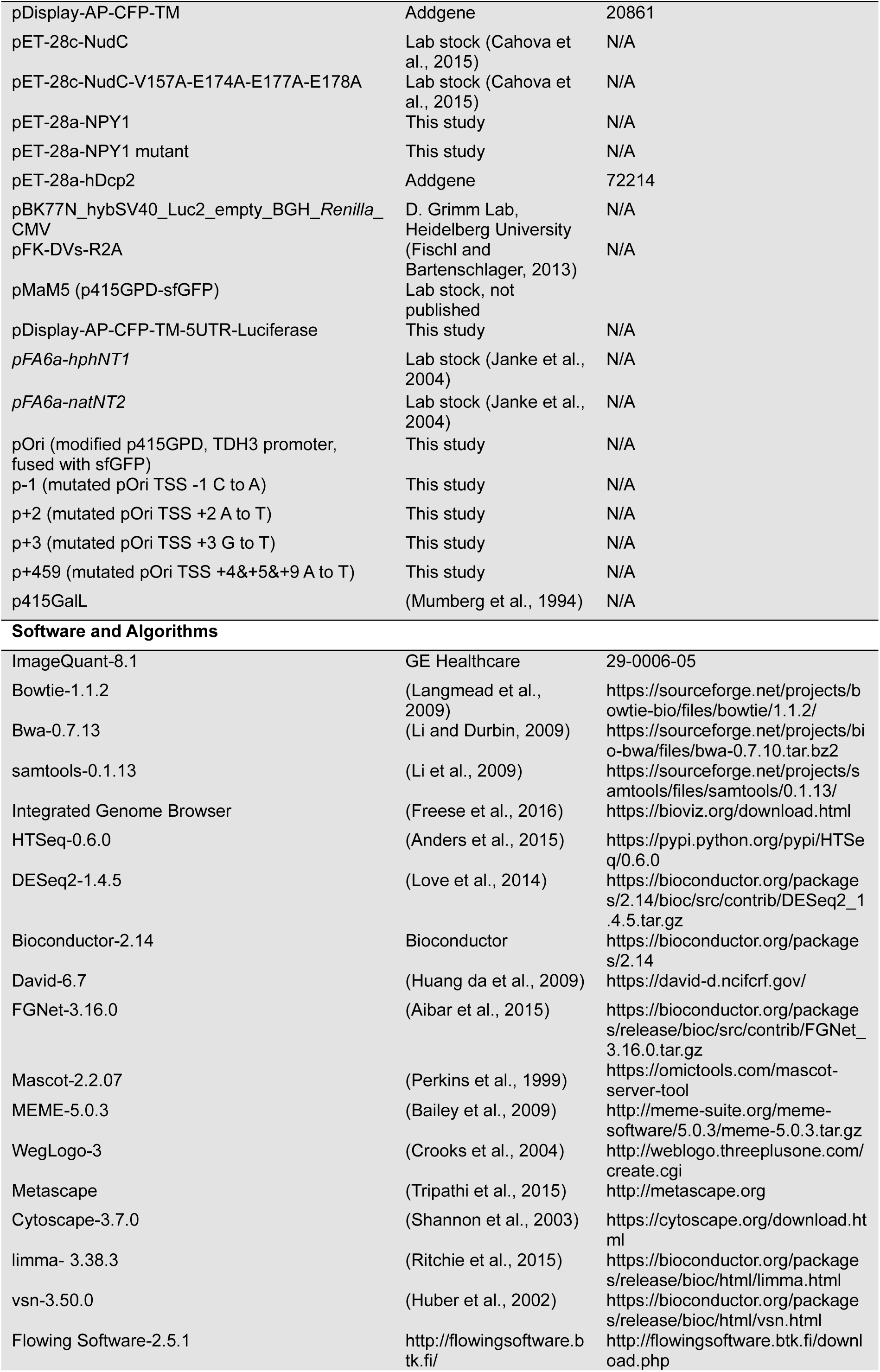
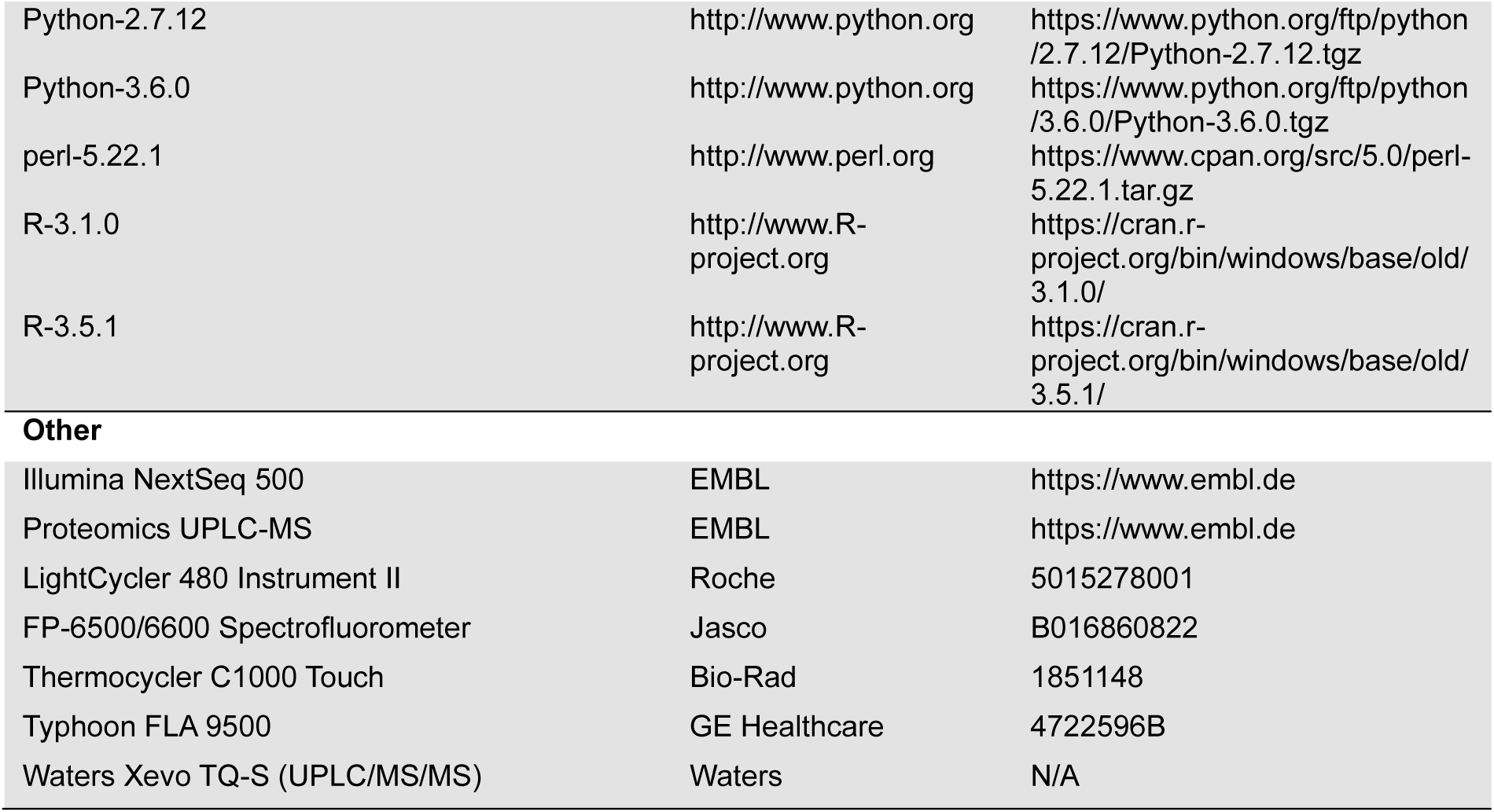

### LEAD CONTACT AND MATERIALS AVAILABILITY

Further information and requests for resources and reagents should be directed to and will be fulfilled by the lead contact, Andres Jäschke. All reagents generated in this study will be made available on request, but we may require a payment and/or a completed Materials Transfer Agreement if there is potential for commercial application.

### EXPERIMENTAL MODEL AND SUBJECT DETAILS

Unless otherwise stated, *S. cerevisiae* strains were grown in yeast extract/peptone/dextrose media (YPD). All strains used in this study except of YDK587-1 and its derivatives, YDK53-7, and C-SWAT mNeonGreen (mNG-I) strains were derivatives of the S288C strain BY4742 (*MATα his3Δ1 leu2Δ0 lys2Δ0 ura3Δ0*) and are listed in the key resource table. YDK53-7 and C-SWAT mNeonGreen (mNG-I) strains were derivatives of the S288C strain BY4741 (*MATa his3Δ1 leu2Δ0 met15Δ0 ura3Δ0*) and YDK587-1 was a derivate of the S288C strain ESM356-1 (*MATa ura3-52 leu2Δ1 his3Δ200 trp1Δ63*). Strains were cultivated in YPD medium at 30 °C according to standard protocols. Antibiotics were used at the following final concentrations: 200 μg/mL geneticin (G418), 300 μg/mL hygromycin B, 100 μg/mL nourseothricin, Gene deletions were performed using standard PCR-based recombination methods as described (Janke et al., 2004; Sikorski and Hieter, 1989), followed by PCR-based confirmation. Single, double and triple mutants were generated by mating and sporulation followed by random spore isolation. Plasmid transformations were performed using standard methods (Knop et al., 1999; Schiestl and Gietz, 1989) and transformants were selected on synthetic complete (SC) medium lacking leucine for p415 based plasmids.

### METHOD DETAILS

#### Fluorescence Microscopy and Colony Fluorescence Imaging

For fluorescent live cell imaging a DeltaVision Elite widefield fluorescence microscope (GE Healthcare) consisting of an inverted epifluorescence microscope (IX71; Olympus) equipped with a light-emitting diode light engine (seven-color InsightSSI module; GE Healthcare), an sCMOS camera (pco.edge 4.2; PCO), and a 60× 1.42 NA Plan Apochromat N oil immersion objective (Olympus) was used. Cells were inoculated in low fluorescence medium (synthetic complete (SC) medium prepared with yeast nitrogen base lacking folic acid and riboflavin; CYN6501, ForMedium) and grown to mid-log phase. Cells were immobilized for imaging in glass-bottomed 96-well plates (MGB096-1-2-LG-L; Matrical) using Concanavalin A and fluorescence filters for GFP, as described previously (Khmelinskii and Knop, 2014). For imaging the fluorescence of cell colonies and cell patches on plates, a custom made fluorescence illumination cabinet was used, equipped with filters for GFP.

#### Total RNA Isolation and Purification

Total RNA from yeast BY4742 strains was isolated by the hot phenol method (Collart and Oliviero, 2001) with minor changes. Cells were harvested (OD_600_ 0.8) from 0.5 L YPD medium, quickly frozen in liquid nitrogen, and stored at -80 °C until all samples were ready. Cell pellets were thawed on ice and washed with dH_2_O, resuspended in 8 mL TES solution (10 mM Tris-HCl (pH 7.5), 10 mM EDTA, 0.5% SDS) and were added to 8 mL phenol. The mixture was vortexed thoroughly for 1 min, incubated for 60 min at 65 °C with occasional shaking (50 seconds/10 min, 550 rpm), and placed on ice for 15-20 min. Then the samples were centrifuged at 14,000 g, 4 °C. The aqueous supernatant was subjected to phenol extraction, P/C/I purification, and chloroform purification. RNA was precipitated with 0.1 volumes 3 M NaOAc (pH 5.5) and 2.5 volumes ethanol at -20 °C, overnight. Precipitated RNA was dissolved in 2 mL dH_2_O. RNA concentration was determined by Nanodrop spectrometry and its integrity analyzed by 1.2% formaldehyde denaturing agarose gel electrophoresis. 1 mg RNA was treated with 100 U DNase I in 1X DNase I buffer (Roche) for 40 min at 37 °C. DNase I was removed by P/C/I extraction (twice), followed by ethanol precipitation of the RNA (−20 °C, overnight). The pellet was dissolved in 100 µL dH_2_O, the RNA concentration determined by Nanodrop and its integrity analyzed as before.

#### NAD captureSeq and Transcriptome Libraries Preparation

Standard, unfragmented NAD captureSeq: Total RNA was subjected to the NAD captureSeq protocol (Winz et al., 2017) with minor modifications, as outlined in the following. Total RNA, obtained in biological triplicates, was used as starting material for library preparation. More specifically, 100 µg total RNA was commonly supplemented with 5 ng NAD-RNAI (‘spike-in’ control) and treated with adenosine diphosphate-ribosylcyclase (ADPRC, Lab stock, 22.5 U), in the presence of 10 µL 4-pentyn-1-ol at 37 °C and for 60 min in 100 µL total reaction volume, including 50 mM Na-HEPES (pH 7.0), 5 mM MgCl_2_. The same reaction mixture, omitting the ADPRC enzyme, served as background control (negative control). The reactions were stopped by adding 100 µL dH_2_O and 200 µL P/C/I reagent. This was followed by performing P/C/I extraction twice, and three additional ether extractions. Subsequent copper-click reactions, capture by streptavidin bead, preadenylated-3’-adapter ligation, reverse transcription (RT), CTP-tailing, anchored-5’-adapter ligation and cDNA PCR amplification with barcode oligos were performed as described previously (Winz et al., 2017). Library quality control was conducted by Sanger sequencing. Specifically, the amplicon derived from the NAD-RNAI spike-in control as well as other cDNA amplicons were amplified using corresponding adapter-ligated sequences. Resulting cDNA libraries were then cloned into plasmid(s) for Sanger sequencing. After the successful pre-sequencing, PCR products were purified and size-selected within a range of 150-300 bp by 10% native polyacrylamide gel electrophoresis (PAGE). Absence of primer-dimers was ensured by Bioanalyzer 2100 (Agilent) analysis using the Agilent High Sensitivity DNA Kit. Concentrations were measured using the Qubit approach. Primer-dimer-free libraries were then multiplexed at a final concentration of ∼20 nM. Based on the cDNA length distribution, the library pools were either supplemented with 20% v3 Phix Control (Illumina) or custom Illumina sequencing primers, which bear three Gs at their 5’-end, preceded by the Illumina standard sequencing, to mitigate library imbalances before NextSeq 500 75 bp single-end (SE) sequencing.

Fragmented NAD captureSeq: Biological triplicates of total RNA (gDNA-free) were used as library starting material. 100 µg total RNA, supplemented with 5 ng NAD-RNAI (as optional spike-in control), were randomly sheared in a 65 µL reaction volume that contained 32.5 µL alkaline fragmentation solution (2 mM EDTA, 10 mM Na_2_CO_3_, 90 mM NaHCO_3_, pH 9.3) at 94 °C (5 min for wild type strain and 20 min for *npy1*Δ strain to approach similar fragment size). Sheared RNA fragments were visualized on a 1.2% formaldehyde denaturing agarose gel. Next, the sheared RNA was precipitated, in a total volume of 200 µL dH_2_O, by addition of 600 µL ethanol, 20 µL 3 M NaOAc (pH 5.5), and 1 µL glycogen at -20 °C, overnight. The precipitated RNA was washed with 75% ethanol and subsequently dissolved in 50 µL dH_2_O. An equivalent of 100 µg sheared RNA was then treated with 100 U T4 PNK, along with 0.1 mM ATP, 100 mM imidazole-HCl (pH 6.0), 10 mM MgCl_2_, 10 mM β-mercaptoethanol and 20 µg/ml RNase-free BSA in a total volume of 200 µL at 37 °C for 5.5 h. RNA extraction was performed twice, employing the P/C/I approach, and followed by triple ether extraction and ethanol precipitation. Precipitated RNA was the washed again, using 75% ethanol, and ultimately dissolved in 20 µL dH_2_O, yielding the library input for the standard NAD captureSeq protocol (Winz et al., 2017). PCR products were size-selected within a range of 150 to 300 bp (referred to as ‘small fragmented NAD captureSeq library’) and of 300 to 500 bp (referred to as ‘large fragmented NAD captureSeq library’), enabled by 10% native polyacrylamide gel electrophoresis (PAGE). Bioanalyzer quality control, library multiplexing and the overall sequencing strategy were executed in a similar manner, as done for the unfragmented NAD captureSeq. Again, the NextSeq 500 75bp SE approach was chosen for sequencing.

Transcriptome libraries: Biological triplicates of total RNA (gDNA-free) served as library input material. 1 µg total RNA was subjected to ribosomal RNA depletion (rRNA) by Ribo-Zero rRNA Removal Kit (yeast). rRNA-depleted RNA was randomly sheared in 10 µL dH_2_O, at 94 °C 10 min. Fragmented RNA was processed using the NEBNext Ultra II Directional RNA Library Prep Kit for Illumina, following the instructions of the manufacturer. cDNA was barcoded by PCR amplification using NEBNext Multiplex Oligos for Illumina. Further cDNA size selection in the range of 300 to 500 bp was performed, employing the Agencourt RNAClean XP kit. Primer-depleted cDNA was examined by Bioanalyzer and the concentration was measured by Qubit. Multiplexed libraries were sequenced by NextSeq 500 75bp SE.

#### NGS Analysis

Unfragmented NAD captureSeq Analysis: Original reads were demultiplexed, based on the PCR barcode, not allowing for mismatches and subjected to standard quality control procedures. Further 5’-end leading (G/N)_n_ and 3’-end adapters (C NNNNNN AGATCG) were trimmed (minimum length 12 nt) by in-house scripts. Reads that mapped to the spike-in internal standard (IS) RNAI sequence (bowtie - v 2, version 1.1.2) were then counted first. IS unmapped reads were subsequently classified as ‘small RNA reads’ (12-17 nt) or ‘normal RNA reads’ (>18 nt).

Normal RNA reads were mapped to the reference genome *S. cerevisae* BY4742 strain (BY4742_Toronto_2012, SGD) (bowtie -v 2). Normal RNA reads that mapped to rRNA genes or tRNA genes were separated and counted individually. Remaining reads that could not be mapped to rRNA genes were remapped to the yeast reference genome (bowtie –best --strata -M 1 -m 20 -v 2). The .sam files were then converted to .bam files and sorted (samtools, version 0.1.13). The .bam files were used to generate .wig files, which could be normalized by Reads per million mapped reads (RPM), for single nucleotide resolution-based analysis. Sorted .sam files were further filtered by strand-specific selection and used for three different types of analysis:

On the one-hand side, the filter-passed reads from sorted .sam files were used for linear trend tests (Agresti, 2014) to perform an alternative transcription starting site (TSS) study. Briefly, the position of the first nucleotide of the mapped reads within the range of the 5’ UTR (Nagalakshmi et al., 2008) with additional 50 nt upstream extension from each gene was collected. A sliding window of 18 nt was applied for searching for TSS clusters from the highest density one to the lowest one. Then a 2 x n table was made, whereby “n” is the number of unique TSS clusters found in the sample group (S, +ADPRC) and/or the negative control group (N, -ADPRC). Values in the table represent the number of reads for these clusters in the S or N group (normalized by the proportion of the respective cluster relative to the sum of reads for all clusters), and are sorted from the shortest (left) to the longest UTR (right). For covariance analysis, the top row (S) was assigned a weight value of 2, while the bottom row (N) was assigned 1 Column (*Y*, “n” columns) weight was set to be the 5’ UTR length of each cluster. The Pearson correlation r was calculated by *cov (X,Y)*/ *(δ*_*x*_ *\*δ*_*Y*_). The operator M was calculated based on the equation *M*^*2*^*= (n-1) r*^*2*^ (n≥30), following the Chi square distribution (freedom 1). The corresponding p value was used for FDR calculation, employing the Benjamini-Hochberg method.

On the other side, the filter-passed reads were counted for RNA hits (htseq-count -m intersection-nonempty, version 0.6.0) based on the annotation .gff3 file (BY4742_Toronto_2012, SGD, mRNA regions were annotated by their 5’ UTR region (−120 to +65, translation starting site (TLS) referred as ‘0’, described in Figure S1). Raw hits were analyzed by DESeq2 (version 1.4.5, relevant package from Bioconductor version 2.14) for NAD-RNA enrichment statistics. Meanwhile, raw hits were normalized by transcripts per kilobase million (TPM) for NAD-ratio calculation. Simply, the relative total number of NAD-RNA (k) was the signal intensity from the UPLC/MS measurement of total RNA. The distribution of all-cap-RNA in total RNA was simulated by their TPM in transcriptome data. The distribution of NAD-RNA was simulated by their TPM in NAD captureSeq data. The relative individual NAD-ratio was calculated by k*TPM_NAD-RNA_/TPM_all-cap-RNA_.

Additionally, the filter-passed reads which mapped to 5’ UTRs of mRNAs were collected for “sharpA” promoter motif analysis. This analysis included the genome-mapped position of the first base of reads referenced to the TSS site. For each individual mRNA, either the unique most abundant mapped position of the first base of reads or one of most abundant mapped positions (several ones with equal abundance) was defined as TSS (reference as +1) but this position was annotated as ‘A’ in the genome. The sequence flanking the TSS from the -10 to +10 position, within the reference genome, was further analyzed for a consensus motif. The sharp value was defined as the ratio between the number of nucleotides accumulating at the +1 position and that at the -1 position, revealed by the generated .wig file. The sharp A feature was screened, based on the premise that the TSS constitutes an ‘A’ and possesses a sharp value bigger than 4.

For the small RNA reads group, reads were purged by additional genome mapping (bowtie -v 0). Next, small RNA clusters mapping to mRNA 5’ UTRs were assembled together (60% sequence similarity, same strand direction, and max fold copy number difference is 50). Differential abundance of small RNA reads was calculated by the enrichment of reads in S group to N groups. Fragmented NAD captureSeq was analyzed in the same way, as done for the unfragmented NAD captureSeq library.

#### Transcriptome Analysis

Raw reads were demultiplexed and quality control was performed using the same parameters, already employed for unfragmented NAD captureSeq analysis. Reads with 5’- / 3’-end adapters were properly trimmed, leveraging an in-house script. Then the sequence of raw reads was converted into its reverse complement. Reads were then mapped to the yeast reference genome (bowtie, -v 2). rRNA genes and tRNA genes reads that mapped independently, as well as rRNA-free hits counting (same annotation region on 5’ UTR of mRNA) were treated in a similar way as done in the NAD captureSeq analysis, described above. Also, .wig files and differential expression statistics were generated and performed according to the same procedure.

Function clusters were generated by David (version 6.7) and FGNet (version 3.16.0). Enriched promoter motifs were analyzed by MEME (version 5.0.3) and WebLogo (version 3). Meta-analysis of function clusters and pathways was enabled by Metascape (metascape.org) and cytoscape (version 3.7.0).

#### Proteomics Sample Preparation and TMT Labeling

Yeast cells were cultured in 100 mL YPD medium and collected at OD_600_ 0.8. The cells (biological triplicates) were washed twice with ice-cold dH_2_O then twice with ice-cold PBS. The pelleted cells were then lysed by passing them twice through a French press (∼0.69 kbar) in 2 mL protein lysis buffer (50 mM Tris-HCl (pH 7.5), 100 mM NaCl, 1 mM EDTA, 1 mM PMSF, 1 µg/mL leupeptin, 1 µg/mL pepstatin A). The lysate was centrifuged at 25,000 g, 4 °C, 20 min. The supernatant was collected and subsequently flash-frozen in liquid nitrogen and stored at -80 °C until all samples were ready. The protein concentration was measured by BCA assay. Reduction of disulfide bonds in cysteine-containing proteins was performed using 10 mM dithiothreitol (56 °C, 30 min, in 50 mM HEPES, pH 8.5). Reduced cysteines were alkylated with 20 mM 2-chloroacetamide (room temperature, in the dark, 30 min, 50 mM HEPES, pH 8.5). Samples were prepared following the SP3 protocol (Hughes et al., 2014) and subsequently trypsin (sequencing grade) was added in an enzyme to protein ratio of 1:50 for overnight digestion at 37 °C. Peptides were labelled using the TMT10plex (Werner et al., 2014) Isobaric Label Reagent, according the manufacturer’s instructions. For further sample clean up, an OASIS HLB µElution Plate (Waters) was used. Offline high pH reverse phase fractionation was carried out on an Agilent 1200 Infinity high-performance liquid chromatography system, equipped with a Gemini C18 column (3 μm, 110 Å, 100 × 1.0 mm, Phenomenex) (Reichel et al., 2016).

#### Proteomics Mass Spectrometry Data Acquisition and Analysis

An UltiMate 3000 RSLC nano LC system (Dionex) was fitted with a trapping cartridge (µ-Precolumn C18 PepMap 100, 5 µm, 300 µm i.d. × 5 mm, 100 Å) and an analytical column (nanoEase M/Z HSS T3 column 75 µm × 250 mm C18, 1.8 µm, 100 Å, Waters). Trapping was carried out with a constant flow of solvent A (0.1% formic acid in water) at 30 µL/min onto the trapping column for 6 minutes. Subsequently, peptides were eluted via the analytical column with a constant flow of 0.3 µL/min with an increasing percentage of solvent B (0.1% formic acid in acetonitrile) from 2% to 4% in 4 min, from 4% to 8% in 2 min, followed by 8% to 28% for a further 96 min, and finally from 28% to 40% in another 10 min. The outlet of the analytical column was coupled directly to a QExactive plus (Thermo Scientific) mass spectrometer using the proxeon nanoflow source in positive ion mode.

The peptides were introduced into the QExactive plus via a Pico-Tip Emitter 360 µm OD x 20 µm ID; 10 µm tip (New Objective) and an applied spray voltage of 2.3 kV. The capillary temperature was set to 320 °C. A full mass scan was acquired with a mass range from 350 to 1400 m/z in profile mode in the FT with a resolution of 70000. The filling time was set to the maximum of 100 ms with a limitation of 3×10^6^ ions. Data-dependent acquisition (DDA) was performed with the resolution of the Orbitrap set to 35000, with a fill time of 120 ms and a limitation of 2 × 10^5^ ions. A normalized collision energy of 32 was applied. A loop count of 10 with count 1 was used and a minimum AGC trigger of 2e2 was set. A dynamic exclusion time of 30 s was used. The peptide match algorithm was set to ‘preferred’ and charge exclusion ‘unassigned’, charge states 1, 5 - 8 were excluded. MS2 data was acquired in profile mode.

IsobarQuant (Franken et al., 2015) and Mascot (v2.2.07) were used to process the acquired data, which was searched against a Uniprot *S. cerevisiae* proteome database (UP000002311) containing common contaminants and reversed sequences. The following modifications were included into the search parameters: Carbamidomethyl (C) and TMT10 (K) (fixed modification), Acetyl (N-term), Oxidation (M) and TMT10 (N-term) (variable modifications). For the full scan (MS1) a mass error tolerance of 10 ppm and for MS/MS (MS2) spectra of 0.02 Da was set. Further parameters were set: Trypsin as protease with an allowance of maximum two missed cleavages: a minimum peptide length of seven amino acids; at least two unique peptides were required for a protein identification. The false discovery rate on peptide and protein level was set to 0.01.

The protein.txt – output file of IsobarQuant was analyzed using an R script. As a quality control filter, only proteins which were quantified with at least 2 unique peptides were used (2256 out of 6049 proteins remained). The signal_sum columns were annotated according to the experimental conditions. Batch-effects were removed with the limma (v3.38.3) package and subsequently the data was normalized using vsn (v3.50.0). limma was used again to test for differentially expressed genes between wild type and *npy1*Δ. Proteins were annotated as a hit with a fold-change bigger 50% and a false discovery rate smaller 5% and as a candidate with a fold change bigger 40% and a false discovery rate smaller 20%.

#### RNA Pull-down and UPLC-MS Analysis

250 µL Streptavidin Sepharose High Performance beads were loaded on Mobicol Classic columns. The column was washed three times with 1X PBS buffer, then five times 25 µL 25 µM biotin-DNA probe (Biomers, Table S1) were added sequentially. The mixture was incubated at 25 °C for 10 min. Next, the column was washed two times with 300 µL 1X PBS, followed by equilibration in 300 µL pull-down buffer (10 mM Tris-HCl (pH 7.8), 0.9 M tetramethylammoniumchloride, 0.1 M EDTA (pH 8.0)). 200-500 µg total RNA (gDNA-free) was added into the column and incubated at 65 °C for 10 min and then rotated (Tube Rotator, VMR) at 20 °C for 25 min. Next, the column was washed six times with 200 µL dH_2_O, in order to remove unspecifically binding RNAs. RNA was eluted by adding four times 200 µL 2 mM EDTA (75 °C, pre-heated) under 10 s/min shaking (350 rpm) at 75 °C for 10 min. The eluate was precipitated with 0.5 M ammonium acetate (pH 5.5) and 50% isopropanol. Precipitated RNA was dissolved in dH_2_O for UPLC/MS analysis.

To determine the amount of NAD that is covalently linked to RNA, the RNA samples were washed three times with 400 μL 8.3 M urea, one time with dH_2_O, two times with 4.15 M urea and again four times with dH_2_O in Amicon Ultra-0.5 mL Centrifugal Filter Units 10 kDa to remove non-covalently bound cellular NAD. The recovered RNA was subsequently concentrated. The pull-down RNA samples or 10 µg urea-washed total RNA samples were treated with 10 µM NudC in the presence of 10 mM MgCl_2_, 0.6 ng/mL d4-riboside nicotinamide as internal standard, and 0.05 U/µL alkaline phosphatase at 37 °C for 2 h. The same reaction with a catalytically inactive NudC mutant served as background reference. The reactions were filtered through Amicon Ultra-0.5 mL Centrifugal Filter Units 10 kDa to remove the enzymes and washed additionally four times with 200 µL dH_2_O. The flow-through contained the cleaved nicotinamide-riboside (NR), resulting from the NudC-treatment of NAD-RNA and d4-NR as internal standard. It was collected and dried under vacuum. The amount of NR was determined by UPLC-MS/MS, and reflects the exact same amout of original NAD-RNA in the digested sample. The employed UPLC-MS/MS setup contained a triple stage quadrupole mass-spectrometer (Waters, Xevo TQ-S) system coupled with an Acquity UPLC system. A BEH Amide column (1.7 μm, 2.1×50 mm) was used with an eluent A (0.01% (v/v) aqueous formic acid with 0.05% ammonia and 5% acetonitrile) and B (acetonitrile included 0.01% formic acid), at a flowrate of 0.8 mL/min. The gradient started to change from 12.5% A /87.5%B to 95% A/ 5% B within 1.8 min. The ratio was changed back to starting conditions within the following 1.0 min. The column was had been pre-equilibrated under starting conditions for 1.0 min. Electrospray ionization (ESI) was performed with a 1500 V capillary voltage, 11 V cone voltage, 150 °C source temperature, 200 °C desolvation temperature, 150 L/h cone nitrogen gas flow, and 800 L/h desolvation gas flow (N_2_). The Xevo TQ-S was automatically tuned to d4-nicotinamide riboside and nicotinamide riboside using the MassLynx V4.1 system software (Waters) with the IntelliStart standard procedures. Multiple reaction monitoring (MRM) measurements were conducted, using collision gas (argon, 0.15 ml/min) for collision-induced decomposition (CID) and MS/MS transitions were monitored in the positive ion mode (N-ribosylnicotinamide: m/z 254.94 to 122.81, d4-N-ribosylnicotinamide m/z 258.94 to 126.81, 20 V, 50 mS dwell time for each mass transition).

#### Flow Cytometry Data Acquisition and Analysis

Cells, in biological triplicates, were grown in low fluorescence synthetic complete medium lacking leucine to mid log phase. Flow cytometry (FCM) of yeast cells expressing GFP from p415 based plasmids was performed on a BD FACSCanto™II (BD Bioscience) equipped with a 488-nm laser and a combination of 502-nm long-pass and 530/30-nm band pass emission filters for GFP detection. Total 300000 events were measured for data analysis using Flowing Software. Events of single cells were isolated. Then the events of fluorescence background were removed. Fluorescence intensity of remained events was analyzed.

#### Gel Electrophoresis

10% native PAGE was utilized to size select cDNA for NGS. Briefly, 10% acrylamide/Bis solution (19:1), 0.1% APS (w/v) and 0.1% TEMED (v/v) along with 1X Tris-borate-EDTA (TBE) in 50 mL volume were mixed and poured between glass plates (19 cm × 27cm). Gel mixtures were polymerized at room temperature for 45 min. The electrophoresis conducted at a stable 27 mA current for 2.5 h. The gel was then stained with SYBR Gold (Thermo Scientific) in 1X TBE buffer, 5 min. The signal intensities were read-out by scanning the gel at 400 V, 50 or 100 µm resolution using a Typhoon FLA 9500. The printout picture of the gel (in its original size) was used for excision of desired size ranges within the corresponding gel lanes. APB gel electrophoresis was utilized to separate NAD-RNA, m^7^G-RNA, and p/ppp-RNA from each other. Briefly, 0.5% (w/v) APB (Lab stock), 10% acrylamide/Bis solution (19:1), 0.1% (w/v) APS, and 0.1% (v/v) TEMED with 2.5 X Tris-acetate-EDTA (TAE) buffer in 50 mL volume were mixed and poured between glass plates (Bio-Rad). Gels were run at a stable current (15-29 mA per gel) in 1X TAE buffer. Gels were then stained with SYBR Gold or gels, containing ^32^P-labeled nucleic acids, were exposed to storage phosphor screens (GE Healthcare) and visualized using a Typhoon FLA 9500. Signal quantification was performed using the ImageQuant software (GE Healthcare).

#### *In vitro* Transcription and NAD/ppp/p/m^7^G-capped RNA Preparation

Radio-labeled RNA: 400 nM double-stranded DNA (dsDNA) served as a template in a transcription buffer containing 40 mM Tris-HCl pH 7.9, 1 mM spermidine, 22 mM MgCl_2_, 0.01% Triton X-100, 10 mM dithiothreitol (DTT), 5% DMSO, along with 60 µCi α-^32^P-ATP, 10 µCi α-^32^P-CTP, 4 mM each CTP/GTP/UTP, 2 mM ATP, 6 mM NAD (NAD-RNA) or without NAD (ppp-RNA), 0.7 µM T7 polymerase (self-prepared) within a total reaction volume of 100 µL. The *in vitro* transcription reaction was incubated at 37 °C for 3 h. Subsequently, 10 U DNase I were added and the reaction incubated for an additional 30 min at 37 °C. ppp-RNA was purified by 10% denaturing PAGE, as described above. NAD-RNA was first purified by 10% denaturing PAGE, followed by a 10% PAGE, supplemented with 0.5% APB, in order to remove ppp-RNA. For p-RNA preparation, 1 µg ppp-RNA was treated with 20 U RNA 5’-polyphosphatase and 1X Reaction Buffer (Epicentre) in a reaction volume of 20 µL for 1 h at 37 °C. The desired p-RNA species was then obtained by performing P/C/I extraction twice, followed by ethanol precipitation, as described. To generate m^7^G-RNA, ∼1.2 µM denatured ppp-RNA (65°C, 5 min) was first added to a mixture of 1 X Scriptcap capping buffer, 1 mM GTP, fresh 0.1 mM S-adenosyl methionine (SAM), and 1 U/µL Script Guard RNase Inhibitor. The mixture was then supplemented with U/ µL Scriptcap Capping Enzyme, in a final reaction volume of 50 µL and incubated for 50 min at 37 °C. The modified RNA reaction was then purified by 10% APB-PAGE as described above, in order to separate uncapped, ppp-RNA, and m^7^G-capped RNA.

Luciferase mRNA: 140 nM (∼6 µg) linear dsDNA template was added into the same transcription buffer as described above, along with 4 mM each CTP/GTP/UTP, 2 mM ATP, 6 mM NAD (NAD-mRNA) or without NAD (ppp-mRNA), 0.7 µM T7 polymerase (self-prepared) with a total reaction volume of 100 µL. The reaction mixture was incubated for 3 h at 37 °C. Similarly, 10 U DNase I were added subsequently to the reaction and incubated for an additional 30 min at 37 °C. RNA integrity was examined by visualization of the reaction products on a 1% formaldehyde denaturing agarose gel. RNA was purified by performing P/C/I extraction twice and precipitated with ethanol. Precipitated RNA was loaded onto Amicon Ultra-0.5 mL Centrifugal Filter Units 10 kDa and washed four times with 400 µL dH_2_O to remove free NTPs and small molecules. Retained RNA was collected and again precipitated with ethanol. Recovered RNA was then washed twice with 75% ethanol and ultimately dissolved in 20 µL dH_2_O. For further isolation of 5’-NAD-modified RNA, 5 µg partially purified NAD-RNA was treated with 5 U RNA 5’-polyphosphatase in1X Reaction Buffer (Epicentre) with an overall volume of 10 µL for 40 min at 37 °C. Remaining NAD-capped RNA was then P/C/I extracted twice and ethanol precipitated. Precipitated mRNAs were further treated with 2 U Xrn-1 in 1X NEBuffer 3 (NEB) reaching a total reaction volume of 40 µL. For p-mRNA preparation, 5 µg ppp-mRNA was treated with 10 U RNA 5’-polyphosphatase in 1X Reaction Buffer (Epicentre) and final volume of 20 µL for 40 min at 37 °C. For m^7^G-mRNA preparation, 1 µM denatured ppp-mRNA (65°C, 5 min) was mixed with 1X capping buffer (NEB), 0.5 mM GTP, fresh 0.1 mM SAM, and 0.5 U/µL Vaccinia Capping Enzyme in 20 µL volume, incubate at 37 °C, 40 min. Processed RNA was purified twice by P/C/I extraction and precipitated with ethanol. All mRNA raw concentrations were measured by nanodrop, followed by quantitative reverse transcription PCR (qRT-PCR) to determine the relative abundance of the mRNA middle region, as well as of the 3’-end region. Final concentration of mRNAs was adjusted accordingly.

#### Plasmid Construction

Npy1: the open reading frame (ORF) of *NPY1 gene* was PCR-amplified from *S. cerevisiae* (BY4742 strain) gDNA, and NdeI and BamHI restriction sites introduced using the respective primers (Fw_NPY1 and Rev_NPY1, Table S1). The PCR products were then purified using the QIAquick PCR Purification Kit (QIAGEN) and digested with NdeI and BamHI-HF. The digested amplicons were subsequently ligated into a NdeI- and BamHI-digested pET-28a (+) plasmid (Novagen) by T4 DNA ligase, which is subsequently referred to as pET-28a-NPY1.

Mutant Npy1 (E276Q): For site-directed mutagenesis, primers encoding the desired alterations as mismatches (Fw_NPY1m and Rev_NPY1m, Table S1) were used to modify the plasmid pET-28a-NPY1, under standard PCR conditions, to obtain pET-28a-NPY1(E276Q) mutant. The plasmids pET-28c-NudC, pET-28c-NudC-V157A-E174A-E177A-E178A (NudC-M1) were available as lab-prepared stock.

mRNA for *in vitro* translation: The cloning procedure to obtain template mRNA sequences of interest had three distinct steps. Firstly, the 5’ UTR and 22 nt of the coding sequence (CDS) of the gene were PCR-amplified from *S. cerevisiae* (BY4742 strain) gDNA using the corresponding primers (Table S1). Secondly, the firefly luciferase sequence (1653 bp, same as from pGL4.10 [Luc2] Vector (Promega)) and the *Renilla reniformis* luciferase sequence (936 bp, same as from pRL-null Vector (Promega)) were cloned from the plasmid pBK77N_hybSV40_Luc2_empty_BGH_*Renilla*_CMV and pFK-DVs-R2A, respectively, using dedicated primer pairs (Fw_Luc2, Rev_luc2, Fw_*Renilla* and Rev_*Renilla* primers; Table S1). Thirdly, the T7 promoter sequence containing a HindIII site was fused with 5’-terminal mRNA sequence, firefly luciferase sequence, and a poly(A)_30_ tail sequence by corresponding, employing appropriate primers (Fw_5UTR, bridge_region and Rev_Luc2_polyA; Table S1) with two rounds of PCR amplification. PCR products were purified by 0.8% agarose gel electrophoresis. Afterwards, 0.02 µM PCR amplicon was phosphorylated by 15 U T4 PNK with 1X T4 DNA Ligase Buffer (Thermo Scientific) in an overall volume of 20 µL. 540 fmol linear vector pDisplay-AP-CFP-TM was digested by EcoRV-HF, then dephosphorylated by 1 U Fast AP Thermosensitive Alkaline Phosphatase (Thermo Scientific) in 1X Fast AP Buffer in a total reaction volume of 20 µL. The 15 fmol phosphorylated PCR product was subsequently ligated with 5 fmol dephosphorylated linear vector by 30 U T4 DNA ligase in a 20 µL reaction at 20°C for 1 h.

YAAG promoter: The plasmid backbone used here was a derivative of the p415GPD vector (Mumberg et al., 1995). The *TDH3* gene promoter and 15 nt of the CDS were obtained from *S. cerevisiae* gDNA by standard PCR amplification, using corresponding primers (F_TDH3_5UTR_15ntCDS_SacI and R_TDH3_5UTR_15ntCDS_XbaI; Table S1). The p415GPD plasmid and the generated PCR products were digested with SacI-HF and XbaI and subsequently ligated. Next, sequence superfolder GFP (sfGFP) was amplified from pMaM5 plasmid under standard PCR conditions, using appropriate primers (F_sfGFP_BamHI and R_sfGFP_HindIII; Table S1). These sfGFP-encoding amplicons and the partially assembled genetic constructs, described above, were again digested with BamHI-HF and HindIII-HF and ligated together as the pOri plasmid. Plasmids carrying mutations at the positions p-1, p+2, p+3, p+459 were generated by PCR using dedicated primer pairs, separately (Mutagenesis_R, Mutagensis_p-1A_F, Mutagensis_p+2T_F, Mutagensis_p+3T_F, and Mutagensis_p+459T_F, Table S1). Correct insert sequences of all plasmids were further confirmed by Sanger sequencing.

#### gDNA Extraction

The gDNA isolation procedure was based on a published method (Harju et al., 2004). 1.5 mL of the yeast overnight culture (BY4742 strain) were pelleted and resuspended in 200 µL lysis buffer, containing 2% Triton X-100, 1% SDS, 100 mM NaCl, 10 mM Tris-HCl (pH 8.0), 1 mM EDTA (pH 8.0). Resuspended cells were kept at -80 °C for 15 min and then directly heated to 95 °C for 1 min. Samples were subjected to two additional freeze-thaw cycles, as described. The mixture was then vortexed for 30 s. Subsequently, 200 µL chloroform was added and the mixture vortexed for 2 min at RT before centrifugation. Then the aqueous layer from the centrifugation was transferred into 400 µL ethanol (ice-cold). The solution was incubated at RT for 5 min. Then the solution was centrifuged at 20,000 g for 10 min at RT. The supernatant was collected and dried under vacuum. The gDNA pellet was then resuspended in 20 µL TE buffer (10 mM Tris-HCl (pH 8.0), 1 mM EDTA (pH 8.0)).

#### Cell-free Extract Preparation and *in vitro* Translation

Preparation of cell-free extracts for *in vitro* translation was performed as previously described (Wu and Sachs, 2014) with minor modifications. Cell pellets were obtained from 0.5 L YPD yeast (BY4742 strain) culture (OD_600_≈15). The cell pellet was washed three times with 50 mL ice-cold Breaking Basic Buffer (30 mM HEPES-KOH (pH 7.6), 100 mM KOAc (pH7.0), 3 mM Mg(OAc)_2_ (pH 7.0), 2 mM DTT), supplemented with 8.5% (w/v) mannitol. Then, the wet weight of the cells was determined. The cells were subsequently resuspended in Breaking Basic Buffer with 8.5% mannitol and 0.5 mM PMSF (Sigma-Aldrich, dissolved in isopropanol), under addition of 1.5 mL per 1 g wet weight. The solution was supplemented with ice-cold, sterile ∼0.5 mm glass beads, whereby the equivalent of six times of the cell wet weight was added. The cells were raptured by manual shaking (70 cm hand path, 2-5 Hz), in overall five rounds, each consisting of 1 min shaking interrupted by 1 min cooling on ice. Glass beads were removed by low speed centrifugation (1000 g, 2 min, 4 °C), then the cleared supernatant was obtained by two consecutive rounds of centrifugation (29000 g, 20 min, 4 °C). The solution was filtered through a 0.45 µm filter and then 2 mL of the supernatant were subjected to FPLC runs, outlined below. HiPrep 26/10 Desalting Columns (containing Sephadex G-25 Fine resin) were pre-equilibrated with 100 mL Breaking Basic Buffer, supplemented with 0.5 mM PMSF. The injected sample was resolved by the indicated column matrix running on a FPLC system (flow rate 1 mL/min, mL collected fractions) employing the same equilibration buffer. Absorption values of each fraction were determined at 260 nm (A_260_) and appropriate fractions, exceeding 75% of the highest A_260_ value, were pooled together. Next,1 mM CaCl_2_ and 50 U/mL micrococcal nuclease were added to the pooled fractions, followed by incubation at 26 °C for 15 min. The reaction was stopped by adding EGTA to a final concentration 2.5 mM. Aliquots of 100 µL each were then flash-frozen in liquid nitrogen and stored at -80 °C.

*In vitro* translation: Master Translation Solution was freshly prepared by mixing 25 mM HEPES-KOH (pH 7.6), 1.25 mM ATP, 0.125 mM GTP, 0.15 U/ µL creatine phosphokinase, 2.5 mM DTT, 125 mM KOAc, 5 mM MgOAc (pH 7.0), 25 µM Amino Acid Mixtures, 1 U/µL RNasin Ribonuclease Inhibitors, 80 nM m^7^G-capped *Renilla* mRNA, and dH_2_O in a final volume 80 µL. The solution was gently mixed and 4 µL of the Master Translation Solution aliquoted to individual PCR tubes. Then, 1 µL mRNA (200 ng or accordingly) and 5 µL of the cell-free extract were added to thus prepared reaction volumes. The ‘ready-for-translation’ solution was again mixed gently and incubated at 26 °C for 30 min. Next, 90 µL dH_2_O were added to each *in vitro* translation reaction. 75 µL of this dilution were then used to conduct a firefly luciferase activity assay (Bright-Glo Luciferase Assay System). The remaining 25 µL, with additional 50 µL of dH_2_O, were subjected to a *Renilla* luciferase activity assay (*Renilla*-Glo Luciferase Assay System), executed according to the manufacturing instructions. Emitted luminescence was read out using a TECAN plate reader.

#### Cellular NAD Quantification

The general experimental procedure was based on the manufacturer’s protocol (NAD/NADH Quantification Kit) with minor modification. Yeast cells were cultured in 50 mL YPD medium in biological quadruplicates (n=4). 1 mL of these cultures, at an OD_600_ of ∼ 0.8, were pelleted by the centrifugation at 4000 g, 1 min, 4 °C. The cell pellets were washed four times with 1 mL ice-cold PBS and resuspended in 1 mL NADH/NAD Extraction Buffer. 400 µL of the resuspension were subjected to three freeze-thaw cycles, each consisting of 10 min on dry ice, alternating with 10 min thawing at room temperature. The homogenized cell lysates were vortexed for 10 s and centrifuged at 14000 g, 4 °C, 10 min. 20 µL of each supernatant were aliquoted for subsequent lysate RNA quantification. The remaining solution was applied onto a 10 kDa spin-filter to remove proteins larger than 10 kDa, by centrifugation at 14000 g, 4 °C, 10 min. The flow-throughs were collected and NAD/NADH quantification was conducted, as stated by the manufacturer. The determined NAD amounts were normalized by the amount of measured lysate RNA.

#### Protein Expression and Purification

*ADPRC*: The *Pichia pastoris* GS115 pPICZαA/CYCLASE-2 strain was a gift from H.C. Lee. The protein expression and purification was performed as described previously with minor modifications (Munshi and Lee, 1997). First, *P. pastoris* was cultured grown on YPD agar plates, containing 100 µg/mL Zeocin. Then, single colonies were used to inoculate 10 mL liquid YPD medium, and subsequently cultured in a final volume of 500 mL YPD medium, until the optical density of the cells reached its plateau phase at about 50 mg cell pellet per mL medium. Next, the YPD medium was replaced by 500 mL BMMY medium (1% yeast extract, 2% peptone, 100 mM K_2_HPO_4_/KH_2_PO_4_ (pH 6.0), 1.34% (w/v) yeast nitrogen base with ammonium sulfate without amino acids, 0.4% mg/L biotin, 0.5% (v/v) methanol). Following the exchange of growth medium and after additional 24 h, as well as 48 h, of culturing the yeast in BMMY medium, 2.25 mL methanol were added. The supernatant of thus treated cultures was then collected, after overall 72 h, by centrifugation at 4 °C, 1000g, 10 min. The 500 mL of supernatant were filtered through a 0.45 µm filter and then concentrated employing Amicon Ultra-15 mL Centrifugal Filter Units 10 kDa, ultimately yielding 25 mL of concentrate. Contained proteins were dialyzed overnight against 1 L dialysis buffer, containing 50 mM NaOAc (pH 5.0), using a 5 kDa cut-off membrane (Carl Roth). Dialysis buffer was exchanged regularly with fresh 1 L dialysis buffer. Dialyzed protein solution was then loaded on 2X HiTrap SP HP 1mL columns at a flowrate of 1 mL/min on an FPLC system (Bio-Rad). ADRPC was eluted by applying a salt gradient from 50 mM NaOAc (pH 4.0) to 50 mM NaOAc (pH 4.0), 1M NaCl at 0.75 mL/min. The fractions that contained the ADPRC band upon SDS-PAGE analysis were pooled. The cyclase activity was determined conducting an NGD fluorometric assay. Briefly, ADPRC was subjected to serial dilution ranging from 0.2 ng/µL to 1.2 ng/µL, with constant amount of 60 µM NGD in HEPES Buffer (50 mM HEPES, 5 mM MgCl_2_, pH 7), in a total reaction volume of 20 µL. According to linear regression of the NGD kinetics, 1 U of activity was defined as 0.125 µg ADPRC that, at a concentration of 1.35 µg/mL, converted 60 mM NGD and reached a cGDPr fluorescence plateau after 130-140s (JASCO spectrophotometer, λ_ex_ = 300 nm; λ_em_ = 410 nm, high sensitivity, bandwidth 2 nm) at 25 °C (Winz et al., 2017).

Expression and affinity purification of Npy1 and Npy1(E276Q) was achieved by the following standard procedures with minor changes. Expression in *E. coli*, carrying the corresponding expression vector, was induced at an OD_600_ of ∼0.7) by adding 0.1 mM IPTG. The cells were then chilled for 20 min at 4 °C and incubated at 16 °C, 150 rpm, for an additional 16 h. Cell pellets were subsequently harvested by centrifugation and washed with ice-cold dH_2_O. The pelleted cells were then resuspended in HisTrap Buffer A (50 mM Tris-HCl (pH 7.8), 0.3 M NaCl, 5 mM MgSO_4_, 5 mM 2-mercaptoethanol, 5% glycerol, 5 mM imidazole) and cells lysed by sonification. After centrifugation (37,500 g, 4°C, 30 min) of thus obtained lysates, the supernatant was filtered through a 0.45 µm filter, before loading it on a HisTrap HP 1 mL Column, using an FPLC system (Bio-Rad). The target protein was then eluted by an imidazole gradient, ranging from HisTrap Buffer A to HisTrap Buffer B, which contained an additional 500 mM imidazole. Based on SDS-PAGE analysis, fractions containing the target protein were pooled and concentrated by Amicon Ultra-15 mL Centrifugal Filter Units 10 kDa (Merck) and the HisTrap Buffer B was exchanged with Buffer G ((50 mM Tris-HCl (pH 7.5), 200 mM NaCl, 0.1 mM DTT). Further purification of Npy1 and Npy1(E276Q) by size-exclusion chromatography (SEC) was achieved on a Sephacryl 16/60 S-200 High Resolution column. The final concentration of all proteins was determined employing the Pierce BCA Protein Assay Kit and stored in 50% glycerol at -20 °C.

#### RNA *in vitro* Decapping and NAD Hydrolysis Kinetic Assays

In general, ^32^P-body-labeled NAD-/m^7^G-capped RNAs or ^32^P-labeled NAD were incubated with 0.4 to 1.6 µM recombinant proteins in 40 µL decapping reaction containing 25 mM Tris-HCl (pH 7.5), 50 mM NaCl, 50 mM KCl, 10 mL MgCl_2_, 1 mM DTT, 1 mM MnCl_2_ (was absent in reactions referred to as “without Mn^2+^ kinetics”) and incubated at 37 °C for 120 min. Samples that were treated with the *E. coli* Nudix hydrolase NudC are referred to as ‘positive control’. Aliquots of 5 µL were taken from the reaction mixtures at indicated time points. Thus treated RNAs were mixed with the same volume of 2 X APB Gel Loading Buffer (50 mM Tris-HCl (pH 7.5), 8 M Urea, 20 mM EDTA, 20% Glycerol, 0.01% Xylene Cyanol, 0.01% Bromophenol Blue) and placed on ice for further APB gel electrophoresis. Reaction mixtures, containing NAD, were stored on ice before performing thin-layer chromatography (TLC, DC-Fertigfolie ALUGRAM Xtra SIL G / UV254, 20 cm × 20 cm). Resolution of nucleotides via TLC, at room temperature for 5.5 h, was achieved employing a flow phase of 1 M NH4OAc and ethanol (4:6).

#### Determination of RNA NAD-Capping Ratios in Total RNA and *in vitro* transcribed NAD-/ppp-mRNA Mixtures

For each assay, 100 µg total RNA (gDNA-free) was subjected to ADPRC treatment as fully-treated group. An equal amount of total RNA was applied to the same treatment without ADPRC as background group. The subsequent copper-click reaction, capture by streptavidin beads (streptavidin-unbound flow-through RNAs (non-NAD-RNA) were collected and precipitated with ethanol), as in the standard NAD captureSeq procedure (Winz et al., 2017). For reverse transcription on beads, to each sample of the fully treated group, as well as the background group, were added 2.5 µM random hexamers, 0.5 mM dNTP mix, and dH2O at 65 °C for 5 min. After reaction, the beads were incubated on ice for 2 min. Then, to the reaction was added 1X SSIV Buffer (Thermo Scientific), 5 mM DTT, 10 U/µL SuperScript IV Reverse Transcriptase, and 50 ng/ µL acetylated BSA (Sigma-Aldrich), and the mixture incubated at 23 °C for 10 min, followed by 1h at 55 °C. This was followed by RNA rebinding to streptavidin beads, washing, NaOH-mediated hydrolysis and cDNA precipitation, as described in the NAD captureSeq protocol (Winz et al., 2017). Equivalents of 1 µg of diluted non-NAD-RNAs were reverse transcribed, employing random hexamer oligos, as described above. The relative abundance of transcripts was then quantified by RT-qPCR. The enrichment of transcripts, assessed by RT-qPCR of cDNA, from the fully treated group, the background group, and the non-NAD-cap group was normalized by ribosomal RNA, RDN5-1. The NAD modification ratio of each RNA species was determined by the equation NAD modification ratio = (NAD-RNA_fully-treated group_ - unspecific-binding-RNAbackgroud group) / (NAD-RNA_fully-treated group_ + other-cap-RNA_non-NAD-cap group_).

For the NAD-TDH3 promoter assay, 100 µg total RNA (gDNA-free) were supplemented with 5 ng of NAD-RNAIII (self-prepared) and 5 ng ppp-RNAI (self-prepared) and subjected to the same ADPRC treatment and copper click reaction, as described above. Then, 50 µL of Hydrophilic Streptavidin Magnetic Beads slurry per reaction was utilized to enrich for NAD-modified RNAs in a 96 well plate (flat-bottom, Corning) format. The beads were hereby pre-washed twice with 150 µL Immobilization Buffer (10 mM HEPES (pH 7.2), 1 M NaCl, 5 mM EDTA) and then blocked using acetylated BSA (100 µg/mL, Sigma) in 100 µL Immobilization buffer at room temperature for 10 min under gentle agitation. Subsequently, beads were washed three times with 150 µL Immobilization Buffer. Precipitated RNA from the copper click reaction was dissolved in 100 µL Immobilization Buffer and then added onto the washed beads. The mixture was incubated at room temperature, while shaking at 500 rpm for 1 h. Next, the 96 well plate was placed on a magnetic rack at room temperature for 10 min. Then, the first supernatant was collected. Beads were washed with 100 µL Immobilization Buffer and following that, the second supernatant was collected and pooled with the first one, subsequently referred to as ‘non-NAD-RNAs’. The RNA from the pooled supernatants was precipitated and 1 µg of diluted non-NAD-RNAs was utilized for reverse transcription to determine transcript abundance by RT-qPCR. RNAs captured by magnetic streptavidin beads (+ADPRC for S group, -ADPCR for N group) were washed five times with 150 µL Streptavidin Wash Buffer (8 M Urea, 10 mM Tris-HCl (pH 7.4)) and then washed three times with 150 µL First strand Buffer (25 mM Tris-HCl (pH) 8.3), 37.5 mM KCl, 1.5 mM MgCl_2_). Subsequently, the beads were re-blocked using 100 µL First strand Buffer, containing acetylated BSA (100 µg/mL). Then, equilibrated beads were washed twice with 150 µL 1 X SSIV buffer (Thermo Scientific). Reverse transcription was conducted using random hexamer oligos. The reactions were then transferred into PCR tubes and heated to 65 °C for 5 min to denature the RNA. The tubes were cooled on ice for 5 min, then incubated at 23 °C for 10 min, followed by a heating step at 55 °C for 30 min to enable the reverse transcription reaction by Superscript reverse transcriptase IV. Next, 100 µL NaOH (0.15 M) were added into the reaction and the mixture incubated at 55 °C for 15 min. The first RT supernatant was collected. The remaining beads were repeatedly subjected to the same treatment, by adding 100 µL NaOH (0.15 M). Then the second, residual RT supernatant was pooled with the first one, and is subsequently referred to as “NAD-RNA” (S group) and “background RNA” (N group). Both were ethanol precipitated and directly subjected to RT-qPCR. The abundances of NAD-TDH3 RNA and non-NAD-TDH3 RNA were normalized by transcript abundances of the NAD-RNAIII and ppp-RNAI controls, respectively. The NAD-capping ratio of TDH3 was calculated by dividing NAD-TDH3 by the sum of NAD-TDH3 and non-NAD-TDH3.

For *in vitro* transcribed NAD/ppp-mRNAs, 200 ng of denatured (∼375 fmol) NAD-mRNAs were supplemented with 10 mM Tris-HCl (pH 8.0), 50 mM NaCl, 2 mM fresh DTT, and 1 µM DNAzyme (design procedure as described (Joyce, 2001), sequences are listed in Table S1) in a total volume 9 µL at 65 °C, 5 min. The reaction was cooled to 37 °C. The reactions were then supplemented with 1 µL 500 µM MgCl_2_ and incubated for an additional 1 h at 37 °C. Reactions containing ppp-mRNAs are referred to as ‘positive control’, while reactions containing NAD-mRNA in the absence of DNAzyme was are referred to as negative control. Reactions, containing NAD-mRNA and an additional 2 µM NudC (added after denaturation) served as reference RNA. The reactions were stopped by adding an equal volume of 2X APB Gel Loading Buffer. Separated and cleaved products of NAD-RNA and ppp-/p-RNA were visualized by APB-supplemented PA gel electrophoresis and subsequent SYBR Gold staining. The gel was scanned using a Typhoon imager for further signal quantification. For those samples with weak signal intensities upon DNAzyme treatment on APB gel, RT-qPCR was performed to quantify RNA abundance.

#### Quantitative Reverse Transcription PCR (RT-qPCR) and Standard PCR Procedures

Real-time PCR was performed using 250 nM Fw/Rev primer, 5 µL diluted cDNA, and 1 X SsoAdvanced Universal SYBR Green Supermix in 20 µL reaction. PCR conditions were the following, denaturing step at 95 °C (2 min), 40 cycles of consecutive annealing/extension steps at 95°C for 7s and 60 °C for 15 s, respectively. Melting curves were generated by heating from 65 °C to 95 °C with an incremental increase of 0.5 °C/s. Fluorescence was measured throughout using a LifeCycle 480 Instrument.

Two rounds of PCR were carried out to generate linear DNA templates for *in vitro* translation reactions: The first round aimed to bridge the 5’ UTR/CDS to the firefly Luc2 sequence. The reactions contained 6.4 pM 5’ UTR/CDS template, 6.4 pM bridge region DNA (Table S1), and 6.4 pM firefly Luc2 DNA template, 200 nM dNTPs, 1X Q5 Reaction Buffer (NEB), 0.02 U/µL hot-start Q5 high-fidelity DNA polymerase in a total volume 50 µL. The PCR was initiated by heating to 98 °C for 40 s and followed by 5 cycles (98 °C 10 s, 65 °C 20 s, 72 °C 2 min). The second round of the PCR aimed to specifically amplify the bridged 5’ UTR/CDS, bearing the Luc2 template extension, by additionally adding corresponding 500 nM Fw_5UTR primer, 500 nM Rev_Luc2_polyA primer, and 1 X Q5 Reaction Buffer (NEB) to the final volume of 56.2 µL.

All other PCRs were performed using the Q5 high-fidelity DNA polymerase for amplifying NAD captureSeq cDNA with barcodes, as described (Winz et al., 2017) and related dsDNA templates for *in vitro* translation assay, as linear templates from plasmids. Otherwise, *Taq* polymerase/Q5 high-fidelity DNA polymerase were employed to amplify 5’ UTR/CDS sequences from gDNA and from plasmids, obtained by using standard PCR procedures.

## QUANTIFICATION AND STATISTICAL ANALYSIS

### Quantification

PAGE gel, APB gel, and TLC intensities were quantified using the ImageQuant software (GE Healthcare).

### Statistical analysis

All samples for NAD captureSeq, transcriptomics, UPLC-MS, RT-qPCR, and proteomics were prepared as biological triplicates. The outlined *in vitro* experiments, including NAD kinetics, RNA decapping kinetics, and *in vitro* translations, were performed as technical triplicates. Mean values (n ≥ 3) with standard deviations, Student’s t test (single-tail and double tail (only FCM data), unequal-variance), ANOVA (single factor) were calculated by R/python. For NGS TSS switching relevant math, covariance with two variables, linear trend test, and corresponding p values as well as FDR value were calculated using python scripts. For NGS NAD captureSeq, NAD-RNA enrichment and transcriptome differential expression analysis, DESeq2 was utilized, as described above.

## DATA AND SOFTWARE AVAILABILITY

NGS raw data and analyzed files are available in the GEO repository under the GEO Accession: GSE146368 (For reviewers: https://www.ncbi.nlm.nih.gov/geo/query/acc.cgi?acc=GSE146368, token r:sjkbyimwtlirfqb). Proteomics raw data and analyzed files are deposited in the ProteomeXchange Consortium via the PRIDE repository: PXD017893 (For reviewers: https://www.ebi.ac.uk/pride, Username; Password: ON2B6mbR).

## Supplemental Information

**Table S1:**
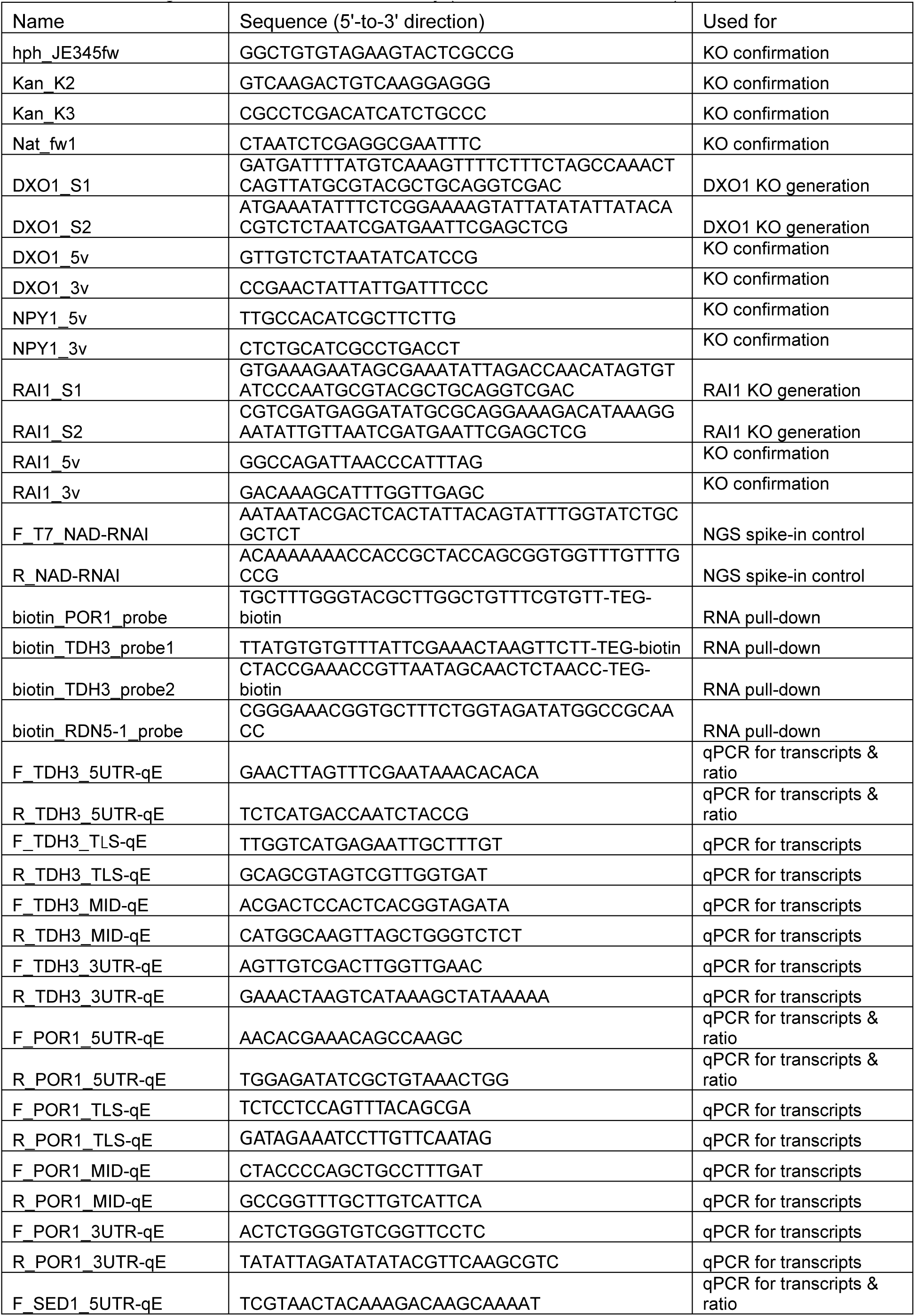
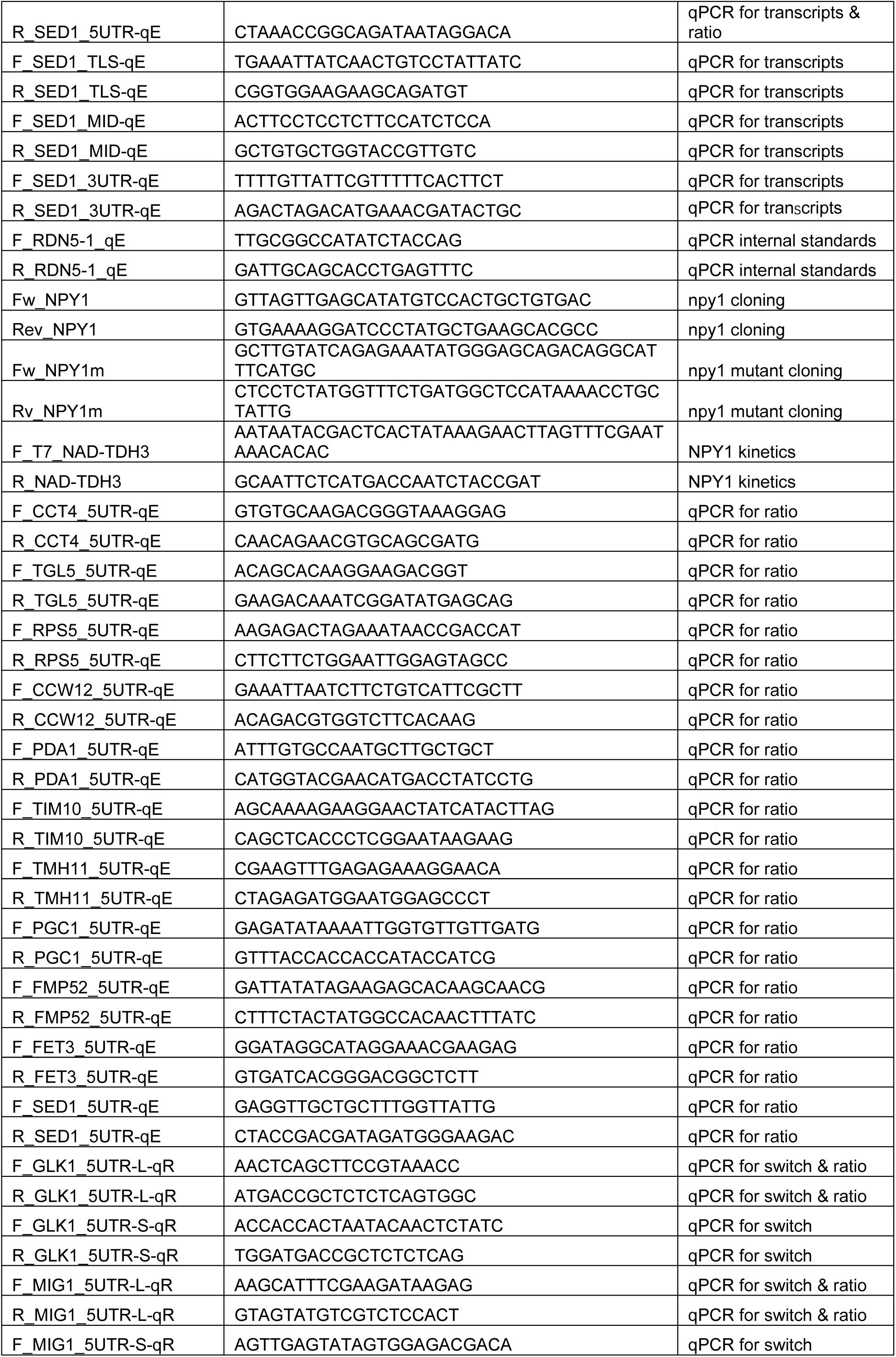
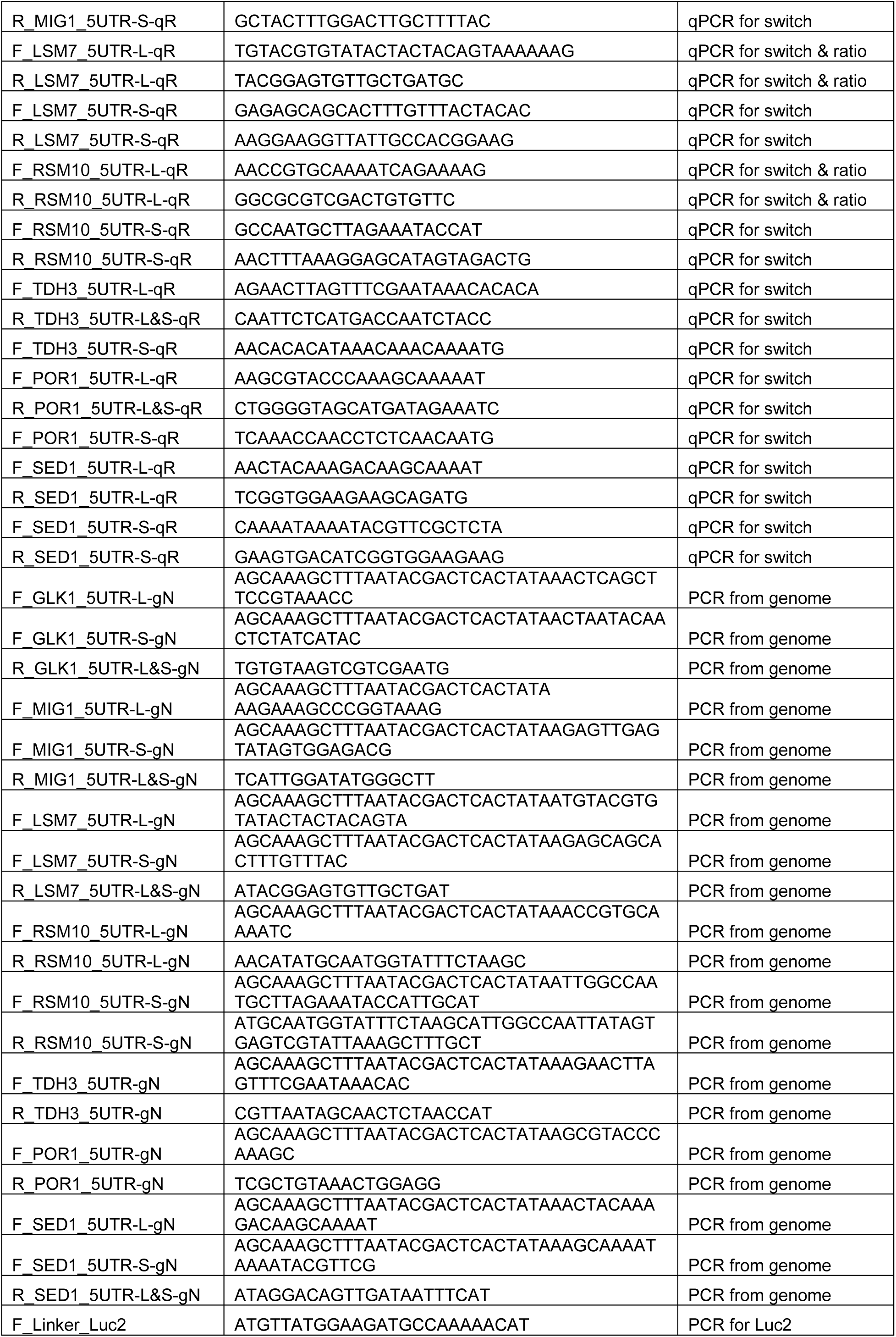
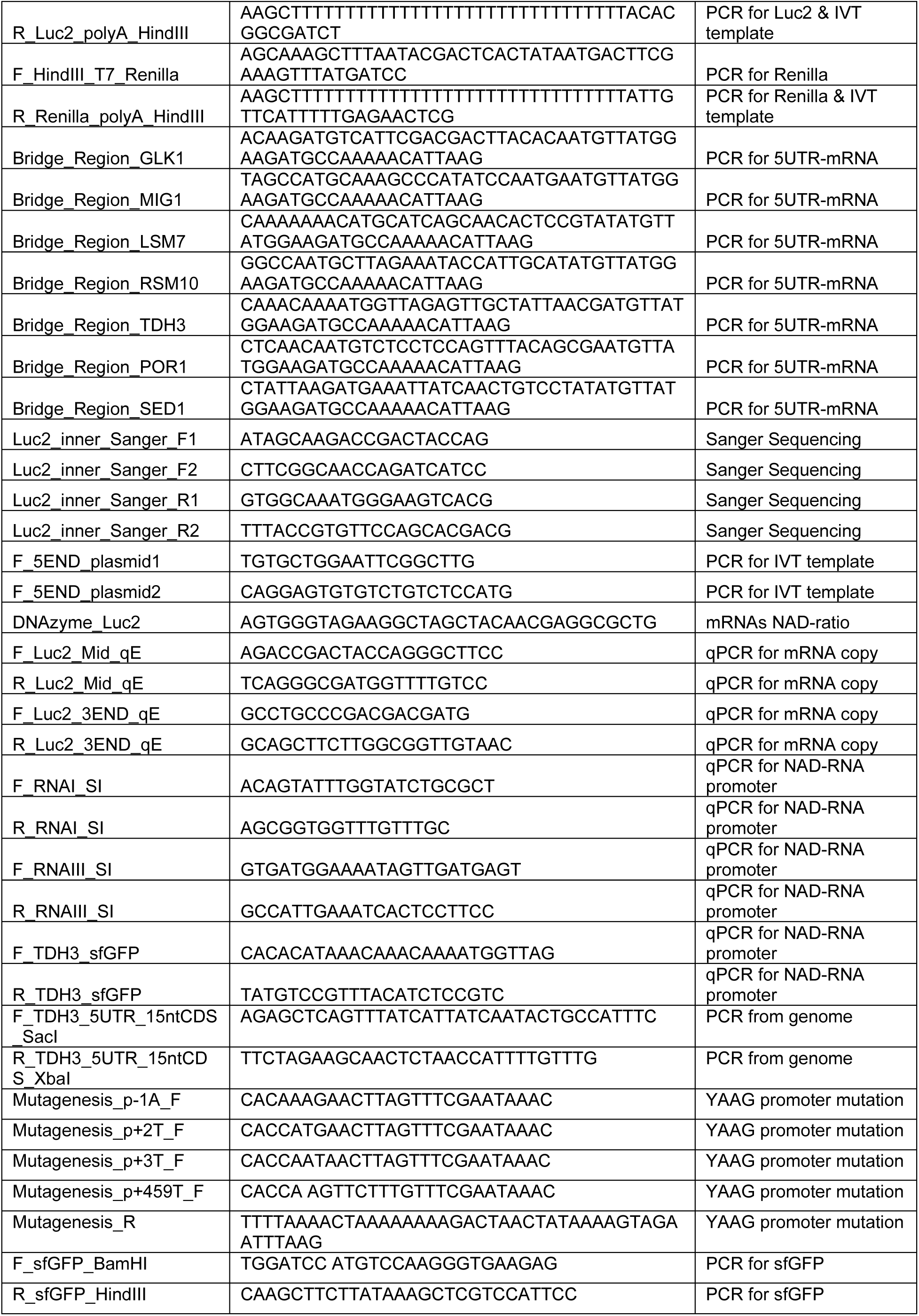
DNA oligonucleotides used in this study (related to STAR methods).

## References

AbdelRaheim, S.R., Cartwright, J.L., Gasmi, L., and McLennan, A.G. (2001). The NADH diphosphatase encoded by the Saccharomyces cerevisiae NPY1 nudix hydrolase gene is located in peroxisomes. Arch. Biochem. Biophys. 388, 18–24.

Baryshnikova, A., Costanzo, M., Dixon, S., Vizeacoumar, F.J., Myers, C.L., Andrews, B., and Boone, C. (2010). Synthetic genetic array (SGA) analysis in Saccharomyces cerevisiae and Schizosaccharomyces pombe. Methods Enzymol. 470, 145–179.

Belasco, J.G. (2010). All things must pass: contrasts and commonalities in eukaryotic and bacterial mRNA decay. Nat. Rev. Mol. Cell Biol. 11, 467–478.

Bird, J.G., Basu, U., Kuster, D., Ramachandran, A., Grudzien-Nogalska, E., Towheed, A., Wallace, D.C., Kiledjian, M., Temiakov, D., Patel, S.S., et al. (2018). Highly efficient 5’ capping of mitochondrial RNA with NAD^+^ and NADH by yeast and human mitochondrial RNA polymerase. Elife 7, e42179.

Bird, J.G., Zhang, Y., Tian, Y., Panova, N., Barvik, I., Greene, L., Liu, M., Buckley, B., Krasny, L., Lee, J.K., et al. (2016). The mechanism of RNA 5’ capping with NAD^+^, NADH and desphospho-CoA. Nature 535, 444–447.

Bresson, S., and Tollervey, D. (2018). Surveillance-ready transcription: nuclear RNA decay as a default fate. Open Biol 8, 170270.

Cahova, H., Winz, M.L., Höfer, K., Nübel, G., and Jäschke, A. (2015). NAD captureSeq indicates NAD as a bacterial cap for a subset of regulatory RNAs. Nature 519, 374–377.

Chang, J.H., Jiao, X., Chiba, K., Oh, C., Martin, C.E., Kiledjian, M., and Tong, L. (2012). Dxo1 is a new type of eukaryotic enzyme with both decapping and 5’-3’ exoribonuclease activity. Nat. Struct. Mol. Biol. 19, 1011–1017.

Frindert, J., Zhang, Y., Nübel, G., Kahloon, M., Kolmar, L., Hotz-Wagenblatt, A., Burhenne, J., Haefeli, W.E., and Jäschke, A. (2018). Identification, Biosynthesis, and Decapping of NAD-Capped RNAs in B. subtilis. Cell Rep. 24, 1890–1901.

Gilbert, W.V., Zhou, K., Butler, T.K., and Doudna, J.A. (2007). Cap-independent translation is required for starvation-induced differentiation in yeast. Science 317, 1224–1227.

Grudzien-Nogalska, E., Bird, J.G., Nickels, B.E., and Kiledjian, M. (2018). “NAD-capQ” detection and quantitation of NAD caps. RNA 24, 1418–1425.

Hahn, S., Hoar, E.T., and Guarente, L. (1985). Each of three “TATA elements” specifies a subset of the transcription initiation sites at the CYC-1 promoter of Saccharomyces cerevisiae. Proc. Natl. Acad. Sci. U. S. A. 82, 8562–8566.

Harlen, K.M., and Churchman, L.S. (2017). Subgenic Pol II interactomes identify region-specific transcription elongation regulators. Mol. Syst. Biol. 13: 900, 1–14.

Höfer, K., Li, S., Abele, F., Frindert, J., Schlotthauer, J., Grawenhoff, J., Du, J., Patel, D.J., and Jäschke, A. (2016). Structure and function of the bacterial decapping enzyme NudC. Nat. Chem. Biol. 12, 730–734.

Jiao, X., Doamekpor, S.K., Bird, J.G., Nickels, B.E., Tong, L., Hart, R.P., and Kiledjian, M. (2017). 5’ End Nicotinamide Adenine Dinucleotide Cap in Human Cells Promotes RNA Decay through DXO-Mediated deNADding. Cell 168, 1015–1027.

Jiao, X., Xiang, S., Oh, C., Martin, C.E., Tong, L., and Kiledjian, M. (2010). Identification of a quality-control mechanism for mRNA 5’-end capping. Nature 467, 608–611.

Kim, M., Krogan, N.J., Vasiljeva, L., Rando, O.J., Nedea, E., Greenblatt, J.F., and Buratowski, S. (2004). The yeast Rat1 exonuclease promotes transcription termination by RNA polymerase II. Nature 432, 517–522.

Lubliner, S., Keren, L., and Segal, E. (2013). Sequence features of yeast and human core promoters that are predictive of maximal promoter activity. Nucleic Acids Res. 41, 5569–5581.

Maicas, E., and Friesen, J.D. (1990). A sequence pattern that occurs at the transcription initiation region of yeast RNA polymerase II promoters. Nucleic Acids Res. 18, 3387–3393.

McMillan, J., Lu, Z., Rodriguez, J.S., Ahn, T.H., and Lin, Z. (2019). YeasTSS: an integrative web database of yeast transcription start sites. Database (Oxford) 2019.

Meijer, H.A., Kong, Y.W., Lu, W.T., Wilczynska, A., Spriggs, R.V., Robinson, S.W., Godfrey, J.D., Willis, A.E., and Bushell, M. (2013). Translational repression and eIF4A2 activity are critical for microRNA-mediated gene regulation. Science 340, 82–85.

Meurer, M., Duan, Y., Sass, E., Kats, I., Herbst, K., Buchmuller, B.C., Dederer, V., Huber, F., Kirrmaier, D., Stefl, M., et al. (2018). Genome-wide C-SWAT library for high-throughput yeast genome tagging. Nat Methods 15, 598–600.

Morales-Filloy, H.G., Zhang, Y., Nübel, G., George, S.E., Korn, N., Wolz, C., and Jäschke, A. (2020). The 5’ NAD Cap of RNAIII Modulates Toxin Production in Staphylococcus aureus Isolates. J. Bacteriol. 202, e00591–00519.

Nagalakshmi, U., Wang, Z., Waern, K., Shou, C., Raha, D., Gerstein, M., and Snyder, M. (2008). The transcriptional landscape of the yeast genome defined by RNA sequencing. Science 320, 1344–1349.

Nübel, G., Sorgenfrei, F.A., and Jäschke, A. (2017). Boronate affinity electrophoresis for the purification and analysis of cofactor-modified RNAs. Methods 117, 14–20.

Rojas-Duran, M.F., and Gilbert, W.V. (2012). Alternative transcription start site selection leads to large differences in translation activity in yeast. RNA 18, 2299–2305.

Shatkin, A.J. (1976). Capping of eucaryotic mRNAs. Cell 9, 645–653.

Tome, J.M., Tippens, N.D., and Lis, J.T. (2018). Single-molecule nascent RNA sequencing identifies regulatory domain architecture at promoters and enhancers. Nat. Genet. 50, 1533–1541.

Topisirovic, I., Svitkin, Y.V., Sonenberg, N., and Shatkin, A.J. (2011). Cap and cap-binding proteins in the control of gene expression. Wiley Interdiscip Rev RNA 2, 277–298.

Vvedenskaya, I.O., Bird, J.G., Zhang, Y., Zhang, Y., Jiao, X., Barvik, I., Krasny, L., Kiledjian, M., Taylor, D.M., Ebright, R.H., et al. (2018). CapZyme-Seq Comprehensively Defines Promoter-Sequence Determinants for RNA 5’ Capping with NAD^+^. Mol. Cell 70, 553–564.

Walters, R.W., Matheny, T., Mizoue, L.S., Rao, B.S., Muhlrad, D., and Parker, R. (2017). Identification of NAD^+^ capped mRNAs in Saccharomyces cerevisiae. Proc. Natl. Acad. Sci. U. S. A. 114, 480–485.

Wang, Y., Li, S., Zhao, Y., You, C., Le, B., Gong, Z., Mo, B., Xia, Y., and Chen, X. (2019). NAD^+^-capped RNAs are widespread in the Arabidopsis transcriptome and can probably be translated. Proc. Natl. Acad. Sci. U. S. A. 116, 12094–12102.

Winz, M.L., Cahova, H., Nübel, G., Frindert, J., Höfer, K., and Jäschke, A. (2017). Capture and sequencing of NAD-capped RNA sequences with NAD captureSeq. Nat. Protoc. 12, 122–149.

Xu, W., Dunn, C.A., and Bessman, M.J. (2000). Cloning and characterization of the NADH pyrophosphatases from Caenorhabditis elegans and Saccharomyces cerevisiae, members of a Nudix hydrolase subfamily. Biochem. Biophys. Res. Commun. 273, 753–758.

Zhang, D., Liu, Y., Wang, Q., Guan, Z., Wang, J., Liu, J., Zou, T., and Yin, P. (2016). Structural basis of prokaryotic NAD-RNA decapping by NudC. Cell Res. 26, 1062–1066.

Zhang, H., Zhong, H., Zhang, S., Shao, X., Ni, M., Cai, Z., Chen, X., and Xia, Y. (2019). NAD tagSeq reveals that NAD^+^-capped RNAs are mostly produced from a large number of protein-coding genes in Arabidopsis. Proc. Natl. Acad. Sci. U. S. A. 116, 12072–12077.

Zhang, Z., and Dietrich, F.S. (2005). Mapping of transcription start sites in Saccharomyces cerevisiae using 5’ SAGE. Nucleic Acids Res. 33, 2838–2851.

## Methods References

Agresti, A. (2014). Categorical Data Analysis (Hoboken: Wiley).

Aibar, S., Fontanillo, C., Droste, C., and De Las Rivas, J. (2015). Functional Gene Networks: R/Bioc package to generate and analyse gene networks derived from functional enrichment and clustering. Bioinformatics 31, 1686–1688.

Anders, S., Pyl, P.T., and Huber, W. (2015). HTSeq--a Python framework to work with high-throughput sequencing data. Bioinformatics 31, 166–169.

Bailey, T.L., Boden, M., Buske, F.A., Frith, M., Grant, C.E., Clementi, L., Ren, J., Li, W.W., and Noble, W.S. (2009). MEME SUITE: tools for motif discovery and searching. Nucleic Acids Res. 37, W202–208.

Brachmann, C.B., Davies, A., Cost, G.J., Caputo, E., Li, J., Hieter, P., and Boeke, J.D. (1998). Designer deletion strains derived from Saccharomyces cerevisiae S288C: a useful set of strains and plasmids for PCR-mediated gene disruption and other applications. Yeast 14, 115–132.

Collart, M.A., and Oliviero, S. (2001). Preparation of yeast RNA. Curr. Protoc. Mol. Biol. Chapter 13, Unit13 12.

Crooks, G.E., Hon, G., Chandonia, J.M., and Brenner, S.E. (2004). WebLogo: a sequence logo generator. Genome Res. 14, 1188–1190.

Fischl, W., and Bartenschlager, R. (2013). High-throughput screening using dengue virus reporter genomes. Methods Mol. Biol. 1030, 205–219.

Franken, H., Mathieson, T., Childs, D., Sweetman, G.M., Werner, T., Togel, I., Doce, C., Gade, S., Bantscheff, M., Drewes, G., et al. (2015). Thermal proteome profiling for unbiased identification of direct and indirect drug targets using multiplexed quantitative mass spectrometry. Nat. Protoc. 10, 1567–1593.

Freese, N.H., Norris, D.C., and Loraine, A.E. (2016). Integrated genome browser: visual analytics platform for genomics. Bioinformatics 32, 2089–2095.

Giaever, G., Chu, A.M., Ni, L., Connelly, C., Riles, L., Veronneau, S., Dow, S., Lucau-Danila, A., Anderson, K., Andre, B., et al. (2002). Functional profiling of the Saccharomyces cerevisiae genome. Nature 418, 387–391.

Harju, S., Fedosyuk, H., and Peterson, K.R. (2004). Rapid isolation of yeast genomic DNA: Bust n’ Grab. BMC Biotechnol. 4, 8.

Huang da, W., Sherman, B.T., and Lempicki, R.A. (2009). Systematic and integrative analysis of large gene lists using DAVID bioinformatics resources. Nat. Protoc. 4, 44–57.

Huber, W., von Heydebreck, A., Sultmann, H., Poustka, A., and Vingron, M. (2002). Variance stabilization applied to microarray data calibration and to the quantification of differential expression. Bioinformatics 18 Suppl 1, S96–104.

Hughes, C.S., Foehr, S., Garfield, D.A., Furlong, E.E., Steinmetz, L.M., and Krijgsveld, J. (2014). Ultrasensitive proteome analysis using paramagnetic bead technology. Mol. Syst. Biol. 10, 757.

Janke, C., Magiera, M.M., Rathfelder, N., Taxis, C., Reber, S., Maekawa, H., Moreno-Borchart, A., Doenges, G., Schwob, E., Schiebel, E., et al. (2004). A versatile toolbox for PCR-based tagging of yeast genes: new fluorescent proteins, more markers and promoter substitution cassettes. Yeast 21, 947–962.

Joyce, G.F. (2001). RNA cleavage by the 10-23 DNA enzyme. Methods in enzymology 341, 503–517.

Khmelinskii, A., and Knop, M. (2014). Analysis of protein dynamics with tandem fluorescent protein timers. Methods Mol. Biol. 1174, 195–210.

Knop, M., and Schiebel, E. (1998). Receptors determine the cellular localization of a gamma-tubulin complex and thereby the site of microtubule formation. EMBO J. 17, 3952–3967.

Knop, M., Siegers, K., Pereira, G., Zachariae, W., Winsor, B., Nasmyth, K., and Schiebel, E. (1999). Epitope tagging of yeast genes using a PCR-based strategy: More tags and improved practical routines. Yeast 15, 963–972.

Langmead, B., Trapnell, C., Pop, M., and Salzberg, S.L. (2009). Ultrafast and memory-efficient alignment of short DNA sequences to the human genome. Genome Biol. 10, R25.

Li, H., and Durbin, R. (2009). Fast and accurate short read alignment with Burrows-Wheeler transform. Bioinformatics 25, 1754–1760.

Li, H., Handsaker, B., Wysoker, A., Fennell, T., Ruan, J., Homer, N., Marth, G., Abecasis, G., Durbin, R., and Genome Project Data Processing, S. (2009). The Sequence Alignment/Map format and SAMtools. Bioinformatics 25, 2078–2079.

Love, M.I., Huber, W., and Anders, S. (2014). Moderated estimation of fold change and dispersion for RNA-seq data with DESeq2. Genome Biol. 15, 550.

Mumberg, D., Muller, R., and Funk, M. (1994). Regulatable promoters of Saccharomyces cerevisiae: comparison of transcriptional activity and their use for heterologous expression. Nucleic Acids Res. 22, 5767–5768.

Mumberg, D., Muller, R., and Funk, M. (1995). Yeast vectors for the controlled expression of heterologous proteins in different genetic backgrounds. Gene 156, 119–122.

Munshi, C., and Lee, H.C. (1997). High-level expression of recombinant Aplysia ADP-ribosyl cyclase in offhia pastoris by fermentation. Protein Expr. Purif. 11, 104–110.

Perkins, D.N., Pappin, D.J.C., Creasy, D.M., and Cottrell, J.S. (1999). Probability-based protein identification by searching sequence databases using mass spectrometry data. Electrophoresis 20, 3551–3567.

Reichel, M., Liao, Y., Rettel, M., Ragan, C., Evers, M., Alleaume, A.M., Horos, R., Hentze, M.W., Preiss, T., and Millar, A.A. (2016). In Planta Determination of the mRNA-Binding Proteome of Arabidopsis Etiolated Seedlings. Plant Cell 28, 2435–2452.

Ritchie, M.E., Phipson, B., Wu, D., Hu, Y., Law, C.W., Shi, W., and Smyth, G.K. (2015). limma powers differential expression analyses for RNA-sequencing and microarray studies. Nucleic Acids Res. 43, e47.

Schiestl, R.H., and Gietz, R.D. (1989). High efficiency transformation of intact yeast cells using single stranded nucleic acids as a carrier. Curr. Genet. 16, 339–346.

Shannon, P., Markiel, A., Ozier, O., Baliga, N.S., Wang, J.T., Ramage, D., Amin, N., Schwikowski, B., and Ideker, T. (2003). Cytoscape: a software environment for integrated models of biomolecular interaction networks. Genome Res. 13, 2498–2504.

Sikorski, R.S., and Hieter, P. (1989). A system of shuttle vectors and yeast host strains designed for efficient manipulation of DNA in Saccharomyces cerevisiae. Genetics 122, 19–27.

Tripathi, S., Pohl, M.O., Zhou, Y., Rodriguez-Frandsen, A., Wang, G., Stein, D.A., Moulton, H.M., DeJesus, P., Che, J., Mulder, L.C., et al. (2015). Meta- and Orthogonal Integration of Influenza “OMICs” Data Defines a Role for UBR4 in Virus Budding. Cell Host Microbe 18, 723–735.

Werner, T., Sweetman, G., Savitski, M.F., Mathieson, T., Bantscheff, M., and Savitski, M.M. (2014). Ion coalescence of neutron encoded TMT 10-plex reporter ions. Anal. Chem. 86, 3594–3601.

Wu, C., and Sachs, M.S. (2014). Preparation of a Saccharomyces cerevisiae cell-free extract for in vitro translation. Methods Enzymol. 539, 17–28.

